# Cell-type label errors inflate neighborhood effects and can reverse niche claims in spatial transcriptomics

**DOI:** 10.64898/2026.11.23.768633

**Authors:** Vlad Elgart, Joseph Loscalzo

## Abstract

Niche analyses ask how a cell’s neighbors shape its expression beyond its type. The type comes from a label, and labels are imperfect. Here we show that label errors inflate neighborhood effects and can reverse niche claims. A mislabeled cell still expresses its true type, and because cells of a type sit near one another, the neighborhood is credited with what the label misses. Standard practice, which calls labels on the modeled counts and selects cells by them, hid, exaggerated, or reversed this inflation. Labeling one half of each cell’s counts and modeling the other revealed it in nearly every population of six datasets. On semi-synthetic colon data, a labeler that over-called one fibroblast subtype reversed claims of the niche method Niche-DE; new draws of the counts and errors spread at random over subtypes almost never did. Our software, nichepeek, reports the direction of this bias in a user’s data.

## 1 Introduction

Spatial transcriptomics lets us ask how a cell’s neighbors shape its expression. Niche methods answer by comparing cells of the same type that sit in different surroundings [1–4], and their claims guide how we read tissues in health and disease. Every such comparison takes the cell’s type, and the types of its neighbors, from labels.

Labels are imperfect. They come from clustering or classifying sparse profiles, and in imaging-based data, many of them change when the counts are resampled (Supplementary Table S8). Statistics tells us what to expect. Cell type confounds neighborhood effects, because cells of a type sit together and express alike. When a confounder is measured with error, adjusting for it leaves residual confounding [5], which is credited to correlated variables [6]. A mislabeled cell should, therefore, pass part of its type’s effect to its neighborhood. Existing corrections address the assignment of transcripts to cells [7–10] and mostly take the labels as given. Tools that aggregate label probabilities over cell types do so on the stated premise that working at the level of cell types avoids noise in single cells [11]; aggregation dilutes errors spread at random, but not errors that concentrate on one subtype. Robustness checks, where reported, often perturb labels at random [12–14].

Standard practice also hides the problem. Analysts call labels on the same counts to which they then fit the niche model, a double use of data that distorts inference after clustering [15–17]. They also select the cells under study by the same labels. Both steps change the estimate in ways that a reader cannot see.

Here, we show that label errors inflate neighborhood effects and can reverse niche claims. We call labels on one half of each cell’s counts, fit the model on the other half, and keep the studied population fixed. With this protocol, errors in a cell’s own label inflated the neighborhood effect in simulations with known truth and in nearly every population of six datasets, and standard practice hid, exaggerated, or reversed that inflation. Labels were also less stable in inflamed colon than in controls, and the estimated bias was larger there. On semi-synthetic colon data, a labeler that over-called one fibroblast subtype reversed claims of the niche method Niche-DE that true labels supported. Errors spread at random over subtypes reversed none. Where we could test it, keeping only claims that a fit with a second annotation also made removed nearly all reversals, at the cost of more than a third of the robust claims of a single fit. Part of a niche claim can, therefore, come from the labels, and a check of it needs errors of the kind that labelers make. We release the protocol as software, nichepeek (Supplementary Fig. S3).

## 2 Results

### 2.1 A label enters a neighborhood analysis twice

We measure the neighborhood effect on a population of target cells, such as the fibroblasts of a tissue section (Fig. 1a; Methods). We describe each target by the label composition of its nearest neighbors and group the targets by these compositions. The groups explain a share of the variance in the targets’ expression, and we divide this share in two by permutation. *Identity*, *I*, is the part that remains when expression is shuffled among targets with the same label; the targets’ own labels account for it. The *neighborhood share*, *V*, is the rest: what the neighborhood explains among cells with the same label. *V* is an association with position rather than a mechanism. It is our measure of the neighborhood effect that niche analyses estimate.

**Figure 1:**
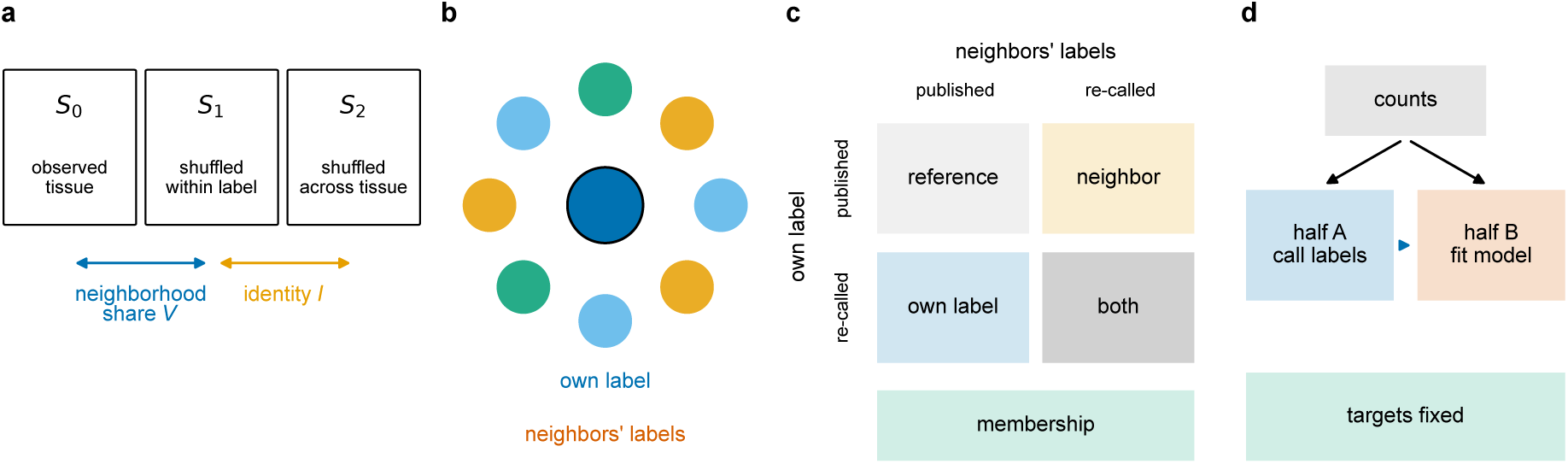
A label enters a neighborhood analysis twice. **a**, The measure. Neighborhood groups explain a share of the variance of target-cell expression in the observed tissue (*S*_0_), after expression is shuffled among targets with the same label (*S*_1_), and after it is shuffled across the tissue (*S*_2_). The neighborhood share *V* and identity *I* are the differences. **b**, The two roles. The own label of a target cell (center) sets the classes within which expression is shuffled; the labels of its neighbors define the composition by which targets are grouped. **c**, The channels. Published or re-called labels in the own role and in the neighbor role give the reference, the own-label channel (*T*), the neighbor channel (*N*), and both together; the membership channel (*M*) is the further effect of selecting the targets by the re-called labels. **d**, The protocol. Labels are called on one binomial half of each cell’s counts, the model is fitted on the other half, and the targets are fixed by the published labels.

A label plays two roles in this estimate (Fig. 1b). As the cell’s own label, it sets the classes within which we shuffle expression. As a neighbor’s label, it enters the compositions that group the targets. Label error can, therefore, act through three channels (Fig. 1c). The *own-label* channel carries errors in the target cell’s own label. It cannot change the variance that the groups explain. Any change in what the cells’ own labels explain, therefore, appears with the opposite sign in the neighborhood share. Its sign and its size are empirical. The *neighbor*channel carries errors in the labels of the cell’s neighbors. The *membership* channel appears when the labels also select which cells are targets.

Standard practice mixes these channels because it calls labels on the counts that it then models and selects the targets by those labels. Our protocol separates them (Fig. 1d). We split each cell’s counts into two independent halves, call labels on one half, and score the model on the other. We keep the target population fixed by the published labels. We then re-call the labels in one role at a time with a nearest-centroid classifier, and we add label error by thinning the counts. We call the response of the neighborhood share to this added error the label effect.

### 2.2 In simulations, own-label error inflates the neighborhood share, and standard practice hides it

We first applied the protocol to simulated tissues with known cell types and a planted neighborhood signal (Methods). Label error inflated the neighborhood share (Fig. 2a). In tissues with no planted signal, it created one, in 33 of 36 sections. A mislabeled cell carries the expression of its true type, and true types are spatially structured. With the channels separated, almost all of the bias came through the cells’ own labels (Fig. 2b).

**Figure 2:**
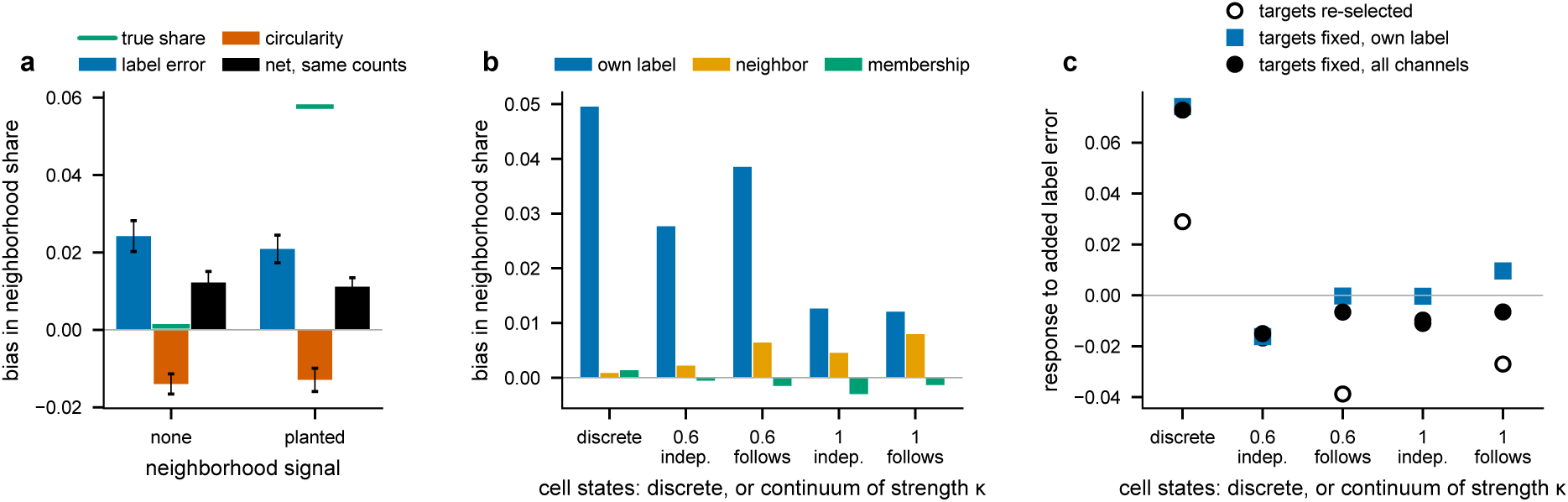
Simulations with known truth. **a**, Bias in the neighborhood share from label error (labels from one half of the counts, model on the other), from circularity (labels scored on the half from which they were called), and net (the same-count analysis), in tissues without and with a planted neighborhood signal. Bars, mean *±* s.e. over sections; green ticks, true neighborhood share. **b**, Native bias by channel with the target population fixed, for discrete cell types and for cell states that vary along the axis between types with strength *κ*, independently of the neighborhood (indep.) or following it (follows). **c**, Response of the label effect to added label error (slope over the thinning levels) in the same regimes, with targets re-selected by the re-called labels and with fixed targets. All panels use the multinomial noise correction applied to the real data.

We expected the inflation, but the estimate does not force it. Errors that are independent of the neighborhood can only move variance from identity to the neighborhood share (Supplementary Note 1). Real errors, however, land on the label typical of the cell’s neighborhood more often than chance (Supplementary Note 2), which offsets the inflation. In none of our simulations did the offset prevail.

Both features of standard practice distorted this result. Scoring labels on the counts from which they were called lowered the neighborhood share, such that a same-count analysis saw only part of the inflation (Fig. 2a; Supplementary Fig. S6a). With targets selected by the recalled labels, the response to added error had the opposite sign to the bias wherever cell states formed a continuum (Fig. 2c). Even with fixed targets, the response can vanish or turn negative for continuous states, although label error inflated the neighborhood share in every regime. A positive own-label response is, therefore, evidence of inflation, whereas a null or negative response is not evidence of accurate labels.

### 2.3 The own-label channel inflated the neighborhood share in real tissue, and the standard design concealed it

We applied the protocol to seven cell populations: mouse colon stroma (Xenium [18]), mouse colon fibroblasts during colitis (MERFISH [19]), hepatocytes of healthy and of fibrotic human liver (MERFISH [20]), and fibroblasts, myeloid cells, and endothelial cells of rheumatoid arthritis synovium (Xenium [21]). The own-label channel inflated the neighborhood share in every one of these populations whose targets had identity to transfer (Fig. 3a; Supplementary Table S1), and within populations, it grew with identity (Fig. 3b). The neighbor channel was negative and usually small. The own-label channel had the same sign in two further datasets: kidney samples from the same patients imaged on two platforms [22], and lung cancer with whole lineages as target sets [10] (Supplementary Note 2; Supplementary Figs. S4 and S5). It was close to zero where the labels explained almost none of the expression: in fibrotic liver and on the largest lung panel.

**Figure 3:**
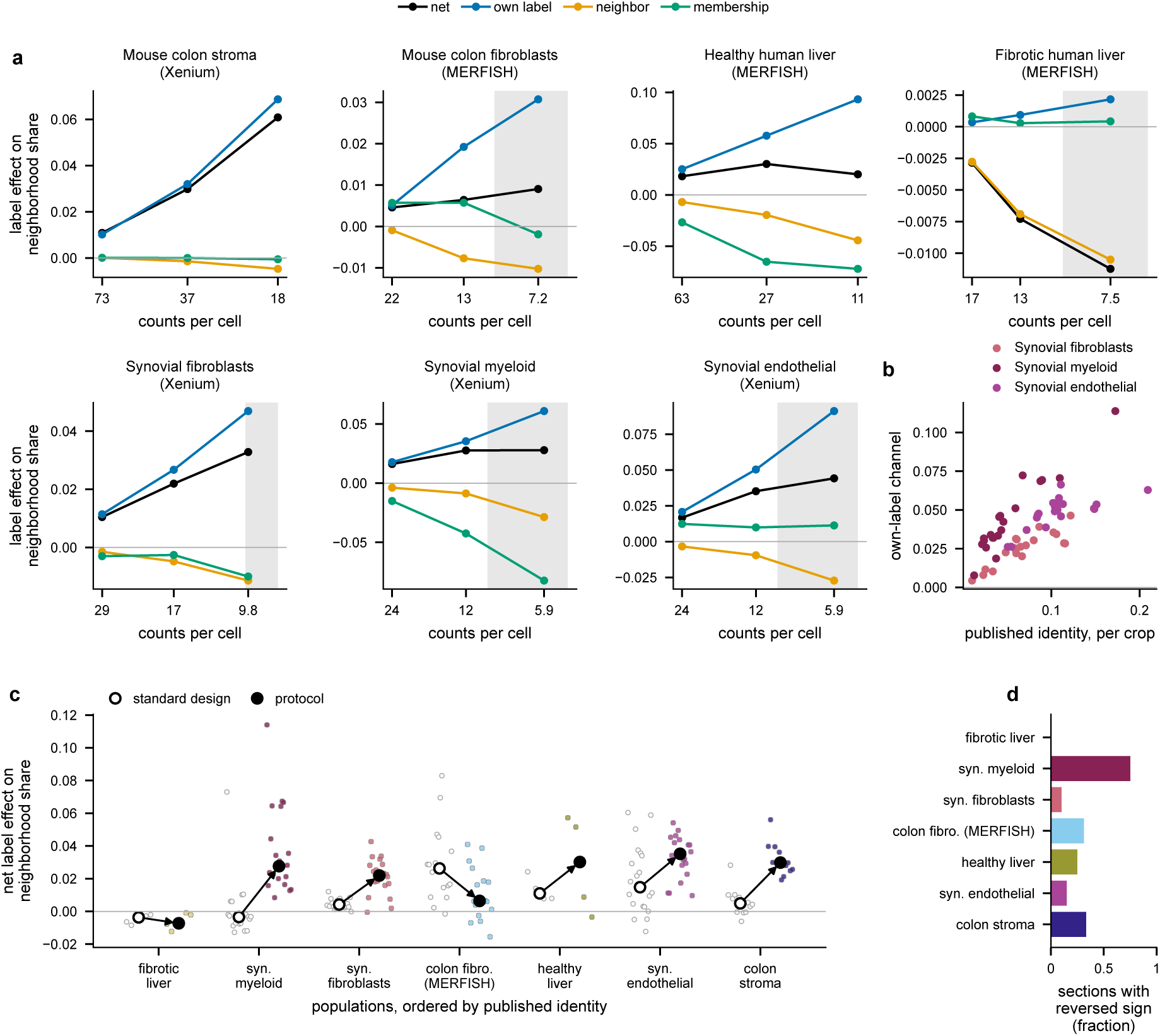
Label error acts through the same channels in every population, and the standard design conceals it. **a**, Label effect on the neighborhood share by channel against the depth that the model sees after splitting and thinning; medians over sections. Count splitting and fixed targets throughout. Gray band, thinning levels that leave fewer than ten counts per cell, which are shown but not interpreted; in four populations, the middle level lies just above this depth. **b**, Own-label channel against published identity per crop at the middle thinning level, for the three synovial populations on identical crops. **c**, Net label effect at the middle thinning level with the standard design (labels and model on the same counts, targets re-selected by the labels; open) and with the protocol (filled). Small symbols, sections (crops in synovium); large symbols, medians. **d**, Fraction of sections or crops whose label effect changes sign between the two designs. Fibro., fibroblasts; syn., synovial (rheumatoid arthritis synovium).

The membership channel had no fixed sign. In healthy liver and in synovial myeloid cells, whose targets are easily confused with other cells, it was negative and larger than the own-label channel. Any analysis that selects its targets by label is exposed to this channel because different labels select different cells. A large membership channel means that a result describes the labeled population rather than the cell type.

Run as a standard pipeline would run it, with labels and model on the same counts and targets selected by the labels, the same diagnostic gave a different answer in most populations (Fig. 3c,d). In colon stroma, the label effect appeared about 6-fold smaller than with the protocol. In synovial myeloid cells, its sign reversed in 15 of 20 crops. In MERFISH colon, where the targets are a small minority of cells, it was overstated. A small label effect in a standard analysis is, therefore, not evidence of accurate labels.

A second estimator showed the same channels. In a niche regression of the NCEM type [3], the own-label channel inflated the niche effect and the neighbor channel deflated it in all seven populations, and the net sign depended on the dataset (Supplementary Note 2).

### 2.4 The estimate has the sign of a known bias, but not its size

Real data have no true labels, and we, therefore, built data that have them (Fig. 4a; Methods). We divided each cell’s counts into two shares. Labels called on the first share served as the truth. The second share served as the experiment: we labeled it again on its own counts and ran nichepeek on it unchanged. The true labels, scored on the same counts, gave the bias directly.

**Figure 4:**
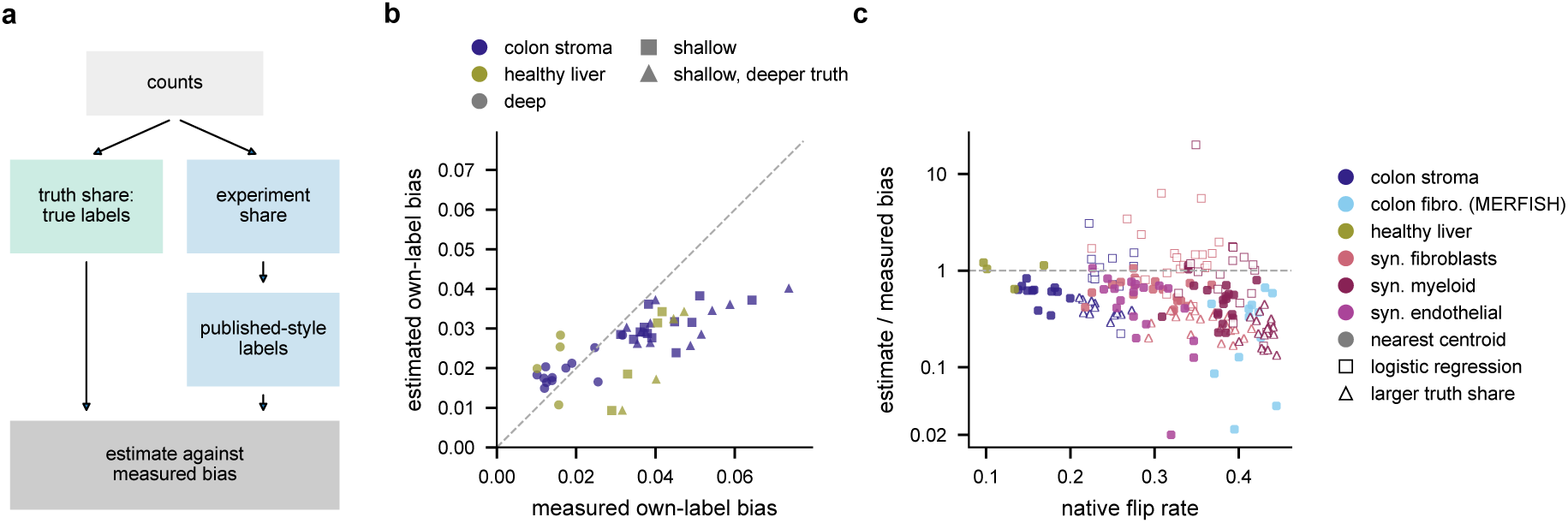
The estimate has the sign of a known bias, but not its size. **a**, Semi-synthetic data. Each cell’s counts are divided into a truth share, whose labels serve as the truth, and an experiment share, which is labeled on its own counts to give published-style labels. nichepeek runs on the experiment share, and further arms score the true labels on the same half and split. **b**, Estimated against measured own-label bias per mouse (colon stroma) and per crop (healthy liver; two crops share one donor), with truth and experiment shares of equal depth (deep), with a thinned experiment share (shallow), and with a thinned experiment share and a larger truth share (shallow, deeper truth); first thinning step *p* = 0.5. Dashed, equality. **c**, Estimate (first thinning step *p* = 0.8) over measured own-label bias against the native flip rate, per section or crop, on semi-synthetic data built from six populations. The re-calling labeler, trained on the synthetic labels, is the nearest-centroid labeler (filled circles), logistic regression (squares), or the nearest-centroid labeler with a larger truth share (triangles). Dashed, equality. The 7 sections with an estimate of zero or below cannot be shown on the log scale; 3 values outside 0.02 to 20, in this panel and in Supplementary Fig. S7c, are drawn at those limits. Panels b and c are measured against the truth labels as called; Supplementary Note 3 corrects for their own errors. Fibro., fibroblasts; syn., synovial.

In colon stroma and healthy liver, the measured own-label bias was positive, and the nichepeek estimate had its sign, in every one of the 15 independent units, at every depth, and with a second labeler (Fig. 4b). The bias was expected to be positive here because the errors in these data are counting noise from the labeler itself (Supplementary Note 1). The test, therefore, shows that the estimate finds a bias of this kind. It does not cover errors that follow the neighborhood.

The size was less reliable (Supplementary Fig. S7a,b). On semi-synthetic data built from six populations, the uncalibrated estimate fell short of the measured bias with our default labeler, by more where labels were less stable, but not consistently with a second labeler (Fig. 4c). Misclassification SIMEX [23] did no better (Supplementary Fig. S7c). A size, therefore, requires the semi-synthetic check, run with the labeler actually used, on the user’s own data (Supplementary Note 3).

The check applies to the labels that the protocol calls. Published labels are called and scored on all of the counts, and semi-synthetic truth from the same classifier cannot measure their bias (Supplementary Table S12). For published labels, the direction rests on simulations with exact truth. There, such labels inflated the neighborhood share where subtypes were confusable and not where they separated well, and the bias reversed where cell states formed a continuum that followed the neighborhood (Supplementary Table S11; Supplementary Note 3).

### 2.5 Label error differed between conditions and between neighborhoods

Niche analyses often compare conditions, and label stability can differ between them. In mouse colon stroma, labels were less stable after colitis induced by dextran sulfate sodium (DSS) than in water-treated mice, whereas the response of the neighborhood share per added label change differed less (Fig. 5a). The estimated own-label bias is the product of the two, and it was larger in every DSS-treated mouse than in any water-treated one (Supplementary Note 2). The difference, though not its complete separation, held with labelers whose class priors match the published label frequencies (Supplementary Table S14). The contrast in the neighborhood share itself was not resolved between the two groups of mice (Supplementary Table S3), such that we do not say how much of it the labels explain. A contrast in neighborhood effects between conditions should, therefore, be reported together with the contrast in label stability.

**Figure 5:**
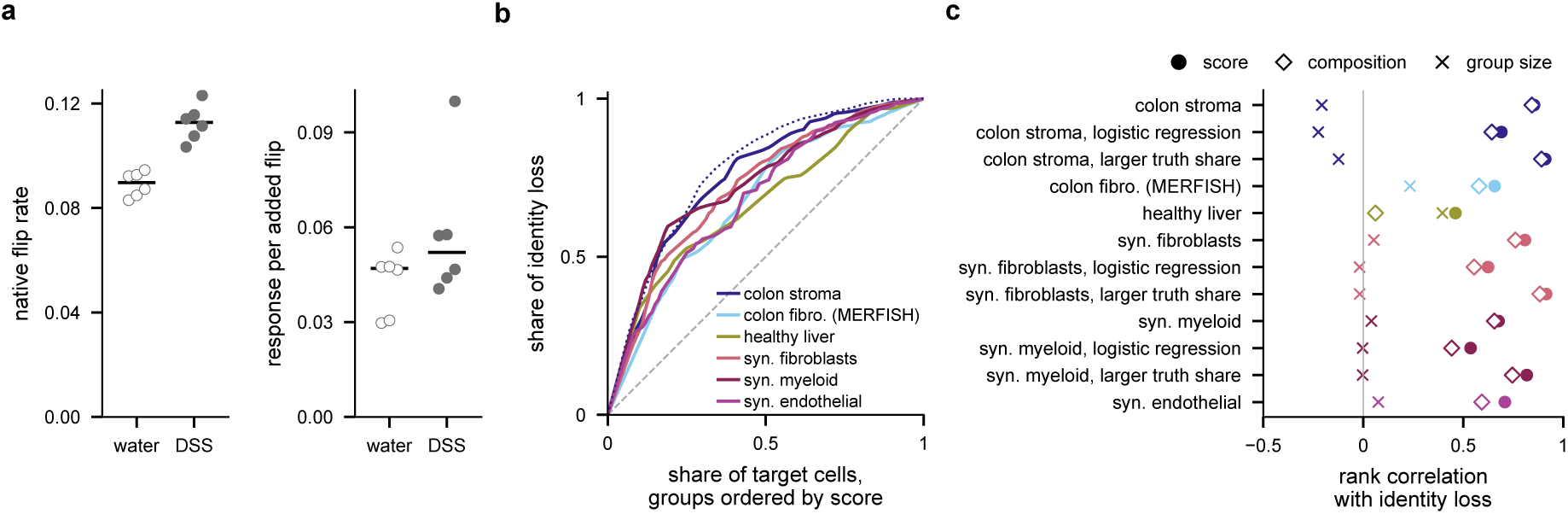
Label error differed between conditions and between neighborhoods. **a**, The two factors of the estimated own-label bias, per mouse in colon stroma after water or after colitis induced by dextran sulfate sodium (DSS). Left, native flip rate, the share of target labels that change when the counts are resampled. Right, response of the neighborhood share per added flip, with the small first thinning step of Fig. 4c. Bars, medians. **b**, Share of the identity lost to label error against the share of targets, with neighborhood groups ordered by the score computed from the re-called labels. Median curve over sections per population (nearest-centroid labeler). Dotted, colon stroma with groups ordered by the loss itself. Dashed, no ranking. **c**, Per run, the median over sections of the Spearman correlation over the 12 neighborhood groups between the identity loss and the score, the imbalance of label composition alone, or group size. The score weights a group’s imbalance between pairs of re-called labels by how different and how often confused the two labels are (Methods). Fibro., fibroblasts; syn., synovial.

Label error also acted unevenly within a tissue. It removed the most identity from neighborhood groups with an uneven mix of confusable subtypes (Fig. 5b). The imbalance of label composition, computed from the labels alone, ranked the groups by their loss in every population but liver (Fig. 5c). A claim about a specific niche is most exposed when it rests on such groups.

### 2.6 Over-calling one subtype reversed niche claims of Niche-DE

We then asked whether label error changes the calls of a widely used niche method. Niche-DE tests, for each gene, whether the expression of an index type depends on the abundance of each neighboring niche type. It reports a call with a sign for each gene, index type, and niche type [4]. We ran it on the semi-synthetic colon data of Section 2.4, on all twelve sections and on three independent splits of the counts (Methods). True labels came from the truth share. The model saw one half of the experiment share, a quarter of the native depth. Labels re-called on the other half entered as the cell’s own label, as the neighbors’ labels, or as both.

At this depth, single fits were not reproducible, even with true labels. Refitting on the other half of the counts changed more than half of the calls, and label error changed a similar share (Supplementary Fig. S8). Effect sizes and signs were stable. Reversals separated the two errors. New draws of the counts and a new seed almost never reversed a call of the true-label fit. We call a claim robust when true labels supported it on both halves of the counts, with a second random seed, and on a second split. Label error reversed 75 robust claims, and most reversals recurred on a further independent split (Fig. 6a,b). Most occurred in one inflamed section, and most occurred with error in the neighbors’ labels alone. Labels re-called on the half that the model saw reversed about as many claims and made about as many false claims, such that splitting the counts mattered little for these calls (Supplementary Note 4). Label error and new draws of the counts also made claims that the true-label fits of the model’s half never made. Label error made more of them, and more of its claims recurred. About half of the reversals involved epithelial genes in fibroblasts, an association that transcripts donated by neighbors create even with correct labels [24].

**Figure 6:**
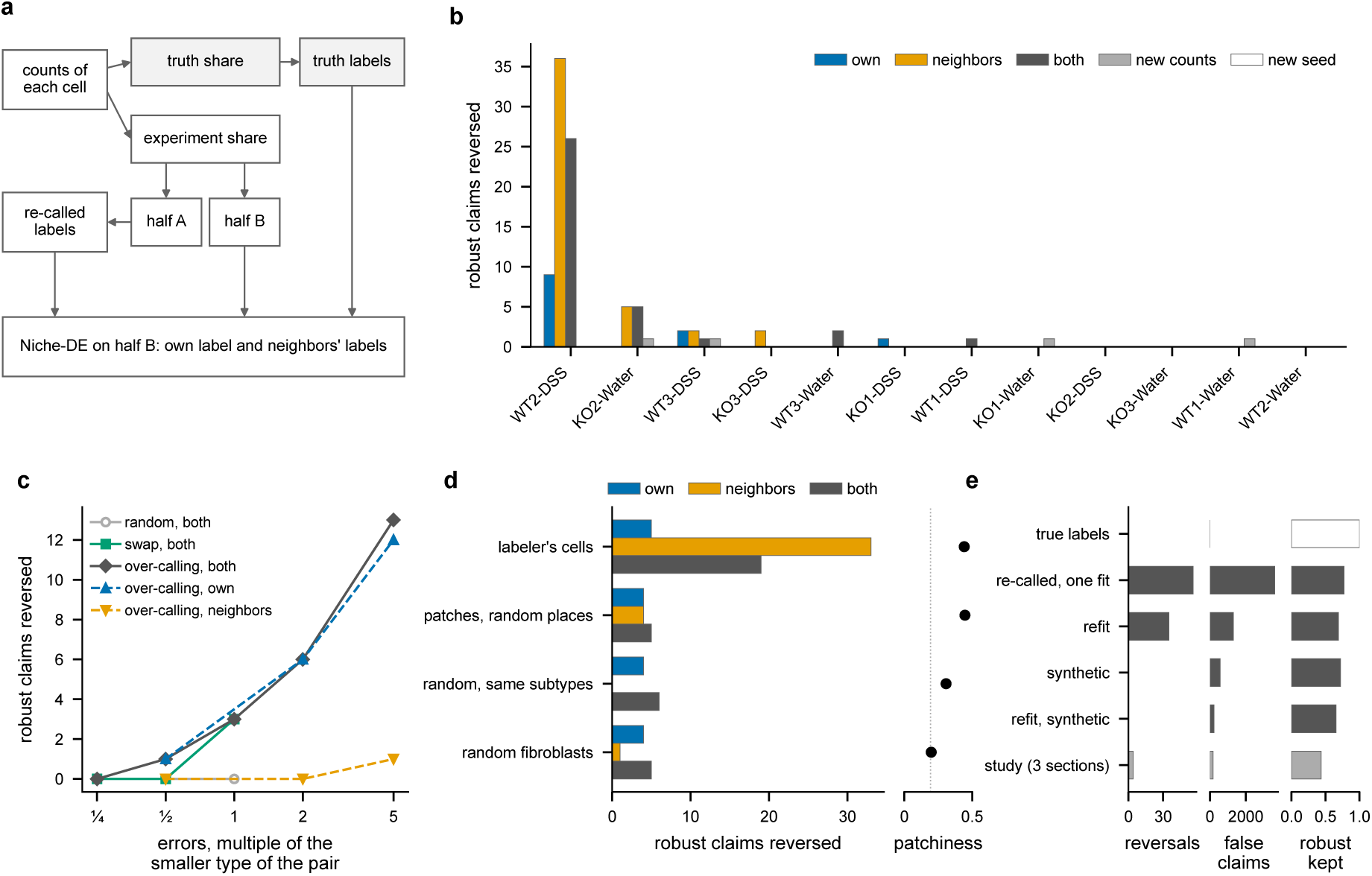
Label error reverses Niche-DE claims on semi-synthetic colon data. **a**, Design. The truth labels come from the truth share of the counts; the experiment share is split into halves A and B. Labels re-called on half A enter the model in the cell’s own role, in the neighbors’ role, or in both; Niche-DE is fitted on half B. **b**, Robust claims reversed per section, by the role of the re-called labels, beside the reversals of any true-label call by new counts and by a new seed. **c**, Injected errors: reversals against the number of errors, for errors spread at random, swapped within the most confused pair, or over-calling its smaller member, in the cell’s own role and in the neighbors’ role. **d**, Inflamed section: reversals by role for the labeler’s over-call of mIAFs at its own cells, as patches of the same tightness at random places, at random cells of the same subtypes, and at random fibroblast cells; patchiness (share of mislabeled cells among the 20 nearest neighbors of a mislabeled cell) beside each bar. **e**, Agreement filters: reversals, false claims, and robust claims kept, for one fit with the re-called labels, a refit on a new split, agreement with a fit on the study’s labels (three sections) or on the synthetic published labels, and the refit with agreement; the true-label row is the reference, which keeps every robust claim by definition. The synthetic published labels reversed no claim; agreement with them, therefore, removes every reversal by construction. Sections are named by genotype (KO, fibroblast-specific deletion of *Glis3* ; WT, floxed controls), replicate, and treatment (DSS or water).

The reversals pointed to one subtype, inflammation-associated fibroblasts (mIAFs). In the semi-synthetic truth, this class is large because its labeler had a flat prior; the study’s labels hold few such cells. We use it as a truth class rather than as a claim about biology. In every section, the labeler called mIAFs on 1.5 to 3.9 times as many cells as the truth held, and most of the extra cells came from two other fibroblast subtypes. The over-call was largest in the section with most reversals, where the labeler called mIAFs on about half of the cells of three other subtypes, including the two largest sources of extra cells. The over-call came mostly from what the labeler had learned. It had learned its subtypes from the synthetic published labels, which a labeler of the same kind had called on the whole experiment share, as a study’s own labels would be called. Trained on the study’s labels instead, at either depth of its training counts, it over-called mIAFs less, and little in the inflamed section. Random errors in the training labels, at the same rate, did not reproduce the over-call. The truth’s own labeler, with centroids from the experiment share and run on the same half of the counts as the re-called labels, mislabeled up to about a fifth of the cells, yet reversed almost no claim (Supplementary Note 4). Its errors were mostly of the counting-noise kind, and they left the claims standing; the reversals came from what the labeler had learned.

The published labels of the original study, called on all counts, including the half the model saw, drew the boundary of the same subtype the other way and called few cells mIAFs. They, too, reversed robust claims in the three sections that held enough of their mIAFs to test, mostly claims with mIAFs as the niche type (Supplementary Note 4). Niche claims about this subtype, therefore, depended on where the labels drew its boundary, in either direction.

To find the cause, we injected errors into the true labels in fixed numbers and changed only where they fell (Fig. 6c). Errors spread at random over the fibroblast subtypes reversed no robust claim at any dose, although they created claims that true labels did not make. Errors concentrated on one subtype, by swapping it with its most confused partner or by over-calling it, reversed robust claims, more so at higher doses. Nearly every reversed claim had that subtype as its index type and came through the cell’s own label. The same errors in the neighbors’ labels reversed almost no claim, even where most cells carrying the over-called label belonged to other types.

The labeler’s own errors acted differently. In the inflamed section, we injected only its overcall of mIAFs into the true labels. In the neighbors’ labels, it reproduced most of the reversals of the full re-called labels; in the cell’s own label, it reversed few (Fig. 6d). We then moved the same errors to random cells of the same subtypes, or of any fibroblast subtype. Reversals through the own label persisted at about the same number, and reversals through the neighbors’ labels almost all disappeared. The labeler’s errors lay in patches: the neighbors of a mislabeled cell were more often mislabeled than those of a moved error. Patches of the same tightness at random places did not restore the reversals. Outside the inflamed section, the over-call of mIAFs alone reversed no claim, and the few reversals there came from other errors of the labeler. Whether a niche claim reverses, therefore, depends on which labels are incorrect and where they lie.

### 2.7 Checks without the truth removed most claims made by label error

We then asked whether a user could remove these claims without knowing the truth. Refitting on a new split of the counts, with labels re-called on its other half, removed more than a third of the reversals and most of the false claims; label error made the rest again on the new split. Keeping only claims that a fit with a second annotation also made removed more. With the study’s labels as the second annotation, the filter removed nearly all reversals and most false claims in the three sections where we could test it, at the cost of more than a third of the robust claims that the single fit kept there (Fig. 6e; Supplementary Table S21). Those labels drew the boundary of mIAFs the other way. With the synthetic published labels as the second annotation, the filter removed every reversal, as it had to: those labels reversed no claim on their own, and the labeler had learned from them (Supplementary Note 4).

## 3 Discussion

A cell-type label enters every estimate of a neighborhood effect twice, and errors in the cell’s own label inflate the estimate. The mechanism is residual confounding by a mismeasured covariate [5, 6]: the neighborhood receives the part of the cell-type effect that the labels miss. Ecology and the social sciences separate composition from context in the same way [25, 26]. We expected this direction, but the estimate does not force it. Errors that follow the neighborhood offset the inflation, and the response that detects it can vanish or turn negative where cell states form a continuum. The inflation nonetheless appeared in every simulated regime. In real tissue, the response was positive in every measured population with identity to transfer but one, the thick ascending limb on the deepest kidney panel, where it was close to zero.

Standard practice obscures the inflation. Labeling and modeling the same counts absorbs noise into identity and lowers the neighborhood share. Selecting the population by labels changes which cells are studied. A published neighborhood share, therefore, carries two biases of opposite sign, and neither the protocol nor the semi-synthetic check gives their sum. An estimate with labels called on one half of the counts and the model fitted on the other has no circularity. Its labels are less stable, and its inflation is larger (Fig. 2a), but that inflation is the bias that the protocol measures. Reported with its own-label channel next to the published estimate, the split estimate is the one whose bias a reader can check.

We recommend four practices.

1. Call labels and fit neighborhood models on independent halves of each cell’s counts. The split removes the circularity of the estimate rather than the systematic errors of a labeler, which both halves share.
2. Define the population under study with the most stable labels available, keep it fixed, and report the membership channel with the estimate.
3. Report the own-label channel with every estimate of a neighborhood effect, and the contrast in label stability with every contrast between conditions. Report the channel as a direction. Give a size only from the semi-synthetic check, run with the labeler actually used on the same data.
4. Report effect sizes rather than call sets. Refit niche methods on a new split of the counts to remove claims that depend on the draw of the counts. Where a second annotation that errs differently is available, keep only claims that a fit with it also makes; in our data, this removed most claims made by label error, at a cost in claims that true labels supported. Errors spread at random test neither.

A claim about a specific niche deserves a further check because label error removes identity mostly from neighborhoods with an uneven mix of confusable subtypes, and their label composition ranks them.

### Which labels are incorrect, and where

In our measure, own-label error inflates the neighborhood share. In the calls of Niche-DE, label error reversed claims by two routes. Through the cell’s own label, errors concentrated on one subtype carried the expression of other types into it, and their positions mattered little. Through the neighbors’ labels, only the labeler’s own errors reversed claims; the same errors at random cells of the same subtypes reversed almost none. The labeler’s errors differed from the moved ones in two ways. They lay in patches, and they were the cells whose expression the labeler took for the over-called subtype. Patches of the same tightness at random places reversed few claims. Our controls do not separate the cells that the labeler chose from the places of their patches. Patches do not contradict the occurrence test, in which neighborhood composition barely predicted which cells changed label: cell states, too, vary in space. The neighbors’ labels changed our measure little, yet carried most reversals of Niche-DE calls in the inflamed section. The thinning estimate of the neighbor channel remains a sensitivity (Supplementary Note 3), and the Niche-DE comparisons rest on true labels rather than on that estimate. Errors spread at random over subtypes, a common check of robustness, reversed no claim at the same number of errors; moved to random cells, the labeler’s errors no longer reversed claims through the neighbors’ labels. Spread over subtypes, they dilute the errors on any one subtype; spread over cells, they miss where the labeler’s errors lie. A claim that survives such a check, therefore, has not faced the errors that labelers make. The companion study finds a related result for segmentation, where random relabeling moves contact statistics against the direction of the true errors [24].

### Reducing label error in niche calls

A refit on a new split of the counts removed claims that depended on the draw of the counts, but label error made most of its reversals again on the new split, such that a refit alone could not separate it from niche biology. Agreement with a fit on the study’s own labels removed nearly all reversals where we could test it, at the cost of more than a third of the robust claims that a single fit kept. Those labels drew the boundary of mIAFs the other way, such that their errors and the labeler’s rarely made the same claim. The synthetic published labels, from which the labeler had learned, made fewer errors than the labeler, and different ones: the labeler that learned from them inflated mIAFs rather than copying their claims. In the colon data, random errors in the training labels did not reproduce the over-call (Supplementary Note 4, with the verdict of the rule in two further datasets). The truth’s own labeler, with centroids from the experiment share and run on the same half of the counts as the re-called labels, reversed almost no claim and made half as many false claims in all, although it still mislabeled up to about a fifth of the cells (Supplementary Note 4). The truth labels came from the same labeler at full depth; its errors were, therefore, mostly of the counting-noise kind. The thinning diagnostic and the agreement check, therefore, cover different errors: the first measures the noise-like part of label error, and the second can remove claims made by systematic error when a second annotation errs differently.

### Subtypes defined by position

Identity exists only at the level at which cells are annotated. Some subtypes are partly defined by position, such as hepatocyte zones or fibroblasts along the colonic crypt axis. For them, label error moves variance across a boundary that the annotation drew, and the bias is defined only relative to that boundary. The inflation did not depend on such subtypes. It also appeared where the target set consisted of whole lineages: immune and stromal lineages in lung cancer, and the proximal tubule with the thick ascending limb in kidney (Supplementary Table S6). On the deepest kidney panel, where labels rarely changed, the channel was close to zero. Coarse annotation adds to the bias. In simulations, labels from clusters that merged the subtypes under study inflated the neighborhood share more than per-cell labels did (Supplementary Table S13).

### Three kinds of label error

Thinning adds errors that behave like counting noise. Segmentation adds a different error (Table 1). Whole-cell counts carry transcripts of cells in contact with the target cell, such that small or thin cells take the type of their larger neighbors [24]. The two errors fall mostly on different cells. Segmentation removes labels mainly from small or thin cells, such as T/NK, B, mast, and mural cells, whereas the error in most of our target sets is a choice between subtypes of one lineage. The kidney stromal set, which includes vascular smooth muscle and mesangial cells, is exposed to both. The two errors meet in lung cancer, where we labeled each cell from its whole-cell and from its nucleus counts [10]. Nucleus labels behaved as thinning predicted in the cell’s own role but not in the neighbor role (Supplementary Note 2; Supplementary Fig. S5). Thinning, therefore, measures the noise-like part of label error. The part from segmentation has to be measured from nucleus and whole-cell labels of the same cells. Thinning adds neither segmentation error nor a third kind, systematic error of the labeler, which reversed Niche-DE calls (Section 2.6).

**Table 1:** Three kinds of label error. Thinning adds counting noise, which this study measures. It does not add systematic error of the labeler, which reversed Niche-DE calls on semi-synthetic data. The companion study measures segmentation error [24]. Lung cancer, labeled from whole cells and from nuclei, carries counting noise and segmentation error.

| Error | Falls on | Effect | Measured by |
|---|---|---|---|
| Counting noise | confusable subtypes of one lineage; which cells change barely depends on the neighborhood, but the new label tends to be the one typical of it | inflates the neighborhood share through the cell's own label; spread at random over subtypes or made by the truth's own labeler, reversed almost no Niche-DE claim | thinning, with count splitting and fixed targets (`nichepeek`) |
| Systematic labeler error | one subtype whose boundary the labeler learned from another labeler's calls; the errors lie in patches | reverses Niche-DE claims, mostly through the neighbors' labels; moved to random cells of the same subtypes, it no longer does so through the neighbors' labels | semi-synthetic truth, with Niche-DE fits (this study) |
| Segmentation | small or thin cells bordering larger cells of another type; the label follows the neighbor in contact with the cell | removes labels of the small type and lowers their contacts | nucleus and whole-cell labels of the same cells (`claimbreak`) |

Transcripts donated by neighbors create niche associations even with correct labels. Neither study removes them, and count splitting cannot, because both halves share them.

### Relation to other work

Methods that correct or flag transcript spillover [8, 10, 27] clean expression given each cell’s label; SPLIT changes a label only by swapping a cell’s primary and secondary types [10]. ResolVI relabels cells with a classifier trained on the existing labels and propagates label uncertainty into neighborhood composition [9], which is the neighbor role. The own-label role, which dominated our measure, is outside that treatment. In the calls of Niche-DE, the neighbor role carried most reversals, and only the labeler’s own errors produced them. Methods that estimate context-dependent expression conditional on labels or composition [4, 28–30] fit labels and context model to the same data, which is the problematic regime that the protocol diagnoses. CSDE corrects differential expression for errors in segmentation and annotation with a small set of cells validated by an expert [31]; our protocol needs no validated cells. Label-free models of niches [32–34] avoid the neighbor role. Whether they avoid the own-label route depends on how they identify intrinsic state. Our diagnostic belongs to the simulation–extrapolation family [23, 35, 36]. We apply it to spatial neighborhood effects, decompose it by the role a label plays, and validate its direction on semi-synthetic data built from real tissue.

### Limitations

Our labels come from classifiers rather than from the pipelines that produced the published labels. The diagnostic measures the response to error of the kind that the re-calling labeler makes. Systematic disagreement between a labeler and the labels it learned is a part that thinning does not see (Supplementary Note 3). The published labels of the colon study and its truth labels disagreed in this way about one subtype, and the disagreement reversed Niche-DE calls (Section 2.6). Two labelers of different kinds agreed in sign but not always in size (Supplementary Note 2). We tested the channels on our own measure, on one niche regression of the NCEM type in our own implementation, and on the calls of Niche-DE (Methods). NCEM itself and MISTy remain untested, and the size of the bias need not transfer to them. The Niche-DE tests used semi-synthetic truth from one tissue, splits of the same sections, and labelers of one kind with a flat prior, which learned their subtypes from labels of a labeler of the same kind or from the study’s labels. With the study’s label frequencies as its prior, the labeler of the truth labels called no cell mIAFs (Supplementary Note 4). The Niche-DE fits ran at a quarter of native depth, where counting noise alone changed most calls; we did not measure reversals at native depth. Most reversals came from one section, and we tested placement there only. We tested how label error changes the calls of a method rather than the conclusions of a published study. The agreement check with an independent annotation rests on the study’s labels in three sections; those labels were called on counts that include the half the model saw. In four populations, the middle thinning level left only 12 to 13 counts per cell, just above the depth of about ten below which the channels can change sign. Count splitting halves the depth that the model sees, assumes Poisson-like technical noise, and cannot remove segmentation spillover, which both halves share. Where cell states form a continuum, the response to added error carries no sign information.

### Conclusions

Errors in a cell’s own label make part of its type look like an effect of its neighborhood, and standard practice can hide, exaggerate, or reverse this bias. Splitting the counts, fixing the target population, and reporting the own-label channel reveal its direction, steps which nichepeek implements. In the calls of Niche-DE, claims reversed when a labeler’s errors concentrated on one subtype or lay where the labeler put them. Checks that spread errors at random, therefore, miss the errors that matter.

## 4 Methods

### 4.1 Data

We analyzed six published datasets (Table 2). Mouse colon stroma was profiled with Xenium in mice treated with dextran sulfate sodium (DSS) or water [18]: mice with a floxed *Glis3* allele, with (KO) or without (WT) *Pdgfra*-Cre, a fibroblast-specific deletion, given water or chronic DSS. Sections are named by genotype, replicate, and treatment (for example, WT2-DSS). Targets were the fibroblast subtypes. Mouse colon during DSS colitis was profiled with MERFISH [19, 37]. We used the published second-tier labels, raw counts, and 16 sections of 6 mice from the healthy, day-9, and day-21 groups. Targets were the non-lamina-propria fibroblast subtypes (fibroblasts 1, 4, 5, 7, 12, and 15, and IAF 5). Blank barcodes and two gene aliases (*Dclk1*, *Gdf1*) were excluded. Healthy and fibrotic human liver were profiled with MERFISH [20]. Targets were the four hepatocyte subtypes, and sections of more than 30,000 cells were analyzed as two crops of 30,000 cells. Human rheumatoid arthritis synovium was profiled with Xenium [21]. Targets were the fibroblast, myeloid, or endothelial subclusters. Each sample was analyzed as two crops of 150,000 cells, and the three populations were analyzed on identical crops. Crops are the 30,000 or 150,000 cells nearest to centers chosen by farthest-point sampling, and they can overlap. Diabetic kidney disease was profiled with CosMx and Xenium Prime 5K in samples from the same patients [22]; we used the 4 patients whose samples were imaged on both, the published labels (annotation_updated), and three versions of the data: CosMx (998 genes), Xenium restricted to the 656 genes on both panels, and the full Xenium panel (5101 genes). Targets were the endothelial subtypes, the stromal types (fibroblasts, vascular smooth muscle, and mesangial cells), proximal tubule with injured proximal tubule, and thick ascending limb with injured thick ascending limb. A fifth set had two target labels, one for each lineage: proximal tubule and thick ascending limb (Supplementary Table S6). Single-label targets (fibroblasts, proximal tubule) were run as a negative control (2 proximal-tubule crops with fewer than 50 target cells excluded, as nichepeek 0.3.5 does by default). Sections were analyzed as up to two crops of 30,000 cells. Non-small-cell lung cancer was profiled with Xenium on three panels [10]: the lung panel (289 genes), the immuno-oncology panel with a custom add-on (340 genes), and the Prime 5K panel (5000 genes). Every cell was labeled from its whole-cell and from its nucleus counts by a multinomial likelihood against reference profiles of nine cell types (tumor subtypes merged), built from Chromium Flex profiles of lung cancers of four patients [38]. The whole-cell labels served as published labels and the nucleus labels as a second label set (--truth-key). The predicted effect of the nucleus labels was, per section, the slope through the origin of a channel against the added flip fraction over the three thinning levels, times the fraction of targets on which re-called and nucleus labels disagreed; the observed effect is the difference between the arms with nucleus and with re-called labels in that role. Sections were analyzed as two crops of 30,000 cells. In every dataset, labels with fewer than 50 cells were removed.

**Table 2:** Datasets and settings. Units, independent mice, donors, or patients; *n*, sections or crops; depth, native median counts per cell, before splitting or thinning; thinning levels *p* applied to half *B* (middle level in bold).

| Population | Platform | units | *n* | depth | thinning levels |
|---|---|---|---|---|---|
| Mouse colon stroma | Xenium | 12 | 12 | 293 | 0.5, **0.25**, 0.12 |
| Mouse colon fibroblasts | MERFISH | 6 | 16 | 72 | 0.6, **0.35**, 0.2 |
| Human liver, healthy | MERFISH | 3 | 4 | 361 | 0.35, **0.15**, 0.06 |
| Human liver, fibrotic | MERFISH | 3 | 6 | 42 | 0.8, **0.6**, 0.35 |
| Synovial fibroblasts | Xenium | 10 | 20 | 98 | 0.6, **0.35**, 0.2 |
| Synovial myeloid | Xenium | 10 | 20 | 98 | 0.5, **0.25**, 0.12 |
| Synovial endothelial | Xenium | 10 | 20 | 98 | 0.5, **0.25**, 0.12 |
| Human kidney (5 target sets) | CosMx | 4 | 7 | 89 | 0.8, **0.6**, 0.4 |
| Human kidney, shared genes | Xenium | 4 | 8 | 85 | 0.8, **0.6**, 0.4 |
| Human kidney, full panel | Xenium | 4 | 8 | 412 | 0.8, **0.6**, 0.4 |
| Lung cancer, lung panel | Xenium | 4 | 10 | 70 | 0.8, **0.6**, 0.4 |
| Lung cancer, immuno-oncology | Xenium | 4 | 10 | 148 | 0.8, **0.6**, 0.4 |
| Lung cancer, Prime 5K | Xenium | 2 | 4 | 209 | 0.8, **0.6**, 0.4 |

### 4.2 Ladder and noise correction

Each target cell is described by the label composition of its *k* = 20 nearest neighbors (all cells of the section or crop), square-root transformed, and targets are grouped by k-means (12 groups, best of 5 initializations). *R*^2^ is the share of expression variance of target cells explained by the groups, micro-averaged over genes on size-normalized counts, with the sampling variance removed from the total and from the within-group variance and negative per-gene residual variances clipped at zero. Because size factors are computed from the same counts, counts are multinomial given the cell total, and the sampling variance of gene *g* in cell *i* is estimated without bias by *Y_ig_*(*t_i_ − Y_ig_*)*/*((*t_i_ −* 1)*s*^2^), with *t_i_* the total and *s_i_* the size factor. The rungs are *S*_0_, the observed tissue; *S*_1_, expression permuted among cells with the same own label and target status, with the groups of *S*_0_ retained; and *S*_2_, cells (label, expression, and target status together) relocated at random, with groups re-fitted from the *S*_0_ centers. *S*_1_ is averaged over 20 permutations (a single permutation in the kidney and lung runs). Identity is *I* = *R*^2^(*S*_1_) *− R*^2^(*S*_2_), and the neighborhood share is *V* = *R*^2^(*S*_0_) *− R*^2^(*S*_1_). Random numbers for k-means, permutations, and thinning are shared between arms of a section so that differences between arms are not differences between random draws.

### 4.3 Count splitting, arms, and channels

Each cell’s counts are split as *Y ^A^ ∼* Binomial(*Y,* 1*/*2), *Y ^B^* = *Y − Y ^A^*, which for Poisson counts gives independent halves that share the cell’s expression [16, 17]. Biological variation between cells, including overdispersion relative to Poisson, is shared by the two halves and is signal here; only the sampling noise must be independent. Where technical noise itself is overdispersed, negative binomial splitting would be required [39]. Where transcripts are assigned to nuclei, the counts to split are those of the set from which the labels are called. Where a nuclear half is too sparse to label [24], the whole-cell counts are split instead. Labels are re-called on *Y ^A^* (giving *c*_1_) and on *Y ^A^* thinned to *p* (*c_p_*), and every arm is scored on *Y ^B^* and on *Y ^B^* thinned to *p*. Target cells are fixed by the published labels *a* in every arm except the membership arm. Each arm is a ladder with a choice of labels in the own role and in the neighbor role:

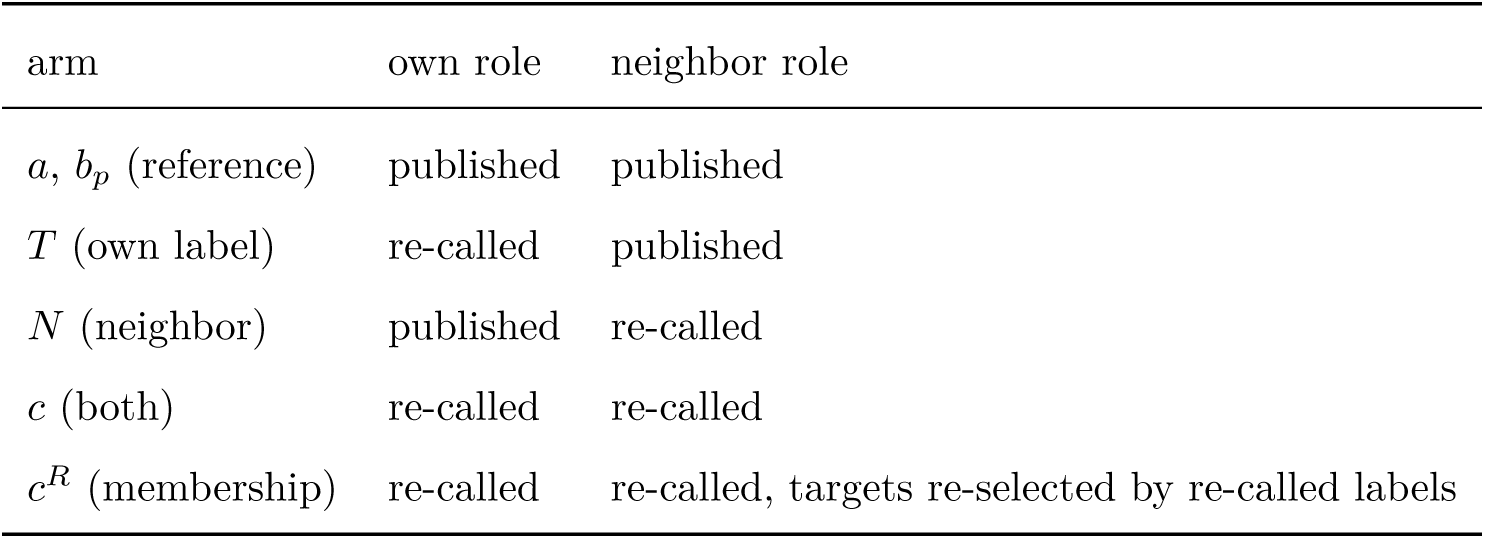

| arm | own role | neighbor role |
|---|---|---|
| *a*, *b<sub>p</sub>* (reference) | published | published |
| *T* (own label) | re-called | published |
| *N* (neighbor) | published | re-called |
| *c* (both) | re-called | re-called |
| *c<sup>R</sup>* (membership) | re-called | re-called, targets re-selected by re-called labels |

For an arm *X* scored at full and thinned depth, the label effect is *L_X_*(*p*) = (*X_p_ − X*_1_) *−* (*b_p_ − a*); the difference removes the effect of depth on the estimator. The own-label, neighbor, and net effects are *L_T_*, *L_N_*, and *L* = *L_c_*. The interaction is *L_c_ − L_T_ − L_N_*, and the membership channel is *L_M_* = *L_c_R − L_c_*. Because the own-label channel cannot change *R*^2^(*S*_0_), we used it to check the implementation (maximum *|L_T_* (*S*_0_)*|* = 0 in every section of every dataset). The own-label channel on *V* is, therefore, minus the change in *R*^2^(*S*_1_), the variance that the own labels explain across the groups. Circularity is the difference between labels *c*_1_ scored on *Y ^A^*, the counts from which they were called, and on *Y ^B^*. Because a single split adds noise of its own, every section was analyzed with 3 independent splits and thinnings, and section-level quantities were averaged. The standard deviation between single splits ranged from 0.003 (Xenium stroma) to 0.014 (MERFISH colon) for the net label effect at the middle thinning level. The standard design (Fig. 3c,d) calls labels and scores the ladder on the full counts and re-selects targets by the re-called labels. Table 3 collects the quantities of this and the next subsection.

**Table 3:**
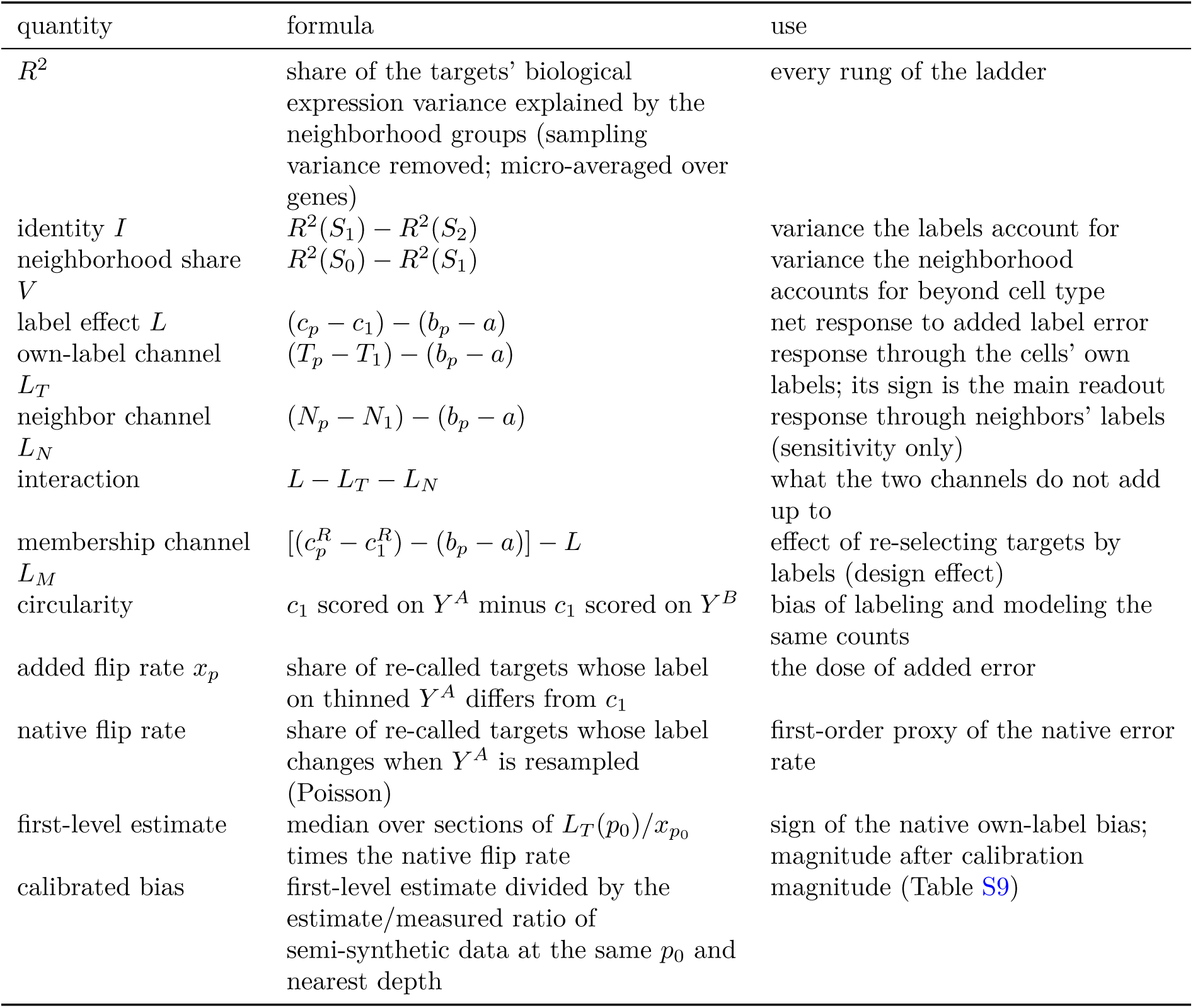
Quantities of the protocol, their formulas, and their use. *X_p_*, the ladder quantity (*V* unless stated) of arm *X* on counts thinned to *p*; *X*_1_, the same at full depth of half *B*.

| quantity | formula | use |
|---|---|---|
| *R*<sup>2</sup> | share of the targets' biological expression variance explained by the neighborhood groups (sampling variance removed; micro-averaged over genes) | every rung of the ladder |
| identity *I* | *R*<sup>2</sup>(*S*<sub>1</sub>) − *R*<sup>2</sup>(*S*<sub>2</sub>) | variance the labels account for |
| neighborhood share *V* | *R*<sup>2</sup>(*S*<sub>0</sub>) − *R*<sup>2</sup>(*S*<sub>1</sub>) | variance the neighborhood accounts for beyond cell type |
| label effect *L* | (*c*<sub>*p*</sub> − *c*<sub>1</sub>) − (*b*<sub>*p*</sub> − *a*) | net response to added label error |
| own-label channel *L*<sub>*T*</sub> | (*T*<sub>*p*</sub> − *T*<sub>1</sub>) − (*b*<sub>*p*</sub> − *a*) | response through the cells' own labels; its sign is the main readout |
| neighbor channel *L*<sub>*N*</sub> | (*N*<sub>*p*</sub> − *N*<sub>1</sub>) − (*b*<sub>*p*</sub> − *a*) | response through neighbors' labels (sensitivity only) |
| interaction | *L* − *L*<sub>*T*</sub> − *L*<sub>*N*</sub> | what the two channels do not add up to |
| membership channel *L*<sub>*M*</sub> | $[(c_p^R - c_1^R) - (b_p - a)] - L$ | effect of re-selecting targets by labels (design effect) |
| circularity | *c*<sub>1</sub> scored on *Y*<sup>*A*</sup> minus *c*<sub>1</sub> scored on *Y*<sup>*B*</sup> | bias of labeling and modeling the same counts |
| added flip rate *x*<sub>*p*</sub> | share of re-called targets whose label on thinned *Y*<sup>*A*</sup> differs from *c*<sub>1</sub> | the dose of added error |
| native flip rate | share of re-called targets whose label changes when *Y*<sup>*A*</sup> is resampled (Poisson) | first-order proxy of the native error rate |
| first-level estimate | median over sections of *L*<sub>*T*</sub>(*p*<sub>0</sub>)/*x*<sub>*p*<sub>0</sub></sub> times the native flip rate | sign of the native own-label bias; magnitude after calibration |
| calibrated bias | first-level estimate divided by the estimate/measured ratio of semi-synthetic data at the same *p*<sub>0</sub> and nearest depth | magnitude (Table S9) |

### 4.4 Classifier, flips, and context tests

Labels are re-called by a nearest-centroid classifier. Counts are scaled to a common total and log-transformed, and each cell is assigned the label whose centroid has the highest correlation with its profile. Centroids are the mean profiles of the published labels on the full counts of all other physical sections so that no cell contributes to the centroids that label it. Crops of the same section are excluded together. The second labeler is multinomial logistic regression (scikit-learn, *C* = 1) on log-normalized counts. It is trained leave-one-section-out on the published labels of at most 400 cells per label, each binomially thinned to a fraction drawn uniformly between 0.05 and 0.6, so that the training depths cover those at which it is applied. Two variants with natural class priors were run on colon stroma: the nearest-centroid correlations divided by a temperature and offset by the log published label frequency of the other sections, with the temperature fitted by maximum likelihood of the published labels on half-depth cells of the other sections (0.0218), and the logistic regression trained on a uniform sample of cells of the same total size. These runs used 3 splits and 20 permutations at *S*_1_ per section, and the comparison between conditions is made on per-mouse values (Uncertainty). The native flip rate is the fraction of re-called target cells whose label changes when their half *A* counts are resampled (Poisson, three resamples). Agreement across subsamples of cells is also used to stabilize labels and bound their error [40]. Our flip rate, like such agreement, measures stability rather than accuracy, which is why we calibrate it against known labels (Semi-synthetic validation). The added flip rate *x_p_* is the fraction whose label on thinned half *A* counts differs from *c*_1_. The first-level estimate of the native bias is the median over sections of (*L_T_* (*p*_0_)*/x_p_*_0_) times the native flip rate, with *p*_0_ the first thinning level. Its calibration is described below. The occurrence test predicts which target cells change label between *c*_1_ and *c_p_*, by logistic regression on the cell’s own label and depth, with and without its neighbor composition. It reports the gain in area under the curve, holding out whole physical sections, and has the structure of the neighbor-prediction regressions used to detect spillover [8, 10]. The destination test considers target cells whose re-called label agrees with the published one and whose label changes on thinning. It asks whether the new label’s typical neighbor composition is closer to the cell’s own neighbor composition than the old label’s. The null assigns the same transitions to random cells with the same label (50 draws).

### 4.5 Niche regression

In each arm, the size-normalized expression of target cells was regressed by ridge regression (*λ* = 1) on the one-hot own label, on the square-root neighbor composition (*k* = 20), or on both. *R*^2^ is the out-of-fold share of variance, with five folds within each section. It is micro-averaged over the 200 genes with the highest mean count in the published targets, the same genes in every arm, with the multinomial sampling variance removed. The niche effect is *R*^2^(both) *− R*^2^(own label).

### 4.6 Uncertainty

Sections or crops of the same mouse, donor, or patient are not independent, and crops of one sample can overlap. Dataset-level statistics are medians over sections. Their uncertainty is a cluster bootstrap over independent units in which units are resampled with replacement together with all their sections. Signs are tested with a two-sided sign test over unit means. Per-mouse values of the two colon-stroma conditions (six mice each) are compared with the exact two-sided Mann–Whitney test, whose smallest attainable *P* with complete separation is 2*/*924 *≈* 0.002. With fewer than five units (liver), we report medians or unit values without intervals or tests. In one synovial sample and one healthy liver donor, the two crops shared 0.98 and 0.89 of their cells, respectively. Crops of other samples overlapped less or not at all. All analyses were exploratory.

#### Condition contrasts

For each condition *c*, *B_c_* = *s_c_f_c_*, where *s_c_* is the through-origin slope of *L_T_* on *x_p_* over the condition’s sections and levels with at least ten counts per cell, and *f_c_* the median native flip rate over units. The label-error term of a contrast is *B*_alt_ *− B*_ref_; the depth-matched contrast reads the deeper condition’s published neighborhood share at the thinning level closest to the other’s native depth. Intervals resample units within each condition. With fewer than five units per condition, none are reported, as for all other statistics.

### 4.7 Simulations

Simulated sections contain six non-target and four target cell types in two families, with spatially structured types (a layer axis, inflammation patches, and within-family segregation) and 150 genes. A planted neighborhood signal shifts 20 genes in target cells in proportion to the standardized local fraction of immune cells among the 20 nearest neighbors, with strength *α*. Expression has cell-level biological variation, and counts are Poisson at a depth that varies log-normally between cells. Labels are called with the nearest-centroid classifier from a deep reference. The grid varied depth (35, 70, and 140 counts per cell), the separation of target families, *α* (0 and 0.5), and spatial segregation, with two sections per setting and 72 in total. Along-axis experiments moved target cells part of the way toward the partner type of the other family, by *κ ∈ {*0, 0.6, 1*}* times a variable drawn per cell (random) or given by the local immune fraction (following the neighborhood), with 16 sections per regime. Channel arms used true labels as the reference. Robustness experiments either added a gamma-distributed technical multiplier to each count’s rate (squared coefficient of variation 0.1 or 0.3), not shared between independent draws, or moved a fraction (0.1 or 0.2) of each gene’s counts to one of the six nearest cells. They compared the split halves with an independent replicate at half depth to measure the circularity left by splitting (six sections per setting and regime). To test conditioning on expression, *S*_1_ permuted expression within k-means clusters (4 to 32) of the targets’ log-normalized half *A* counts instead of within labels. A further experiment weakened the marker genes of the six non-target types (strength 2.5, 1.2 or 0.8; six sections each) to vary how often neighbors’ labels are incorrect. All simulations used the multinomial noise correction applied to the real data.

Geometry simulations used three sections of 6000 cells per tissue, divided into Voronoi niches of about *m* cells (*m ∈ {*8, 30, 120, 300*}*). Each niche held *D* of four target subtypes in two confusable pairs (*D ∈ {*1, 2, 4*}*, equal shares), and non-target cells whose mix varied between niches. A planted neighborhood signal made 20 target genes depend on the local share of one non-target type. Optional spillover replaced a fraction *s* of each cell’s expected profile by the mean of its six nearest neighbors. “Published” labels were called by nearest centroid on the same counts, and the own-label bias was measured with the truth arms described below (3 independent tissues per setting). A separate test counted how often target cells were labeled as their confusable partner type for cells with and without that type among their six nearest neighbors.

#### Exact-truth checks of published labels

To measure what labels called and scored on the same counts carry, we called labels with the nearest-centroid classifier on all counts of each simulated cell and scored the ladder on the same counts, against the true labels (108 sections over three depths, three identity strengths, and tissues with and without a planted neighborhood signal). For cluster-based labels, cells were clustered (30 principal components, a 15-nearest-neighbor graph, Leiden clustering at resolutions 0.5, 1.5, and 3), and each cluster was annotated by its nearest reference centroid (18 sections).

### 4.8 Semi-synthetic validation

For colon stroma (all sections) and healthy liver (the three healthy donors), each cell’s counts were split binomially into a truth share *Y ^T^ ∼* Binomial(*Y, f*) and an experiment share *Y ^E^* = *Y − Y ^T^*, and the experiment share was thinned to Binomial(*Y ^E^, q*). Settings with the same *f* share the split and differ only in *q*. True labels were called on *Y ^T^* and synthetic published labels on the thinned *Y ^E^*, both by the nearest-centroid classifier with centroids from the full-depth counts and published labels of the other physical sections. nichepeek was run on the thinned *Y ^E^* with the synthetic published labels, exactly as on real data, with three additional arms on half *B* of the same split: true labels in both roles (*tt*), true own labels with published neighbor labels (*t*), and re-called own labels with true neighbor labels (*tc*). The directly measured own-label bias is *T*_1_ *− t* (re-called against true own labels, neighbors published), the net bias *c*_1_ *− tt* and the neighbor bias *c*_1_ *− tc*, all on the neighborhood share and at the depth of the re-called labels *c*_1_. The first-level estimate was computed per section as in the real data, and its ratio to the measured bias from the medians over sections. Liver sections were cropped as in the real runs.

To estimate the effect of errors in the truth labels, we also called truth labels on the truth share thinned further to *r* = 0.8 and 0.6 of its counts. The truth arms were scored again with each thinned truth, and the measured bias was extrapolated to a truth flip rate of zero by a linear fit per unit, then summarized as the median over units. Quadratic fits and fits to the medians over units served as sensitivity analyses. An extrapolation was used only if the truth flip rate rose, from *r* = 1 to *r* = 0.6, by at least a quarter of its value at *r* = 1. A further setting (*f* = 0.4, *q* = 1) placed the ladder at about 88 counts per cell in colon and 108 in liver.

Published labels were also scored against the truth labels (Supplementary Table S12). In each section, the neighborhood share was computed twice on all counts of the experiment share, once with the synthetic published labels and once with the truth labels, each in both the own and the neighbor role, with the same seed, groups, and permutations. The net effect of the published labels is the difference; the own-label channel replaced only the own labels by the truth labels. Target cells were those with the published target label in both computations. For colon stroma and healthy liver, we scored the semi-synthetic files of this subsection. For these two populations and for colon fibroblasts, fibrotic liver, and the three synovial populations, we also built the semi-synthetic data from the real data with the arguments of each population’s main run (*f* = 0.5, *q* = 1), with the split of nichepeek mcsimex --semisynthetic (Magnitude check across populations) and the nearest-centroid labeler described above. Liver and synovium were scored in the crops of their main runs, and each crop counts as a section in the table.

### 4.9 Niche-DE on semi-synthetic data

We used the semi-synthetic colon stroma data of Section 4.8, all twelve sections. We split the experiment share of each cell into halves A and B by binomial thinning, with three independent splits. We fitted Niche-DE [4] (development version 0.0.0.9000, GitHub kaishumason/NicheDE, commit 87e0e89, with Seurat removed from the declared dependencies; the same build on both machines), in R 4.6.1, on half B with kernel bandwidths of 175, 250, and 300, in units where the median distance to the third nearest neighbor is 100; that distance was 20 to 25 *µ*m across sections. Effective niches came from CalculateEffectiveNicheLargeScale. The fitting step used *C* = 150, *M* = 50, and *γ* = 0.8, with a version of the package’s serial fitting function, niche_DE_no_parallel, that allows a separate *γ* for each cell type; with *γ* = 0.8 for every type, it reproduced the package’s own function exactly on one section (Supplementary Note 4). Calls came from get_niche_DE_genes at the interaction level with *α* = 0.05, on all genes. Types with more than 50 cells entered the model.

Labels were the truth labels, labels re-called on half A or on half B, the study’s labels, the synthetic published labels, or truth labels with injected errors. In a further analysis, the same labeler with centroids from the study’s labels re-called the labels on half A (Supplementary Note 4). All except the study’s labels came from nichepeek’s nearest-centroid labeler. For the truth labels, its centroids came from the full-depth counts and the study’s labels; for the re-called labels, as nichepeek runs on this file, from the experiment share and the synthetic published labels. Each set entered in the cell’s own role, in the neighbors’ role, or in both. A claim is a gene, an index type, a niche type, and a sign. A claim is robust when the truth-label fit calls it on both halves of the first split, with a second random seed, and on a second split. A reversal is a robust claim that a fit with other labels calls with the opposite sign on two splits; for the study’s labels, which were the same on every split, we required it on all three. A false claim is one that a fit with other labels calls on two splits and that no truth-label fit calls in either sign. Injected errors changed a fixed number of labels, a multiple of the size of the smaller type of the most confused pair. We placed them at random among fibroblast subtypes, as a swap within the pair, or as over-calling of its smaller member from other subtypes; the first two keep the composition. In the section with most reversals, we also injected the labeler’s own over-call of mIAFs alone, and the same number of errors moved to random cells of the same source subtypes or of any fibroblast subtype. We also placed the same number of errors per subtype as clumps of cells of that subtype around random seeds, with the clump size matched to the labeler’s patchiness, and we kept a random half of the labeler’s errors, either on its own cells or on nearby cells of the same subtype that it had labeled correctly. We described where errors lay by the share of mislabeled cells among the 20 nearest neighbors of each mislabeled cell. We wrote down the design of each run, dated, before the run. The analyses without Niche-DE fits differed. The checks of the class prior were exploratory. The comparison of labelers across datasets followed a rule written before the third dataset was built. The comparison with random training errors followed a rule dated before its run; we report its verdict, with a qualification in two datasets (Supplementary Note 4). The agreement filters followed a design dated before their run. We defined reversals and false claims after the first two runs and tested them on a further split with predictions written before it (Supplementary Note 4).

To compare labelers, we rebuilt the two labelers of the semi-synthetic data and checked that they reproduced the stored labels exactly. We then trained the same nearest-centroid labeler on each combination of counts (full depth or experiment share) and labels (the study’s or the synthetic published labels) and counted, per subtype, the cells it called on half A, against the truth. We did this in the colon data, in healthy liver (MERFISH), and in a further MERFISH dataset built for this test after its criteria were written down. To test agreement filters, we used the stored fits: claims of the re-called labels kept on one split, on that split and a new split with the same sign, or only where a fit with the synthetic published labels or with the study’s labels made the same claim, each scored against the true-label fits with the definitions above.

### 4.10 Magnitude check across populations, and neighborhood groups

nichepeek (nichepeek mcsimex --semisynthetic 0.5) builds the split of the previous sub-section on the fly (*f* = 0.5, *q* = 1; variants *f* = 0.75). Truth labels are called on *Y ^T^*, and published labels on *Y ^E^*, by the labeler of the main runs trained on the data’s own labels. The labeler that re-calls labels in the protocol is then trained on the synthetic published labels of *Y ^E^* (--semisynthetic-train synthetic, the default), which reproduces the run on the semi-synthetic files of the previous subsection exactly. As a design comparison, every run was repeated with the labeler that made the truth and the synthetic labels as the re-calling labeler (--semisynthetic-train reference). The variants used the logistic-regression labeler. The own-label arm runs on half *B* of *Y ^E^* with the arguments of each population’s main run. Synovium was analyzed in crops of 30,000 cells, two per section. The first-level estimate uses *p*_0_ = 0.8 and the native flip rate. The measured bias is *T*_1_ *−t*, as above, and ratios to it are ratios of medians over sections. Misclassification SIMEX [23] replaced the own labels by draws from Π*^λ^* for *λ* = 0.5, 1, 1.5, 2 (two draws each). Π is the native confusion matrix: the probability that a target with re-called label *a* is labeled *b* on a Poisson resample of half *A*. The neighborhood share was extrapolated to *λ* = *−*1 linearly and, for comparison, quadratically in *λ*. The oracle version used the confusion of re-called against true labels. Colon was also run with seeds 1 to 9 (seeds 1 to 4 in the design comparison) (--seed), which change the semi-synthetic split and the count splits, thinnings, and resamples of the protocol, but not the seed of the ladder’s grouping and permutations, which is fixed per section; with the run above and the file-based split of the previous subsection, these give 11 random splits at the calibration setting.

For the neighborhood groups, the targets of each section were divided into the 12 groups of the ladder (*k*-means on the compositions of published neighbor labels). The identity of group *g* under a label set is *n_g_*∥*µ̄_g_* − *ȳ*∥^2^, as a share of the section’s total sum of squares. Here *µ̄_g_* is the mean, over the group’s targets, of their label’s mean expression, and *ȳ* is the mean expression of all targets (size-normalized counts of half *B*). The loss is the identity with true labels minus the identity with re-called labels. The score is

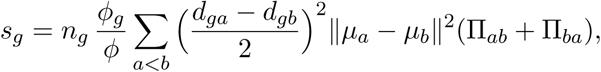

where *d_ga_* is the group’s share of re-called label *a* minus the section’s share, *µ_a_* is the mean expression of label *a*, and *ϕ_g_* and *ϕ* are the native flip rates of the group and of the section.

The score uses neither true labels nor the ladder. Composition imbalance is ∑_*a*_*d*^2^_*ga*_. Spearman correlations over the groups were computed per section and summarized by their medians.

### 4.11 Calibrated magnitudes

The real colon, healthy-liver, and MERFISH datasets were each also run at all three thinning levels 0.8, 0.65, and 0.5. The calibrated own-label bias is the first-level estimate at *p*_0_ = 0.8 divided by the estimate-to-measured ratio of the semi-synthetic data of the same tissue with the same first step (deep setting for colon and liver); for MERFISH, whose depth matches the shallow settings, it is divided by the shallow ratios of both tissues and reported as a range. Intervals come from a joint bootstrap: independent units (mice or donors) of the real data and of the semi-synthetic data were resampled separately (reported only where both sides have at least five units, i.e., colon; for liver we give the donor values), the ratio recomputed from medians over units, and for MERFISH one of the two shallow settings drawn at random in each replicate. Default-setting estimates are those of the main runs. The truth-corrected value divides by the corrected ratio of the deep setting. Its interval comes from the same joint bootstrap, with each semi-synthetic unit’s extrapolated bias in place of its measured bias. The calibration table packaged with nichepeek carries both ratios.

### 4.12 Use of large language models

The authors used Claude (Anthropic; several model versions, 2026) as an assistant throughout the study. It was used to draft and revise analysis code, the nichepeek package and its tests, and drafts of manuscript sections; to check the text for internal consistency against the computed numbers; and to propose analyses and critique drafts. All analyses were run by the authors on their own computers and on an institutional computing cluster. No number in this manuscript was produced by the language model (Software, code, and numbers). The authors reviewed, tested, and edited all code and text and take full responsibility for the content of this article.

### 4.13 Software, code, and numbers

We release the protocol as nichepeek [41]. A single call on an AnnData object runs count splitting with fixed targets and the three channels and returns per-section tables and a one-page report (Supplementary Fig. S3), which calls a direction only when at least three quarters of at least four sections agree. Further commands build the semi-synthetic check, give a calibrated magnitude, and rank neighborhood groups by their exposure to label error. Users can supply their own labeling function.

Ladder, channel, context, and niche-regression analyses were run with development versions of nichepeek (0.1.0.dev0 to 0.2.7). Release 0.3.5 reproduces their per-section results exactly, in every shared column of the per-section tables; we reran colon stroma and healthy liver to confirm it. The magnitude check and the neighborhood groups were run with nichepeek 0.3.3, whose computations are identical to those of the release. We produced the semi-synthetic data and simulations with our analysis scripts and kept dated notes of the analysis decisions. Every number in this manuscript is generated from the outputs of these runs by one script, inserted by placeholder, and every sign and order stated in the text is checked against the numbers at each build; the result tables and that script will be deposited on publication (Data availability). The two packages of this work have different jobs: nichepeek measures how label error biases estimates of neighborhood effects, and claimbreak [42] measures how segmentation changes the labels themselves [24].

## Data availability

This study reanalyzed only published data; ethics approvals and consent for the original human and animal studies are described in the original publications [10, 18–22]. All datasets analyzed in this study are publicly available: mouse colon stroma (Xenium) [18], Broad Single Cell Portal SCP3384; mouse colon during DSS colitis (MERFISH) [19, 37], Dryad, doi: 10.5061/dryad.rjdfn2zh3; healthy and fibrotic human liver (MERFISH) [20], Dryad, doi: 10.5061/dryad.37pvmcvsg (file adata_healthy_diseased_merfish.h5ad); rheumatoid arthritis synovium (Xenium) [21], Synapse, doi:10.7303/syn74734050.1; diabetic kidney disease (CosMx and Xenium) [22], Zenodo 19868428; non-small-cell lung cancer (Xenium) [10], GEO GSE311609, with the single-cell reference of Dong et al. [38] (ArrayExpress E-MTAB-14560 and E-MTAB-14566; CZ CELLxGENE, dataset 4ca422ae-e4ad-4884-9073-7c3799cd9143). The semi-synthetic data were built from the colon stroma (Xenium), healthy liver, and colon MERFISH data above with our analysis scripts (Code availability). The result tables behind every value in this article, the numbers file, the claim checks, and the script that recomputes every number and figure from the tables are available from the corresponding author on request and will be deposited at Zenodo on publication.

## Code availability

nichepeek is available at https://github.com/velgart/nichepeek (archived version nichepeek 0.3.5, doi:10.5281/zenodo.23987423). It runs on any platform with Python *≥* 3.10 and numpy, scipy, pandas, anndata, h5py, scikit-learn, and matplotlib, under the BSD 3-Clause license, with no restrictions on non-academic use. The analysis scripts for the semi-synthetic data, the simulations, and the Niche-DE fits are in Python and in R (R 4.6.1, with the Niche-DE build described in Methods; cluster jobs under Slurm). They are available to editors and reviewers at submission and from the authors on request.

## Acknowledgements

The authors wish to thank Rose Cannistraro for expert technical assistance.

## Author contributions

VE conceived the study, developed the method and the nichepeek software, performed the analyses, and wrote the manuscript. JL conceived and supervised the study, contributed to the interpretation of the results, and wrote the manuscript. Both authors read and approved the final manuscript.

## Supplementary Information

### Supplementary Note 1: the own-label channel in a two-type model

#### Setting

Consider targets of two true types with equal frequencies and mean expression *µ*_1_ and *µ*_2_ (the argument is gene-wise; the micro-average sums over genes). The ladder groups targets into neighborhood groups *g* at *S*_0_; in the own-label arm these groups are the same in every arm because targets are fixed and neighbors keep their published labels. Let *π_g_* be the share of type-1 targets in group *g*, and let expression be *y* = *µ_τ_* + *β_g_* + *e*, with a neighborhood signal *β_g_* that does not depend on the type *τ*. Each target’s own label is incorrect with probability *ε*, symmetrically, independently of its group and of *e*, given its true type.

#### Derivation

*S*_1_ permutes expression within own-label classes, such that the expected mean of group *g* at *S*_1_ is ∑*_ℓ_ π̃_ℓ|g_ m_ℓ_* up to a constant, where *π̃*_1_*_|g_* = *ε*+(1*−*2*ε*)*π_g_* is the share of the group’s targets whose *own* label is 1 (not the neighbor composition, which the own-label arm keeps fixed) and *m_ℓ_* is the mean expression of targets labeled *ℓ*. With equal frequencies, *m*_1_ *− m*_2_ = (1 *−* 2*ε*)(*µ*_1_ *− µ*_2_). The variance of the *S*_1_ group means is, therefore, (1 *−* 2*ε*)^2^ Var*_g_*(*π_g_*) (1 *−* 2*ε*)^2^(*µ*_1_ *− µ*_2_)^2^: the identity captured at *S*_1_ is attenuated twice, by (1 *−* 2*ε*)^4^ in all. *R*^2^(*S*_0_) does not involve own labels, such that

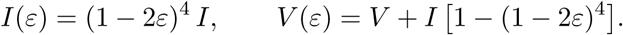

The own-label error moves identity into the neighborhood share; the inflation is non-negative, increasing, and concave in *ε*, and proportional to *I*, which is why it vanished where identity did (fibrotic liver). A Monte Carlo check with 12 groups and 48,000 cells reproduced the formula (neighborhood share 0.1009 against a predicted 0.1011 at *ε* = 0.1, and similarly from 0.05 to 0.3).

#### What the model does not give: the magnitude

Taken literally, the formula predicts far more inflation than we measured: in Xenium stroma (native flip rate 0.10, identity 0.169), it gives an own-label bias of about 0.10, larger than the whole published neighborhood share (0.065). Real errors are mostly swaps between similar subtypes. With many types and a confusion matrix, the attenuation of identity depends on the error rate weighted by the contrast between the types confused: a swap between near-identical subtypes removes little identity. The two-type model explains the sign and shape of the response; its magnitude depends on the confusion structure and the type contrasts, which thinning does not recover, which is why the magnitude has to be calibrated on data (Supplementary Note 3) rather than read off the model.

#### What the thinning response measures

Write *f* (*ε*) = *V* (*ε*) *− V*, with *f* (0) = 0. The protocol measures the chord of *f* from the native error *ε*_0_ to *ε*_0_ + *x_p_*, and the first-level estimate multiplies that chord by the native flip rate. For the two-type model, *f* is concave, such that a chord from *ε*_0_ is at most *f* (*ε*_0_)*/ε*_0_. If the flip rate did not exceed the error rate, the estimate would be a lower bound. Two conditions violate this assumption in real tissue. First, the flip rate understates the error rate, as it did in simulations (at most 0.93 of it) and on the semi-synthetic data (median 0.75), which lowers the estimate. Second, with many cell types, thinning does not add errors with the native confusion structure: added flips land on more dissimilar types (in 98 of 98 sections or crops of the real data), each removes more identity, and a long chord is steeper than the response that produced the native error, which raises the estimate. The two effects offset each other, in part, after a large first step in deep data and not otherwise, which is why the magnitude depends on the step and the depth (Supplementary Note 3, Table S9). With a small first step, the confusion structure of added flips stays close to the native one (the confusion patterns of added and native flips correlated at 0.84 or more at every level, and most closely at the first), the chord approaches the average response, and the estimate is low by about the flip-rate factor. This result holds against the truth labels as called. Against a truth corrected for its own errors, the estimate is lower still (Supplementary Note 3), and, thus, the flip-rate factor alone does not account for it.

#### Independence and the destination test

The derivation assumes that errors do not depend on the neighborhood given the true type. The occurrence test found little dependence of *which* cells change label on their neighborhood, but the destination test shows that *where* they land does: changes land on the label typical of the cell’s neighborhood more often than expected (Supplementary Note 2). When mislabeled cells take the label typical of their context, the own-label classes partly align with the neighborhood, *S*_1_ absorbs part of the neighborhood share, and the inflation is offset. This offset is the main mechanism by which real data depart from the model, and it is one way the own-label response can shrink where cell states vary with the neighborhood. In simulations, the response also turned negative where states varied independently of the neighborhood (Supplementary Note 2), which this argument does not explain.

### Supplementary Note 2: further results and checks

#### Simulations

Label error and circularity also moved identity, in opposite directions (label error *−*0.028, circularity +0.019). Circularity fell steeply with depth (Supplementary Fig. S6a), and at higher depth, it canceled a larger share of the label-error bias. A small net bias, therefore, does not show that labels are accurate.

When cell states varied continuously along the axis between types, label error still inflated the neighborhood share, whether the variation was independent of the neighborhood or followed it (Supplementary Fig. S6b). Only the net, same-count bias turned negative, in the most extreme neighborhood-following regime, and there circularity rather than label error made it so. Published-style labels, called and scored on all counts, lowered the neighborhood share against exact truth in that regime (*−*0.015; Supplementary Table S11).

With the channels separated and targets fixed, the native bias was almost entirely the own-label channel in every regime (Fig. 2b). The neighbor channel was small and positive. In these tissues, the non-target types that define the planted signal are well separated, and their labels are rarely incorrect. When we made them more difficult to distinguish with weaker marker genes, the neighbor channel turned negative (*−*0.020, negative in 6 of 6 sections, against +0.005 with the default markers), while the own-label channel changed little. Deflation through neighbors’ labels, as we find in real data, therefore, appeared only when the labels of the neighbor types were themselves error-prone, as is expected in panels where many types must be discriminated.

With re-selected targets, the thinning response had the opposite sign to the native bias whenever cell states formed a continuum (Fig. 2c). Fixing targets removes this reversal, but not every limitation. The own-label response had the sign of the native bias for discrete cell types. For continuous states, it vanished where states varied with the neighborhood and was negative where they varied independently of it, although label error inflated the neighborhood share in both regimes. A positive own-label response is, therefore, evidence of inflation, but a null or negative one is not evidence of its absence.

#### Seven populations

Supplementary Table S1. gives the channels of the seven populations, with intervals over independent units, and Supplementary Table S2 gives MERFISH colon by mouse. Within the synovial populations, the own-label channel grew with the published identity of each crop (Fig. 3b). This outcome is expected, since the own-label channel can only transfer variance that identity holds. In MERFISH colon, where target populations are small, the own-label channel was resolved at the mouse level, whereas the net effect was not (sign test over mice, *P* = 0.22). The populations differ in depth and were thinned to different levels (Table 2), such that magnitudes are not comparable across datasets, but signs are.

The membership channel was largest, and negative, where targets are easily confused with other cell types: in healthy liver, where hepatocytes are confused with non-hepatocytes at low depth, and in synovial myeloid cells. Re-selection swaps targets whose re-called label leaves the target set for non-targets that are called targets. Where the two are easily confused, the incoming cells carry expression unrelated to the target population’s neighborhood signal, which is a plausible reason as to why the swap lowered the estimate, although we did not isolate the mechanism. Because re-selection changes which cells are studied, *M* mixes a bias with a change of estimand. We, therefore, treat it as a design effect rather than as a label-error bias comparable with *T* and *N*.

In fibrotic liver, whose near-zero identity is consistent with the loss of zonation in fibrosis, the own-label channel was close to zero and the neighbor channel dominated, such that the net effect was negative. With little identity, there is little for the own-label channel to transfer. But the own-label response can also vanish where cell states form a continuum. Thus, a near-zero response alone does not show that the labels are accurate.

#### Samples from the same patients on two platforms

The populations above differ in tissue, platform, and labeling at once. To separate the platform, we used a kidney atlas in which samples from 4 patients (one with diabetic kidney disease, three controls) were imaged on both CosMx and Xenium [22]. We ran the protocol on four target sets: endothelial subtypes (glomerular, peritubular, and vasa recta), the stromal types (fibroblasts, vascular smooth muscle, and mesangial cells), and healthy and injured proximal tubule and thick ascending limb. A fifth set had one label for each of the two lineages, proximal tubule and thick ascending limb. Each ran on CosMx, on Xenium restricted to the genes shared with CosMx, and on the full Xenium panel (Supplementary Table S6). At the shared genes, the two platforms gave the model similar depth and similar label stability.

The own-label channel rose with identity across all runs (Supplementary Fig. S4b). It was positive on both platforms for 15 of 16 combinations of patient and one of the first four target sets. Per patient, the two platforms agreed only loosely in size (Supplementary Fig. S4a), and the size on Xenium was 0.4 to 2.3 times that on CosMx. Targets defined by a single label have no identity by construction, and in such runs, both identity and the own-label channel were at most 0.0010 in absolute value.

On the full Xenium panel, healthy and injured thick ascending limb were labeled very stably, and there the own-label response vanished. A null response carries no information about the bias where labels change this rarely, or where states form a continuum, as injury states do. The net effect there was nonetheless positive. Its largest component was the interaction between the two channels, which arises only when a cell and its neighbors are relabeled together; we did not resolve it further. Circularity was large in these data. Scored on the counts from which the labels were called, the neighborhood share fell by more than the own-label channel raised it in 9 of 12 runs. Thus, an analysis on the same counts could hide the inflation entirely. With one patient with disease, these data support no condition contrast.

#### Lung cancer: whole-cell against nucleus labels

Thinning adds errors that behave like counting noise; segmentation adds errors when transcripts of a neighbor fall inside a cell’s boundary. To compare the two, we labeled each cell of non-small-cell lung cancer sections (Xenium, three panels [10]) from its whole-cell and from its nucleus counts and used the whole-cell labels as published labels (Methods). We ran the protocol on two target sets, immune cells (T and NK cells, B cells, and myeloid cells) and stroma (fibroblasts and muscle cells, and vessels), in 48 crops of 4 donors (Supplementary Table S7). On the lung and immuno-oncology panels, where target identity was at least 0.02, the own-label channel was positive in 39 of 40 crops. On the largest panel, identity and the own-label channel were both close to zero, as in kidney below this identity.

The protocol’s response per added flip predicted what the nucleus labels did as the targets’ own labels. For each crop, we multiplied the slope of the own-label channel over the thinning levels by the fraction of targets on which the re-called and the nucleus labels disagreed. The median ratio of observed to predicted was 1.02, and within runs the two ranked crops alike (Supplementary Fig. S5a). Whatever its source, label disagreement in a cell’s own label moved variance from identity into the neighborhood share at the rate that thinning measured.

In the neighbor role, the nucleus labels lowered the neighborhood share in 4 of 4 runs, by more than thinning predicted in 4, and in 1 of them the prediction had the opposite sign (Supplementary Fig. S5b). With nucleus labels in both roles, the neighborhood share was lower than with the re-called labels in 4 of 4 runs, whereas the protocol’s net effect was positive in 4. Where segmentation dominates label error, the neighbor channel, and with it the net effect, can, therefore, be larger than the protocol reports. Nucleus and whole-cell labels disagree mostly on cells in contact with other types [24], whereas thinning places flips as counting noise would.

#### The standard design in every population

Besides the populations named in the main text, the standard design reported +0.011 in healthy liver, against +0.030 with the protocol. Both distortions disappear when targets are fixed and counts split. With fixed targets, circularity barely reached *R*^2^(*S*_0_) (for example +0.0005 in MERFISH).

#### Condition contrasts

For each contrast, we multiplied each condition’s own-label sensitivity by its native flip rate and took the difference. This difference gives a first-order estimate of how much of the contrast a difference in label error could produce, and is a sensitivity rather than a bound (Methods; Supplementary Table S3). In mouse colon stroma, labels were less stable after DSS than in water-treated mice at equal depth, and this difference alone corresponded to a difference in the neighborhood share nearly as large as the observed contrast, which the interval over mice did not resolve. In MERFISH colon, the label-error term was small beside the contrast with either labeler, with two mice per condition. Between healthy and fibrotic liver, the label-error term was not negligible beside the contrast (three donors per condition and, therefore, no interval). The contrasts here are in our own estimate of the neighborhood share rather than findings of the original studies, and the label-error term is a first-order approximation.

In colon stroma, the first-level estimate of the own-label bias (Supplementary Note 3) was larger in every DSS-treated mouse than in any water-treated one (Mann–Whitney *P* = 0.002, the smallest attainable with six mice per group). Because the estimate is the response per added flip times the flip rate, part of this difference is built in: the flip rate was 1.26-fold higher after DSS, and the response per added flip only 1.11-fold; thus, the difference came mainly through label stability. The difference did not follow identity, which was the same in both groups. Consistently, the truth labels of the semi-synthetic data agreed with the published labels less often in DSS sections than in water-treated ones: inflamed tissue is harder to label.

The difference did not come from the labeler’s class prior. Both labelers have flat priors, which can over-call rare types and subtypes, and in colon stroma several targets are rare. With labelers whose priors match the published label frequencies (Supplementary Table S14), the estimate remained larger after DSS, and labels stayed less stable after DSS. The separation of the estimates was no longer complete: 2 and 3 of the 36 pairs of a DSS-treated and a water-treated mouse overlapped. The contrast is, therefore, robust to the class prior, but its complete separation is specific to the nearest-centroid labeler.

#### Occurrence and destination of label changes

Across all populations, the neighborhood barely predicted which cells changed label. The gain in held-out AUC over the cell’s own label and depth was +0.0004 in Xenium, +0.014 in MERFISH, and +0.003 in synovial fibroblasts. Where cells went, however, followed the context. Among cells whose re-called label agreed with the published one, 0.33 of Xenium label changes landed on the label typical of the cell’s neighborhood against 0.24 for the same transitions in random cells. In healthy liver, the shares were 0.16 and 0.08 (Supplementary Fig. S2a). A cell’s expression carries the neighborhood signal that the analysis measures as the neighborhood share, and label error moves that signal into the label, i.e., the own-label mechanism seen from the other perspective. The destination result has a second explanation. Segmentation spillover adds neighbors’ transcripts to a cell and pulls its label toward the neighborhood’s typical type. Protein staining has shown such spillover-driven label swaps directly [10]. In our simulations, 20% spillover raised the share of a mislabeled cell’s true neighbors that carried its incorrect label from 0.064 to 0.084. The destination test shows that label changes follow context, but not which of the two mechanisms moves them.

#### A standard niche regression

We also scored every arm with a niche-regression model of the NCEM type [3]. It regresses expression on the cell’s own label and its neighbor composition, with cross-validation, and defines the niche effect as the gain from adding the neighbors. The own-label channel inflated the niche effect in all seven populations, and the neighbor channel deflated it in all seven (Supplementary Fig. S2c). Relative to the ladder, the regression was more exposed to the neighbor channel in five of the six populations with a measurable own-label channel (all but healthy liver). This outcome is expected when neighbor compositions enter directly as regressors because label noise in them attenuates their coefficients. The net sign, therefore, depended on the dataset. It was positive in the three synovial populations, Xenium stroma, and healthy liver, and negative in fibrotic liver and in MERFISH colon, where many fine labels make neighbor labels error-prone. With labels called on the same counts, the noise-corrected type effect of the regression exceeded the total biological variance at low depth, which shows directly that same-count labels fit sampling noise.

#### Labeler dependence

We repeated the protocol for all seven populations with a second labeler of a different kind: multinomial logistic regression trained on thinned counts (Methods; Supplementary Table S8). Its labels were less stable in every population. No population-level sign changed: the own-label channel was positive and the neighbor channel negative in every population with non-negligible identity. The number of units with a positive net effect depended on the labeler in three populations: healthy liver, synovial myeloid cells, and synovial fibroblasts. In the DSS–water contrast, the label-error term with the second labeler was again as large as the observed contrast. On the semi-synthetic data, the estimate with the logistic-regression labeler had the same sign as, and a somewhat smaller overstatement than, the estimate with the labeler that made the labels (Supplementary Table S9).

Both labelers have flat class priors: the nearest-centroid labeler has none, and the logistic regression was trained on at most 400 cells per label. Flat priors can over-call rare types and subtypes. Abundant classes also dominate standard embeddings [40]. In colon stroma, we, therefore, added two labelers with natural priors (Methods): the nearest-centroid correlations turned into a posterior with the published label frequencies as prior, and logistic regression trained on a uniform sample of cells (Supplementary Table S14). Their labels agreed more often with the published ones and were more stable. The own-label channel kept its sign and changed little in size, and label changes still landed on context-typical labels. Lower flip rates and a steeper response per added flip offset each other in the first-level estimate of the nearest-centroid labeler. We did not repeat this analysis for the other populations or the calibration.

#### Analysis settings and technical noise

Varying the neighborhood size and the number of groups, or omitting the clipping of negative variance estimates, did not change the sign of the own-label channel (Supplementary Table S5). The small neighbor channel of colon stroma turned slightly positive with *k* = 40 and with six groups, and the small MERFISH net effect changed sign with *k* = 10 (*−*0.002). Across these settings, the own-label response at the middle thinning level ranged from 0.34 to 0.64 of the neighborhood share in colon stroma and from 0.40 to 0.48 in healthy liver. Below about ten counts per cell, the channels could change sign, for which reason such levels are shown but not used. Labels called on half of the counts are less stable: on resampling, 0.10 of target labels changed at half depth in colon stroma, against 0.07 at full depth, and 0.08 against 0.07 in healthy liver.

Count splitting assumes Poisson-like technical noise. We, therefore, simulated overdispersed technical noise (gamma–Poisson, squared coefficient of variation 0.3). The circularity left after splitting, measured against an independent replicate, was *−*0.003 (Supplementary Table S4). Overdispersion diluted all variance shares (true neighborhood share 0.069 without it, 0.023 with it) but did not change the signs of the channels. Segmentation spillover, which both halves share, raised the apparent neighborhood share with true labels from 0.069 to 0.081 at 20% spillover, and the own-label channel persisted (+0.042).

Section-level estimates are noisy where target populations are small. In MERFISH colon, the split-to-split standard deviation of the net effect (0.014) was larger than the effect itself. We, therefore, averaged 3 independent splits and 20 permutations at the *S*_1_ rung (a single permutation in the kidney and lung runs); the package defaults are a single split and a single permutation; both were, therefore, set explicitly. nichepeek reports the spread between splits and summarizes datasets at the level of independent units (mice, donors, or patients) rather than sections or crops.

### Supplementary Note 3: the size of the bias

This note gives the calibration of the own-label estimate and the checks of its size. The values describe the bias of the labels that the protocol re-calls, measured against truth labels that carry errors of their own.

#### Semi-synthetic data

The published labels cannot serve as truth because they were called from the same counts as the data, and a classifier differs from them systematically. We, therefore, built semi-synthetic data for colon stroma (Xenium) and healthy liver (MERFISH) with truth and experiment of equal depth, with a shallower experiment, and with the same shallow experiment against a deeper truth (Methods).

The truth labels were stable on resampling, yet they agreed with the real published labels for a smaller share of targets (Supplementary Table S15). Most of the disagreement between a classifier and published labels is, therefore, systematic. The synthetic published labels carried a label error in the range of the real data.

The sign result also held when labels were re-called by the logistic-regression labeler, which differs from the classifier that made the synthetic published labels (Fig. 4b; the 96 unit-settings share units and splits and are not independent). The measured bias was large beside the neighborhood share: 22% of it in colon at the deeper setting and 59% at the shallower. These shares describe the construction rather than the real tissue. The synthetic published labels were called at lower depth than the real published labels and carry correspondingly more noise-driven error. In the two shallow settings, the deeper truth gave the larger measured bias, as expected if errors in the truth labels shrink the measured bias. The calibration below corrects for this effect.

The size of the first-level estimate depended on depth and on the first thinning step (Supplementary Fig. S7a; Supplementary Table S9). This estimate is the own-label response per added flip at the first thinning level times the native flip rate. With the default step (*p* = 0.5), it overstated the bias in the deeper setting and understated it in the shallower.

The two factors of the estimate behaved differently (Supplementary Fig. S7b). The native flip rate understated the true error rate by a median factor of 0.75 (0.48 to 0.87 over independent units and settings). The response per added flip overstated the bias per unit of error after a large first step in deep data, and much less after a small step (*p* = 0.8). The reason for this outcome is that thinning adds flips that land on more dissimilar types than native flips do: the dissimilarity of added flips rose with thinning in 98 of 98 sections or crops across all seven populations. A long step, therefore, measures a steeper response than the one that produced the native error.

With a small first step, the estimate understated the bias in deep data, and more so in shallow data and against a truth corrected for its own errors (Supplementary Table S9). Across the units of one setting, the estimate ranked the bias only moderately: the Spearman correlation across individual mice in the colon data at the deeper setting was 0.46. Magnitudes, therefore, describe datasets rather than individual mice. The net estimate was less reliable than the own-label estimate at low depth, and the measured neighbor-channel bias had opposite signs in the two tissues. We, therefore, report the neighbor channel as a sensitivity rather than a validated bias.

#### Calibration in colon stroma and healthy liver

A size needs a calibration made with the same labeler on semi-synthetic data from the same tissue, and colon stroma is the worked example. We re-estimated the two deepest datasets with a first step of *p* = 0.8 and divided by the factor measured on semi-synthetic data from the same tissue (Methods; Supplementary Table S16). The values refer to the neighborhood share *V* of the ladder. The niche regression responds with the same signs (Supplementary Note 2), but whether the size of the bias transfers to other estimators of neighborhood effects is untested.

In colon stroma, errors in cells’ own labels accounted for 10% of the estimated neighborhood share. Its interval covers the sampling of mice only. The divisor also varies with the random split that builds the truth: over 11 random splits of the colon data at this setting, the uncorrected share lay between 9% and 11% (Supplementary Table S10).

The calibration treats the truth labels of the semi-synthetic data as correct, but they are not. Their own errors inflate the neighborhood share scored with the true labels. The measured bias is, therefore, too small, and the divisor too large. We estimated this effect by calling the truth labels again on thinned truth shares and extrapolating the measured bias to a truth flip rate of zero under a reliability rule (Methods). In colon, the correction raised the share to 23%. Both colon intervals cover the sampling of units only. They do not include the uncertainty of the extrapolations in truth error and in depth. In liver, thinning barely raised the truth flip rate, which is too small a range for a reliable extrapolation; liver data, therefore, remain uncorrected. Extrapolated nonetheless over about 10 times the observed range of truth flip rates, the liver values serve only as a sensitivity analysis.

The calibration also extrapolates in depth. The real colon data give the ladder twice the depth of the deepest semi-synthetic setting. After the truth correction, the ratio of estimate to measured bias was about the same at two depths (0.29 and 0.29) and lower in a further, deeper setting (0.22), such that depth gives no bound on the calibrated value in either direction. With the default thinning levels, the first-level estimate was 1.3- and 1.4-fold larger, as the semi-synthetic data predict for a large first step in deep data.

For MERFISH colon, the calibration does not transfer. MERFISH colon matches the depth of the shallow semi-synthetic settings, but its labels are less stable than in any semi-synthetic setting, and the first-level estimate at *p* = 0.8 was positive in only 13 of 16 sections. Divided by the shallow ratios of the two tissues, the estimate gives a very wide interval (Supplementary Table S16). We, therefore, report no calibrated magnitude for MERFISH. Label stability and depth vary together across the datasets, such that a statement that the share is larger where labels are less stable would be a joint statement about both.

#### Published labels

The calibration describes the labels that the protocol re-calls. Published labels are called and scored on all of the counts, such that they carry label error and circularity together. In simulations with exact truth, such labels inflated the neighborhood share by about 13% of its true value (Supplementary Table S11). The inflation held where subtypes were confusable, shrank with depth, and was about zero where subtypes separated well. It reversed where cell states formed a continuum that followed the neighborhood (Supplementary Note 2). Semi-synthetic data cannot measure this bias. Their truth labels come from the same classifier, at the same depth, as the published labels, such that the label errors of the two cancel. Scored against such truth, the published labels lowered the neighborhood share in all eleven runs in which the truth and experiment shares were equal (Supplementary Table S12), as circularity alone would.

#### Niche geometry

We varied the niches of synthetic tissues (Methods; Supplementary Fig. S1). The own-label bias grew with niche size and tracked the identity that the neighborhood carries, staying between 0.44 and 0.60 of the true identity at every size and in each of 3 independent tissues. Confusable subtypes that share niches produced no bias at the same error rate. With counting noise alone, errors were no more frequent at niche boundaries than inside niches. Segmentation-like spillover made cells next to their confusable partner type more likely to take its label. The bias is, therefore, largest where confusable subtypes occupy separate niches that are large relative to the neighborhood the analysis uses, and small where they intermix.

#### Magnitude check across populations

The calibration above needs semi-synthetic data from the same tissue at a similar depth. To see how the uncalibrated estimate behaves in other tissues, we built semi-synthetic data for six populations with the arguments of their main runs (Methods; Supplementary Table S10). As in the calibration, the labeler that re-calls labels learned the synthetic published labels, as it learns the published labels of real data. With the nearest-centroid labeler, the default, the first-level estimate fell below the measured own-label bias in 47 of 51 units (mice, donors, or patients, counted per population; Fig. 4c). Healthy liver was the exception: there the estimate matched the measured bias.

The shortfall grew as labels became less stable (Spearman correlation between the ratio and the flip rate over runs, *−*0.80). In each of the three populations with a smaller experiment share, the flip rate rose and the ratio fell. Depth and stability changed together in these runs.

The shortfall depends on the labeler. With logistic regression, the estimate was 0.91 times the measured bias in colon stroma, 1.49 times in synovial fibroblasts, and 1.66 times in synovial myeloid cells. The difference lies in the measured bias rather than in the estimate. We repeated every run with the labeler that made the truth and the synthetic labels as the re-calling labeler, so that re-called and synthetic labels differ only by counting noise. The estimates barely changed. When the labeler learned the synthetic labels, the measured bias was larger in 7 of 9 runs with the nearest-centroid labeler and smaller in 3 of 3 with logistic regression. Thinning measures the response to errors that behave like counting noise. Systematic disagreement between the labeler and the labels it learned adds a part that thinning does not see, and that part raised the bias with one labeler and lowered it with the other.

Misclassification SIMEX [23] adds errors from the estimated confusion matrix and extrapolates them back to none. It did not do better (Supplementary Fig. S7c), and the true confusion matrix gave a similar range. Thus, the error lies in the extrapolation rather than in the estimated confusions. Quadratic extrapolation, the other common choice, overstated the measured bias in 7 of 9 runs. A calibration, therefore, transfers only for the same labeler and between data of similar label stability. For another labeler, the size of the bias comes from the same check run with that labeler on the user’s own data: nichepeek mcsimex --semisynthetic builds it, and the Python interface accepts the user’s labeling function.

#### Neighborhood groups

For each of the 12 neighborhood groups of the ladder, we measured the identity lost between true and re-called labels and asked whether the re-called labels alone could predict it. A score that applies the two-type formula to each pair of confusable labels did so (Methods; Fig. 5c). With the nearest-centroid labeler, it ranked the groups by their loss with a median Spearman correlation of 0.68 to 0.91 in colon stroma and in synovial fibroblasts and myeloid cells, and less well in synovial endothelium, MERFISH colon, and liver. The imbalance of label composition ranked the groups as well as the score did over 11 random splits of the colon data. Only in liver was the score clearly better (0.46 against 0.06). Group size did not predict the loss. The 3 groups with the highest score held a median of 24% of the targets and 56% of the loss. Ranked by the imbalance of label composition instead, the top groups held 21% of the targets and 57% of the loss.

### Supplementary Note 4: Niche-DE on semi-synthetic data

#### Design

Methods (Section 4.9) give the data, the settings of Niche-DE, and the definitions of claims, reversals, and false claims. Each fit used one set of labels for the cell’s own role and one for the neighbors’ role. The reference fit used the truth labels in both roles on half B. Three fits measured noise without label error: the truth labels on half A (counting noise), the truth labels on half B with a second random seed of Niche-DE, and the truth labels on half B thinned to half and to a quarter of its counts (depth). The re-called labels entered in the cell’s own role, in the neighbors’ role, or in both, and in both roles as labels re-called on half B, the counts that the model saw. We wrote a dated design note before every run. The first three runs also carried predictions with decision rules (Supplementary Table S17), and we report each of them, including those not supported. We defined reversals and false claims after the first two runs and confirmed them on a third split with predictions written before it. The later runs had design notes with readouts but no predictions.

#### Turnover, depth, and the noise floor

Turnover is the number of calls that a fit made or lost against the reference, divided by the calls of the reference, on triples that both fits tested. Counting noise alone changed at least 0.40 of the calls in every section of the first run, with a median of 0.63, and label error changed a similar share in most of them (Supplementary Table S17, N1 and N2; Supplementary Fig. S8). Turnover from counting noise rose as the depth fell (N5). Niche-DE’s own randomness was larger than we had predicted: a second seed changed 0.09 to 0.15 of the calls (N6, not supported), although effect sizes ranked almost alike (median Spearman correlation 0.99). The excess of label error over counting noise did not stay the same on a new split (N4, not supported). In a post-hoc check, the number of true-label calls itself changed between splits by up to 3.1 times in one section, which made the counting-noise turnover unstable; the label-error turnover changed by at most 0.13 between splits. The inflamed section was the exception to N2: label error changed 0.92 of its calls, against 0.49 for counting noise, and at most 0.72 in any other section. The pilot section entered the later analyses but not the calls of the first run, which counted eleven sections.

#### Reversals and their confirmation

Counting noise and the seed reversed almost no call of the reference fit in any section, whereas the re-called labels reversed many in the inflamed section. We, therefore, defined a strict reversal: a claim that the reference made on both halves of the first split, with the second seed, and on the second split, and that a fit with re-called labels reversed on both splits. Label error made 75 strict reversals in seven sections. On a third split, all five predictions held (Supplementary Table S17, C1 to C5): most reversals in the inflamed section recurred, counting noise and the seed again reversed almost nothing, and false claims of label error recurred more often than those of counting noise. Outside the inflamed section, 10 of 20 reversals recurred. The three splits share sections, cells, and truth labels; they test reproducibility across draws of the counts and of the re-called labels rather than across tissue.

#### Epithelial genes

Of the 75 strict reversals, 37 concerned epithelial-type genes measured in fibroblasts, from a list fixed before the split-2 run (Epcam, Krt8, Krt18, Car1, Car2, Car4, Hmgcs2, Slc26a3, Slc6a14, Fabp2, Cyp2c55, Gpx2, S100a14, Saa1, Ly6i, Sult1a1, Xbp1, Stard10, and Tmem176a). Transcripts donated by neighbors create such associations even with correct labels [24], such that these claims are technical from the start; label error changed their sign. The reversals of fibroblast genes, among them Col1a1, Col18a1, Wnt5a, Grem1, Ptgs2, and Tgfbi, also recurred on the third split (C3).

#### Published labels

Two published annotations exist for these data. The synthetic published labels were called by a labeler of the same kind on the whole experiment share, as a study’s own labels would be called, and the labeler of the re-called labels learned its subtypes from them. The study’s labels are the annotation of the original study. Both were called on counts that include half B, which the model saw, and both are the same on every split. The synthetic published labels reversed no claim (R1, not supported), and their false claims exceeded twice those of counting noise in only three of twelve sections (F1, not supported); false claims did not rise with their disagreement from the truth (D1, not supported). The study’s labels held few cells of mIAFs: a check written into an amendment before the run found enough of them to fit Niche-DE in only three sections; we, therefore, reported that arm without a decision rule. In those sections, the study’s labels reversed 26 claims, in every one of them (R2), and counting noise reversed none. Most of these had mIAFs as the niche type. The study’s labels called mIAFs on at most 0.08 times as many cells as the truth held, and the re-called labels on 1.5 to 3.9 times as many: the two annotations draw the boundary of this subtype in opposite directions. The truth class is large because the truth labeler had a flat prior (Labelers, below). The result, therefore, shows that two annotations that differ systematically in one subtype reverse niche claims; it does not show an error in the original study.

#### Injected errors

We injected errors into the truth labels of four sections, in numbers that were multiples of the size of the smaller type of the most confused pair (Supplementary Table S18). Errors placed at random among fibroblast subtypes kept the composition and reversed no claim at any dose, although they made false claims. Swaps within the pair and over-calling of the smaller type reversed claims, more at higher doses, and nearly always with that type as the index type. Over-calling through the cell’s own label reversed claims; the same errors in the neighbors’ labels reversed almost none, even when we inflated mIAFs nine-fold at random cells, such that most cells labeled mIAFs belonged to other types.

#### The labeler’s own over-call

In five sections, we replaced the truth label with mIAFs on every cell that the labeler called mIAFs, and kept all other truth labels. In the inflamed section, the over-call in the neighbors’ role reversed 33 claims, 29 of them among the 36 of the full re-called labels in that role. In the cell’s own role, it reversed 5 (4 of them among the 9 of the full labels), and in both roles 19 (16 of 26). In the other four sections, the over-call reversed none, while the full re-called labels reversed seven through the neighbors’ role; those reversals came from other errors of the labeler.

#### Placement in the inflamed section

We moved the labeler’s over-call of mIAFs, keeping the number of errors per source subtype, to random cells of the same subtypes, to random fibroblast cells, and to clumps of cells of the same subtype around random seeds, with the clump size set to match the labeler’s patchiness. We also kept a random half of its errors, either on the labeler’s own cells or on nearby cells of the same subtype that it had labeled correctly (Supplementary Table S19). We describe each placement by three measures: patchiness, the share of mislabeled cells among the 20 nearest neighbors of a mislabeled cell; the share of true mIAFs among those neighbors; and a state score, how far the expression of the changed cells on half B leaned toward mIAFs, above the mean of cells of the same true subtypes. Moving the errors within their subtypes changed the share of true mIAFs among their neighbors little, yet removed every reversal through the neighbors’ role; proximity to true mIAFs does not explain the reversals. Clumps matched the labeler’s patchiness and reversed few claims. Half of the labeler’s errors reversed 5 claims against 33 for all of them; the effect of dose was, therefore, not linear, and reversals could not separate the labeler’s cells from their places, as the amendment to the design had foreseen. In a post-hoc comparison of false claims, half of the labeler’s errors reproduced 138 of the false claims of all of them, and the same number of nearby other cells reproduced 42; the first set is part of the full one, such that some overlap was certain. Two limits apply: patchiness was matched at the scale of 20 neighbors only, and the nearby cells kept the places of the labeler’s errors approximately and part of their state score.

#### Labelers

The truth labels and the re-called labels came from the same nearest-centroid labeler with a flat prior and different centroids: from the full-depth counts and the study’s labels for the truth, and from the experiment share and the synthetic published labels for the re-called labels. We rebuilt both and checked that they reproduced the stored labels exactly in every section, after checking that the source file was unchanged and that the random stream regenerated the split. On half A, with a flat prior, centroids from the study’s labels called mIAFs on 0.96 to 1.51 times the truth’s cells with full-depth counts and 1.08 to 1.76 with the experiment share; centroids from the synthetic published labels called 1.51 to 3.74 and 1.52 to 3.88 times (the last are the re-called labels). With the study’s label frequencies as prior, the labeler with study-label centroids called no cell mIAFs, at either depth; with a prior shrunk halfway toward flat, it called 2218 cells mIAFs, against 12,342 in the truth, over all sections. These checks were exploratory, run without a decision rule.

#### Labelers across datasets

We repeated the comparison of labelers in healthy liver (MER-FISH) and in a further dataset of mouse colon (MERFISH), built for this test after its rule was written down (Supplementary Table S20). For each label, the chain is the ratio of the cells called on half A by the labeler trained on the synthetic published labels to those called by the labeler trained on the study’s labels, both on the experiment share. The chain fell with the truth share of the label in colon and in the further MERFISH data, and tended to in liver: labelers that learned from re-called labels inflated rare labels more than frequent ones. The rule required a negative correlation, a chain below one for the most frequent label, and at least one label inflated by more than 30%, all of them with less than 5% of the cells. It held in the further dataset and, applied after the fact, in colon; it failed in liver, where the chain of the most frequent label was just above one. Depth alone changed little: with the study’s labels, centroids from the experiment share and from the full counts called about the same numbers of cells. Whether a label lay between others in expression did not predict its chain.

#### Random errors in the training labels

Any noise in training labels can inflate rare subtypes: a few cells of a frequent subtype make up a large share of a rare one. To test whether learning from re-called labels matters beyond such noise, we trained two null labelers like the one that learned from the study’s labels, on the experiment share, after changing each study label with the probability at which the synthetic published labels differ from it on that label. The new label was drawn either uniformly or in proportion to subtype frequency. We compared each labeler’s calls on half A with those of the labeler trained on the study’s labels, per subtype, and summed the excess over one. A rule dated before the run called the result chain-specific in a dataset if both nulls had less than half the excess of the re-called labels, and generic if a null had at least half the excess and a negative Spearman correlation with log truth share. In the colon data, the result was chain-specific: neither null inflated mIAFs (0.87 at most, against 1.81 for the re-called labels). In liver and in the further MERFISH data, the rule called the result generic. Its excess counted any distortion, and the nulls distorted subtype frequencies there by more than half as much as the re-called labels; its sign criterion accepted any negative correlation, and no null correlated significantly negatively with log share in either dataset. The nulls, therefore, did not favor rare subtypes, which is what the rule was meant to detect. The colon result bears on the semi-synthetic Niche-DE analysis. For the other two datasets, we report the rule’s verdict, generic, with this qualification.

#### A labeler trained on the study’s labels

To ask whether the reversals came from the chain of two labelers, we re-called labels on half A of both splits with the same labeler, prior, and counts, but with centroids from the study’s labels, and fitted Niche-DE with them in each role in the inflamed section and the four sections of the titration (Supplementary Table S22). A design note with a reading rule was dated before the run; an amendment, written before any label was saved, built the labeler with the random seed of each split’s export, as the re-called labels had been built. The new labels agreed with the truth on 0.80 to 0.85 of the cells, against 0.68 to 0.83 for the re-called labels. They still over-called mIAFs, but less: in the inflamed section, they called 1.1 times the truth’s cells, against 3.9 times, and only 0.55 of the cells that they called mIAFs were mIAFs. In the five sections, they made one strict reversal, against 61 distinct claims for the re-called labels in the same roles, some of them reversed in more than one role (Supplementary Table S22 counts them per role), and none in common with them. Their persistent false claims were 0.50 times those of the re-called labels in both roles and 0.38 times in the neighbors’ role, and in the inflamed section still more than counting noise made (196 against 74). Effect sizes ranked closer to those of the true-label fit. By the rule written before the run, the labeler trained on the study’s labels removed most label-error reversals. The truth labeler also learned from the study’s labels, at full depth, such that the two labelers differ only in the depth of their centroids and in the counts they label, and the new labels carry mostly counting noise.

#### Labels from the model’s half

Labels re-called on half B, the half the model saw, in both roles made 31 strict reversals and 1284 persistent false claims over all sections, against 35 and 1303 for labels re-called on half A. Splitting the counts removes circularity from our measure (Section 2.3), but for these calls it changed little, because the labeler’s systematic errors are the same in both halves.

#### Agreement filters

We scored filters on the stored fits against the true-label fits, in every section (Supplementary Table S21). A refit on a new split keeps by definition every reversal and false claim that label error made on both splits, such that it can remove only those made once. Agreement with the synthetic published labels removes every reversal by construction, because those labels reversed none and the labeler learned from them. Agreement with the study’s labels is the informative filter, and it exists in three sections only; those labels share counts with the model’s half. The row with truth labels is the reference: robust claims are defined from the truth fits, such that it keeps all of them by definition. The row for the single fit in the three sections gives the study’s labels their baseline.

#### Software and compute

Niche-DE was the development version 0.0.0.9000 from GitHub (kaishumason/NicheDE, commit 87e0e89), with Seurat removed from the declared dependencies; the patch is archived with the scripts. The same build ran on the computer of one author and on the cluster, where its functions were checked to be identical. Each fit ran on one core; in the main run, the median fit took 9 minutes, and its 228 fits took 37 core-hours in all. On one section, with true labels in both roles, a fit with the package’s own niche_DE_no_parallel reproduced the stored fit exactly, calls and per-kernel statistics alike. The package’s parallel function, niche_DE, fits the same model gene by gene; we did not run it.

## Supplementary figures

**Figure S1:**
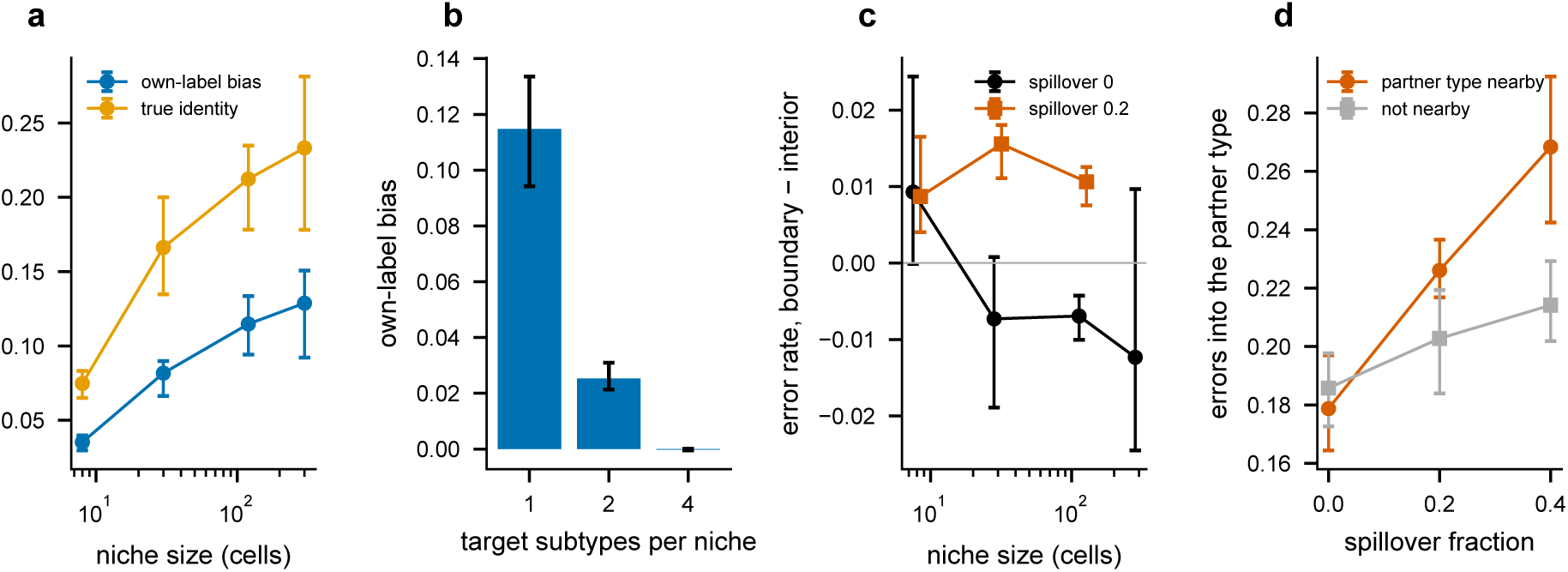
What controls the size of the bias. (synthetic tissues with controlled niches; means and ranges over independent tissues). **a**, Own-label bias and true identity against niche size, one target subtype per niche, no spillover. **b**, Own-label bias against the number of target subtypes sharing a niche (niches of 120 cells). **c**, Error rate at niche boundaries minus inside niches, without and with 20% spillover. **d**, Share of target cells labeled as their confusable partner type, with and without that type among their six nearest neighbors, against spillover.

**Figure S2:**
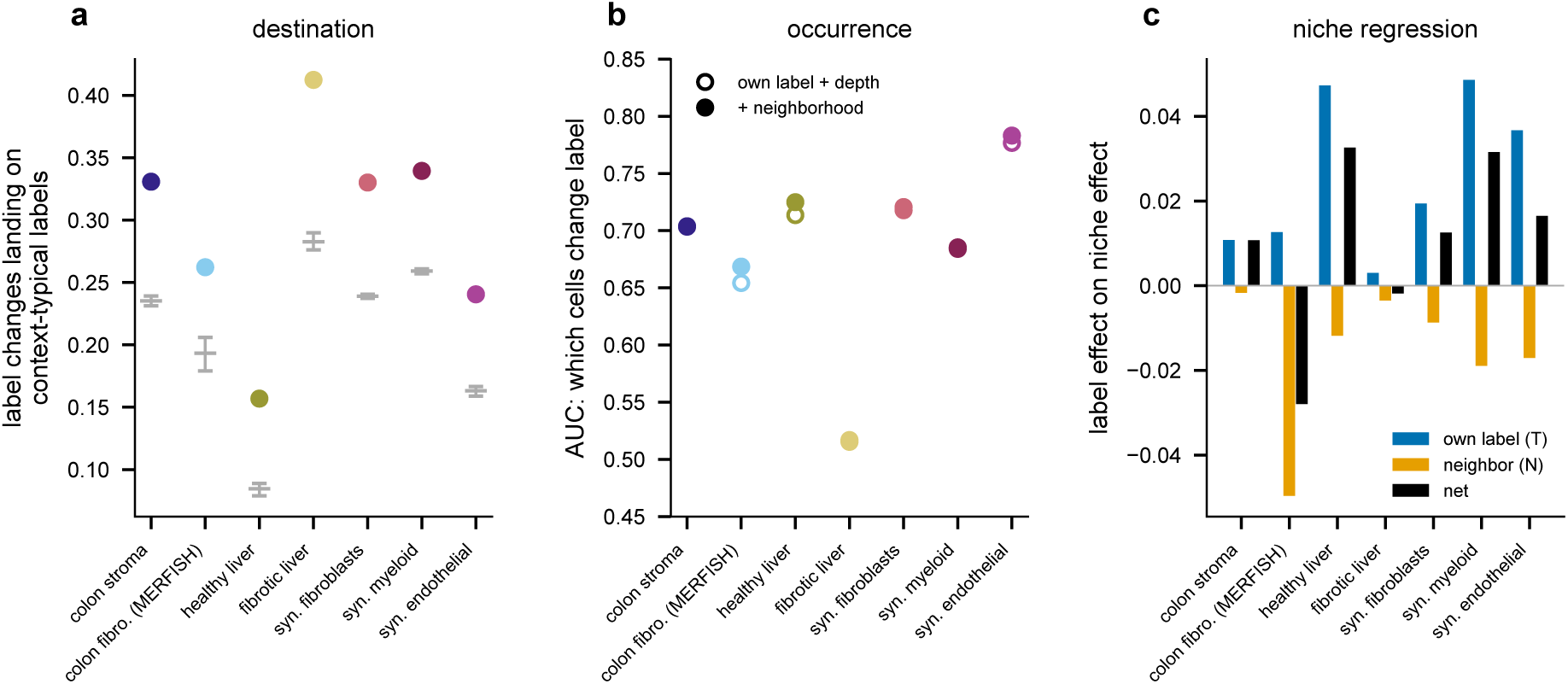
Context tests and a standard niche model. **a**, Fraction of label changes landing on the label typical of the cell’s neighborhood (dots), among cells whose re-called label agrees with the published one, against the same transitions assigned to random cells with the same label (gray, 95% interval). **b**, Held-out AUC for predicting which cells change label from the cell’s own label and depth (open) and with the neighborhood composition added (filled). **c**, NCEM-style niche regression: label effect on the niche effect through the own-label and neighbor channels, and net. All panels at the middle thinning level. Population names as in Fig. 3.

**Figure S3:**
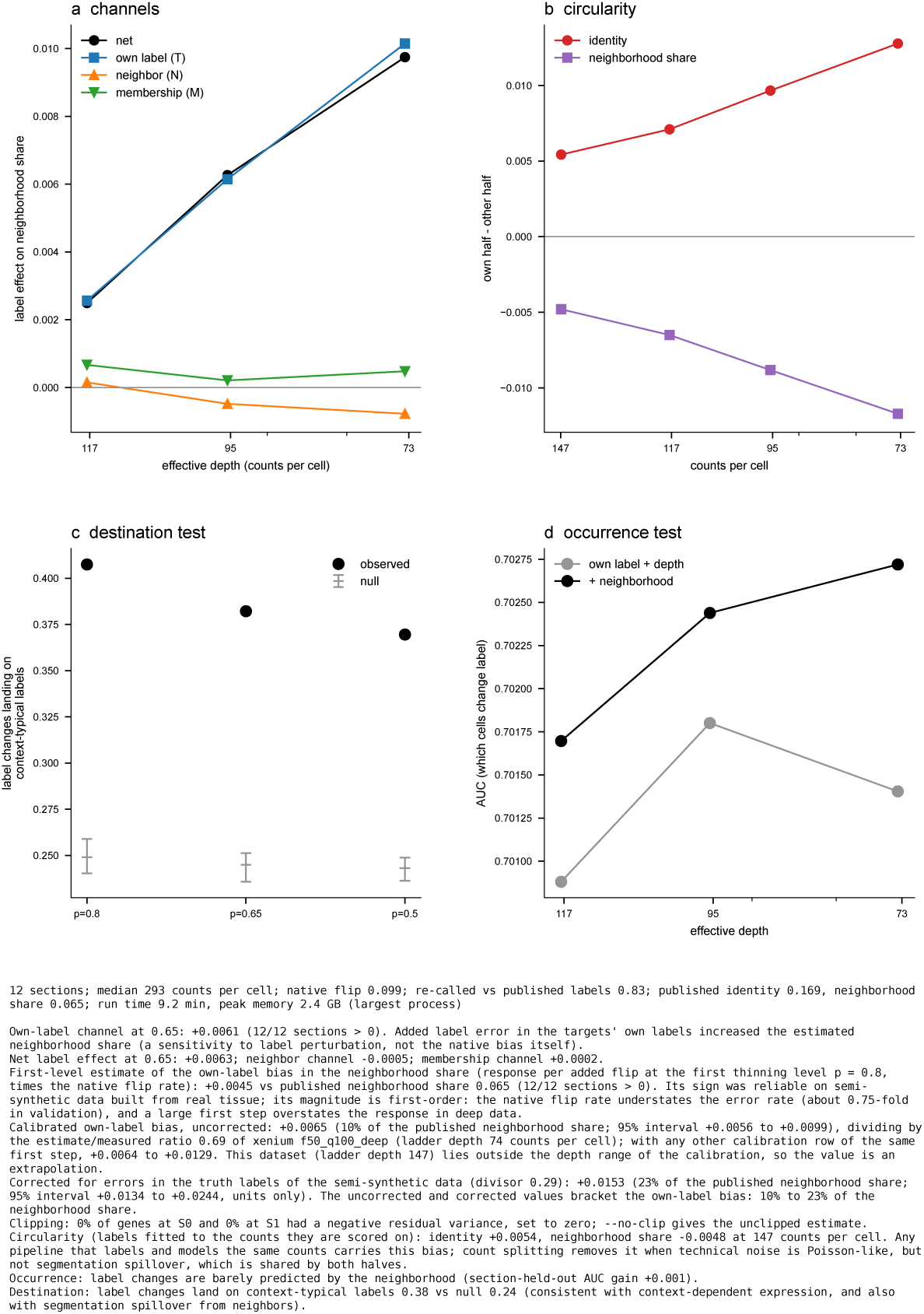
nichepeek. The one-page report for the mouse colon stroma dataset, produced by the single call shown with release 0.3.5, a first thinning step of *p* = 0.8, and the packaged calibration table. The report gives the uncorrected and the truth-corrected calibrated value. Its intervals resample the semi-synthetic units only, such that they differ from the joint intervals in the text. Because the thinning levels differ from the main runs, its other values (for example of the destination test) differ slightly from those in the text. **a**, label effect by channel against depth; **b**, circularity; **c**, destination test; **d**, occurrence test; below, a plain-language reading with the limits of the diagnostic. Top, the call that produced it (Python, or the equivalent command line); the input is an AnnData object or .h5ad file with published labels, section identifiers, coordinates, and raw counts.

**Figure S4:**
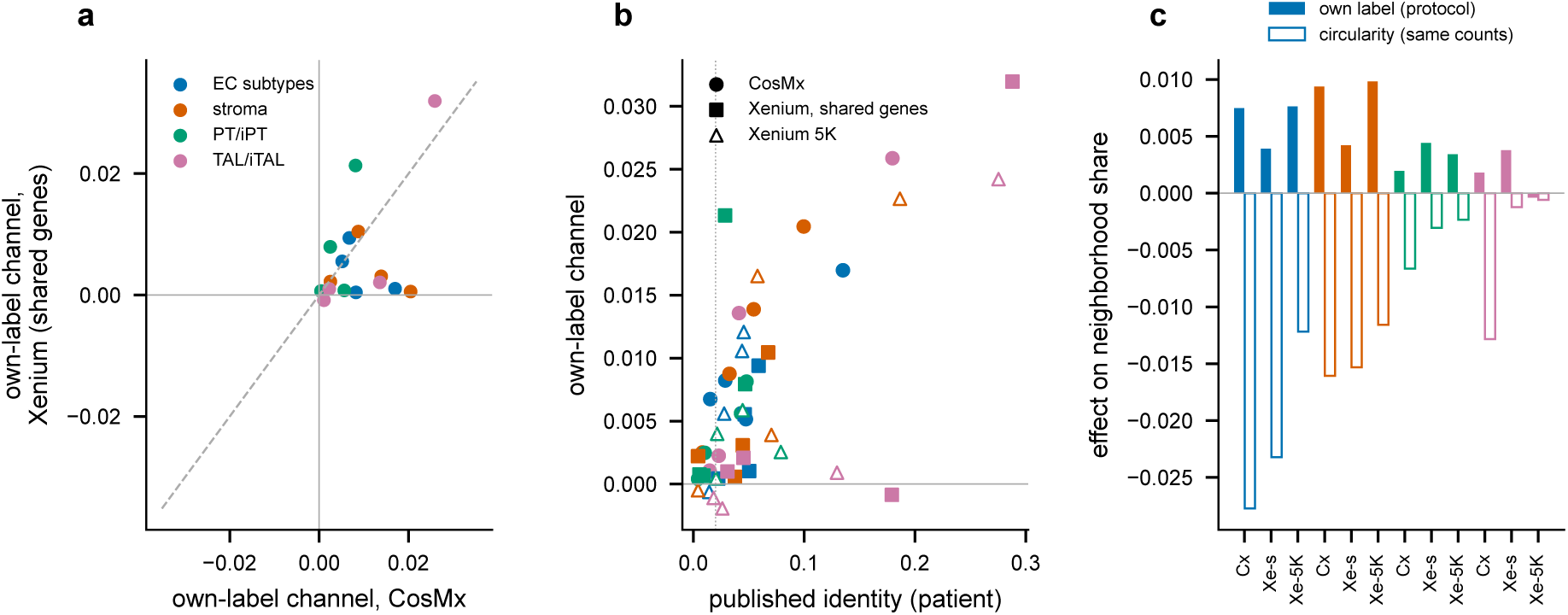
Kidney, samples from the same patients on two platforms. **a**, Own-label channel per patient and target set on CosMx against Xenium restricted to the shared genes (middle thinning level); dashed, identity. **b**, Own-label channel against published identity for every patient, target set, and platform (filled circles, CosMx; filled squares, Xenium shared genes; open triangles, full Xenium panel); dotted, identity 0.02. **c**, Per run, the own-label channel (filled) and the circularity on the neighborhood share at native depth (open), colored by target set as in a; Cx, CosMx; Xe-s, Xenium restricted to the shared genes; Xe-5K, the full Xenium panel. EC, endothelial cells; PT and iPT, proximal tubule and injured proximal tubule; TAL and iTAL, thick ascending limb and injured thick ascending limb.

**Figure S5:**
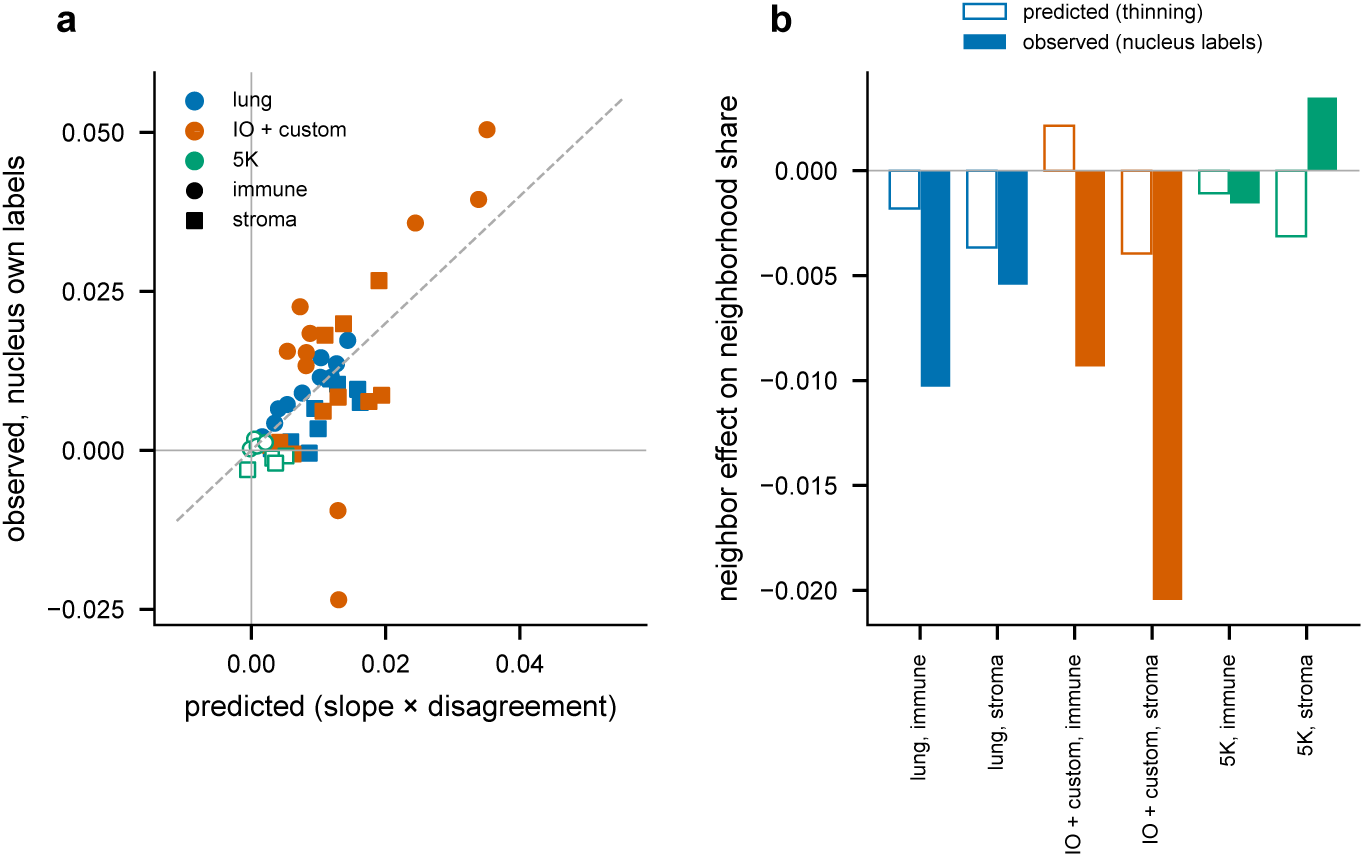
Lung cancer, whole-cell against nucleus labels of the same cells. **a**, Per crop, the effect on the neighborhood share of replacing the targets’ own labels by their nucleus labels (observed) against the protocol’s prediction (own-label response per added flip times the disagreement between re-called and nucleus labels); filled, runs with identity of at least 0.02; open, runs below it (both on the 5K panel); dashed, identity. Lung, lung panel; IO + custom, immuno-oncology panel with custom genes; 5K, 5K panel. **b**, Per run, the effect of nucleus labels in the neighbor role (filled) and the thinning prediction (open); medians over crops.

**Figure S6:**
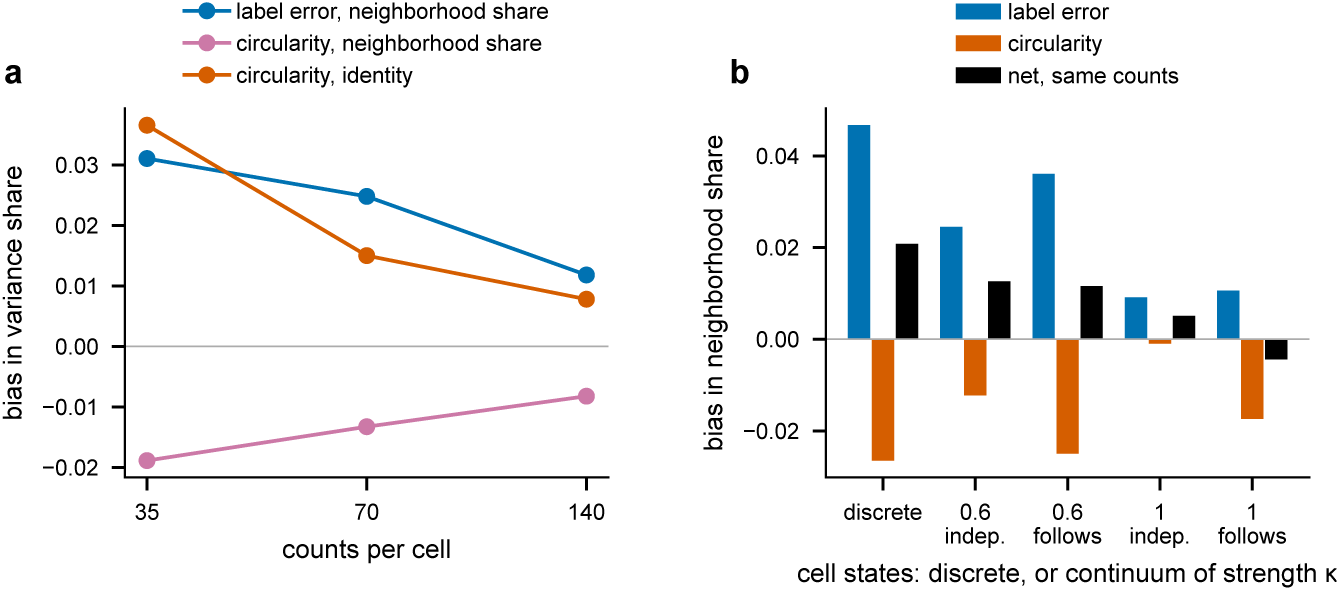
Simulations: depth and regimes. **a**, Bias from label error and from circularity against depth, in the neighborhood share and in identity. **b**, Bias in the neighborhood share from label error, from circularity, and net, in the regimes of Fig. 2b.

**Figure S7:**
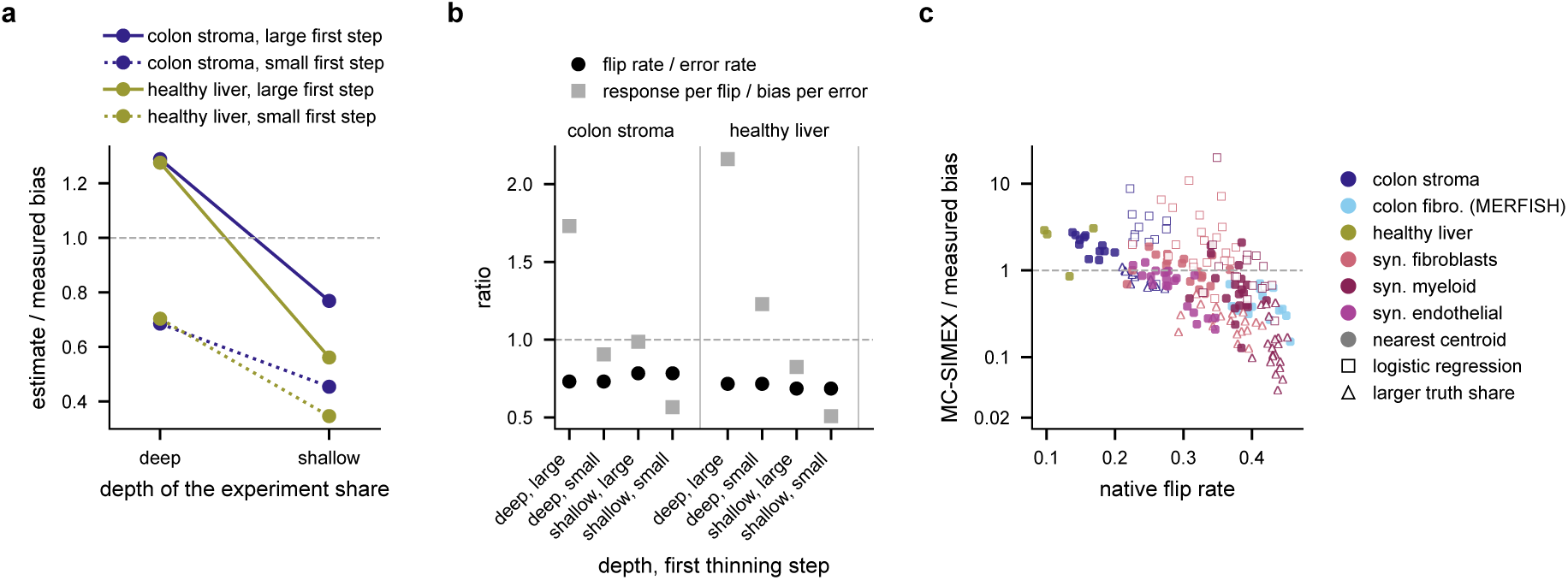
The size of the estimate. **a**, Estimate over measured own-label bias by depth of the experiment share, with a large (*p* = 0.5) and a small (*p* = 0.8) first thinning step; medians over independent units. **b**, The two factors of the estimate: native flip rate over the true error rate, and response per added flip over the measured bias per unit of error, per setting. **c**, Misclassification SIMEX (linear extrapolation) over the measured own-label bias against the native flip rate; symbols and axis limits as in Fig. 4c. Panels a and b are measured against the truth labels as called; Supplementary Note 3 corrects for their own errors.

**Figure S8:**
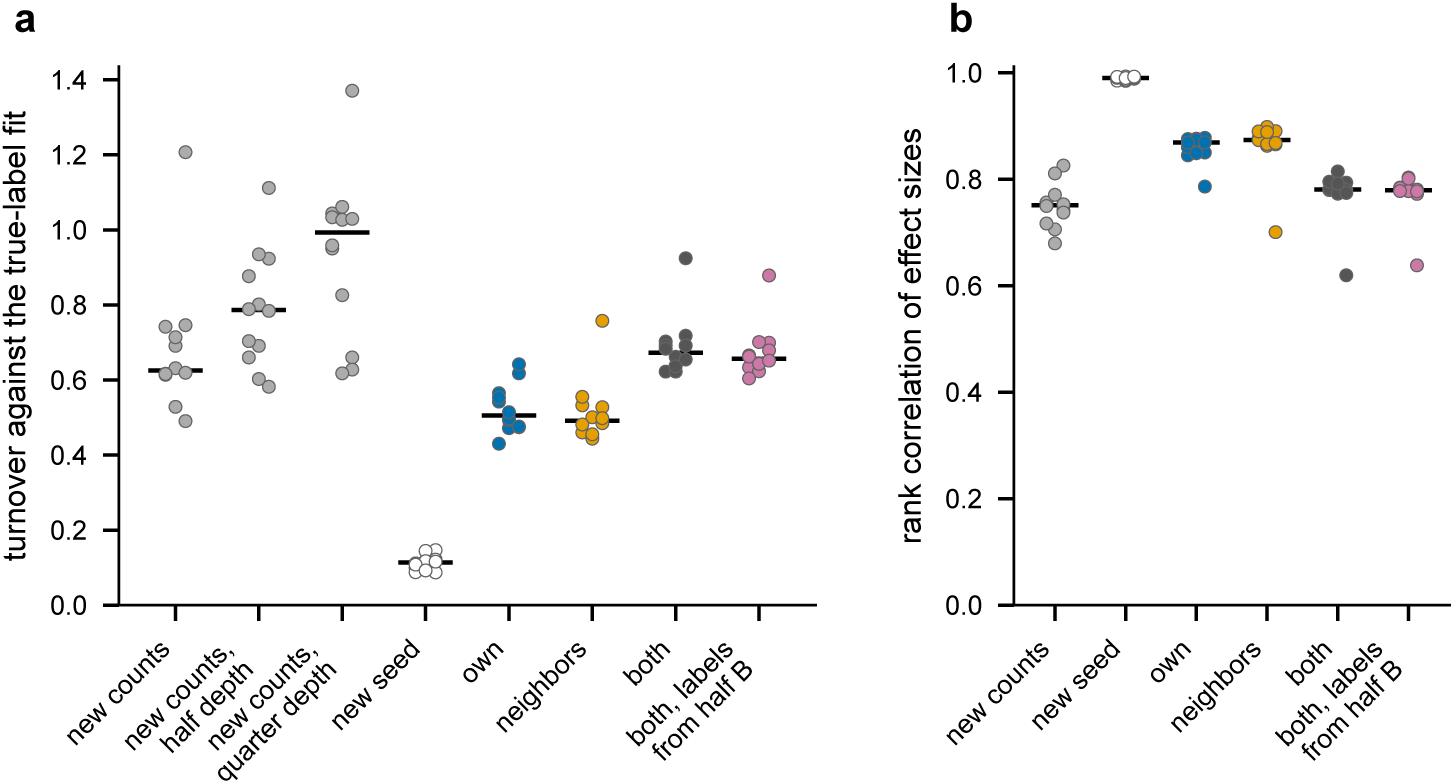
Turnover of Niche-DE calls on semi-synthetic colon data. **a**, Calls of the true-label fit that a second fit lost or made anew, as a share of its calls, per section: new counts (the other half) at full, half, and a quarter of the depth, a new seed of Niche-DE, and the re-called labels in the cell’s own role, in the neighbors’ role, or in both, the last also with labels re-called on half B. Bars, medians. **b**, Spearman correlation of the per-kernel effect sizes with those of the true-label fit. New counts alone changed about half of the calls or more, but almost never reversed one.

## Supplementary tables

**Table S1:** Label effect on the neighborhood share by channel. (middle thinning level; count splitting, fixed targets). Medians over sections, with 95% intervals from a cluster bootstrap over independent units (mice, donors, or patients), under them. *n*, sections or crops; depth, counts per cell that the model sees at this level (half the native depth, thinned); *I*, published identity; flip, native flip rate (share of target labels that change when half A is resampled); the membership column *M* re-selects the targets by the re-called labels and is not commensurable with *T* and *N* ; units, independent units whose mean own-label (*T*) or net effect is positive; *P*, two-sided sign test of the net effect over units. For liver (three donors), no interval and no test (Methods). LR, logistic-regression labeler.

| Population | *n* | depth | *I* | flip | own label *T* | neighbor *N* | membership *M* | net | units *T*>0 / net>0 | *P* (net) |
|---|---|---|---|---|---|---|---|---|---|---|
| Colon stroma (Xenium) | 12 | 37 | 0.169 | 0.099 | +0.032<br>[+0.030, +0.038] | −0.001<br>[−0.002, 0.000] | 0.000<br>[−0.001, +0.002] | +0.030<br>[+0.025, +0.038] | 12/12 of 12 | 0.00049 |
| Colon fibroblasts (MERFISH) | 16 | 13 | 0.083 | 0.354 | +0.019<br>[+0.013, +0.043] | −0.008<br>[−0.012, −0.002] | +0.006<br>[−0.005, +0.021] | +0.006<br>[+0.001, +0.026] | 6/5 of 6 | 0.22 |
| Liver, healthy (MERFISH) | 4 | 27 | 0.098 | 0.085 | +0.058 | −0.020 | −0.065 | +0.030 | 3/2 of 3 | – |
| Liver, fibrotic (MERFISH) | 6 | 13 | 0.008 | 0.347 | +0.001 | −0.007 | 0.000 | −0.007 | 3/0 of 3 | – |
| Synovium fibroblasts (Xenium) | 20 | 17 | 0.063 | 0.217 | +0.027<br>[+0.017, +0.031] | −0.005<br>[−0.006, −0.002] | −0.003<br>[−0.009, −0.001] | +0.022<br>[+0.013, +0.028] | 10/10 of 10 | 0.002 |
| Synovium myeloid (Xenium) | 20 | 12 | 0.040 | 0.280 | +0.035<br>[+0.031, +0.051] | −0.009<br>[−0.013, −0.008] | −0.042<br>[−0.048, −0.031] | +0.028<br>[+0.020, +0.044] | 10/10 of 10 | 0.002 |
| Synovium endothelial (Xenium) | 20 | 12 | 0.105 | 0.186 | +0.050<br>[+0.046, +0.054] | −0.010<br>[−0.011, −0.007] | +0.010<br>[+0.005, +0.027] | +0.035<br>[+0.028, +0.042] | 10/10 of 10 | 0.002 |
| Colon stroma, LR labeler | 12 | 37 | 0.169 | 0.171 | +0.048<br>[+0.045, +0.054] | −0.009<br>[−0.013, −0.007] | −0.004<br>[−0.005, −0.001] | +0.036<br>[+0.030, +0.041] | 12/12 of 12 | 0.00049 |
| Colon fibroblasts, LR labeler | 16 | 13 | 0.083 | 0.436 | +0.027<br>[+0.009, +0.043] | −0.010<br>[−0.022, −0.003] | +0.002<br>[−0.010, +0.005] | +0.006<br>[+0.006, +0.019] | 6/5 of 6 | 0.22 |

**Table S2:** MERFISH colon by mouse. (middle thinning level). Means over each mouse’s sections; NC, nearest-centroid labeler; last column, own-label channel with the logistic-regression labeler.

| Mouse | condition | *T* | *N* | *M* | net | *T* (LR) |
|---|---|---|---|---|---|---|
| 082421_D0_m6 | healthy | +0.017 | −0.004 | 0.000 | +0.009 | +0.011 |
| 082421_D0_m7 | healthy | +0.056 | −0.001 | −0.007 | +0.036 | +0.043 |
| 100221_D9_m2 | DSS day 9 | +0.028 | 0.000 | +0.007 | +0.021 | +0.023 |
| 100221_D9_m3 | DSS day 9 | +0.012 | −0.007 | +0.024 | +0.003 | +0.039 |
| 082421_D21_m1 | DSS day 21 | +0.021 | −0.006 | +0.008 | +0.012 | +0.017 |
| 082421_D21_m2 | DSS day 21 | +0.017 | −0.017 | −0.005 | −0.007 | +0.018 |

**Table S3:**
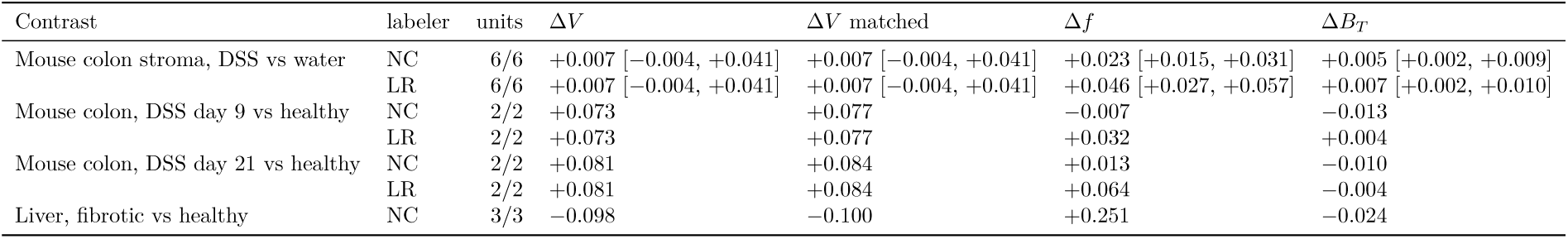
Condition contrasts. (Methods). Units, reference/alternative condition; Δ*V*, published contrast in the neighborhood share; matched, the same at matched depth; Δ*f*, difference in native flip rate; Δ*B_T_*, first-order label-error term. Intervals (95%, resampling units within conditions) only with at least five units per condition. NC, nearest-centroid; LR, logistic-regression labeler.

| Contrast | labeler | units | Δ*V* | Δ*V* matched | Δ*f* | Δ*B*<sub>*T*</sub> |
|---|---|---|---|---|---|---|
| Mouse colon stroma, DSS vs water | NC | 6/6 | +0.007 [−0.004, +0.041] | +0.007 [−0.004, +0.041] | +0.023 [+0.015, +0.031] | +0.005 [+0.002, +0.009] |
|  | LR | 6/6 | +0.007 [−0.004, +0.041] | +0.007 [−0.004, +0.041] | +0.046 [+0.027, +0.057] | +0.007 [+0.002, +0.010] |
| Mouse colon, DSS day 9 vs healthy | NC | 2/2 | +0.073 | +0.077 | −0.007 | −0.013 |
|  | LR | 2/2 | +0.073 | +0.077 | +0.032 | +0.004 |
| Mouse colon, DSS day 21 vs healthy | NC | 2/2 | +0.081 | +0.084 | +0.013 | −0.010 |
|  | LR | 2/2 | +0.081 | +0.084 | +0.064 | −0.004 |
| Liver, fibrotic vs healthy | NC | 3/3 | −0.098 | −0.100 | +0.251 | −0.024 |

**Table S4:** Robustness simulations. (planted neighborhood signal *α* = 0.5, six sections per row). True neighborhood share on half *B* with true labels; native own-label (*T*) and neighbor (*N*) channels with fixed targets; residual circularity after count splitting, measured against an independent replicate; share of a mislabeled cell’s true neighbors that carry its incorrect label.

| Regime | technical noise | true *V* | *T* | *N* | residual circularity | incorrect-label neighbors |
|---|---|---|---|---|---|---|
| discrete types | Poisson | 0.069 | +0.050 | +0.002 | −0.006 | 0.064 |
|  | overdispersed, CV<sup>2</sup> 0.1 | 0.036 | +0.030 | +0.003 | −0.004 | 0.057 |
|  | overdispersed, CV<sup>2</sup> 0.3 | 0.023 | +0.017 | +0.001 | −0.003 | 0.061 |
|  | spillover 10% | 0.076 | +0.040 | +0.003 | −0.002 | 0.068 |
|  | spillover 20% | 0.081 | +0.042 | +0.004 | +0.001 | 0.084 |
| continuum, following the neighborhood | Poisson | 0.081 | +0.025 | +0.004 | +0.001 | 0.052 |
|  | overdispersed, CV<sup>2</sup> 0.1 | 0.072 | +0.022 | +0.004 | −0.001 | 0.050 |
|  | overdispersed, CV<sup>2</sup> 0.3 | 0.036 | +0.011 | +0.004 | −0.001 | 0.049 |
|  | spillover 10% | 0.085 | +0.024 | +0.004 | +0.001 | 0.053 |
|  | spillover 20% | 0.091 | +0.023 | +0.005 | +0.001 | 0.073 |

**Table S5:** Sensitivity to analysis settings. Protocol runs with a different neighborhood size *k*, a different number of spatial groups, or without clipping negative per-gene residual variances at zero; middle thinning level (depth as in Supplementary Table S1); medians over sections. *I* and *V* with published labels.

| Population | setting | *I* | *V* | own label *T* | neighbor *N* | net |
|---|---|---|---|---|---|---|
| Mouse colon stroma (Xenium) | default (*k* = 20, 12 groups) | 0.169 | 0.065 | +0.032 | −0.0014 | +0.030 |
|  | *k* = 10 | 0.184 | 0.056 | +0.036 | −0.0022 | +0.032 |
|  | *k* = 40 | 0.139 | 0.072 | +0.025 | +0.0006 | +0.025 |
|  | 6 groups | 0.164 | 0.051 | +0.031 | +0.0005 | +0.032 |
|  | 24 groups | 0.175 | 0.072 | +0.033 | −0.0025 | +0.030 |
|  | no clipping | 0.169 | 0.065 | +0.032 | −0.0014 | +0.030 |
| Mouse colon fibroblasts (MERFISH) | default (*k* = 20, 12 groups) | 0.083 | 0.108 | +0.019 | −0.0077 | +0.006 |
|  | *k* = 10 | 0.088 | 0.095 | +0.019 | −0.0120 | −0.002 |
|  | *k* = 40 | 0.074 | 0.117 | +0.019 | −0.0015 | +0.019 |
|  | 6 groups | 0.065 | 0.074 | +0.013 | −0.0016 | +0.008 |
|  | 24 groups | 0.086 | 0.124 | +0.019 | −0.0094 | +0.004 |
|  | no clipping | 0.083 | 0.108 | +0.019 | −0.0078 | +0.007 |
| Human liver, healthy (MERFISH) | default (*k* = 20, 12 groups) | 0.098 | 0.132 | +0.058 | −0.0195 | +0.030 |
|  | *k* = 10 | 0.094 | 0.112 | +0.054 | −0.0270 | +0.018 |
|  | *k* = 40 | 0.099 | 0.142 | +0.056 | −0.0080 | +0.048 |
|  | 6 groups | 0.098 | 0.124 | +0.055 | −0.0147 | +0.033 |
|  | 24 groups | 0.100 | 0.139 | +0.059 | −0.0210 | +0.033 |
|  | no clipping | 0.098 | 0.132 | +0.058 | −0.0195 | +0.030 |
| Synovial myeloid | default (*k* = 20, 12 groups) | 0.040 | 0.147 | +0.035 | −0.0087 | +0.028 |
|  | no clipping | 0.040 | 0.147 | +0.035 | −0.0087 | +0.028 |

**Table S6:**
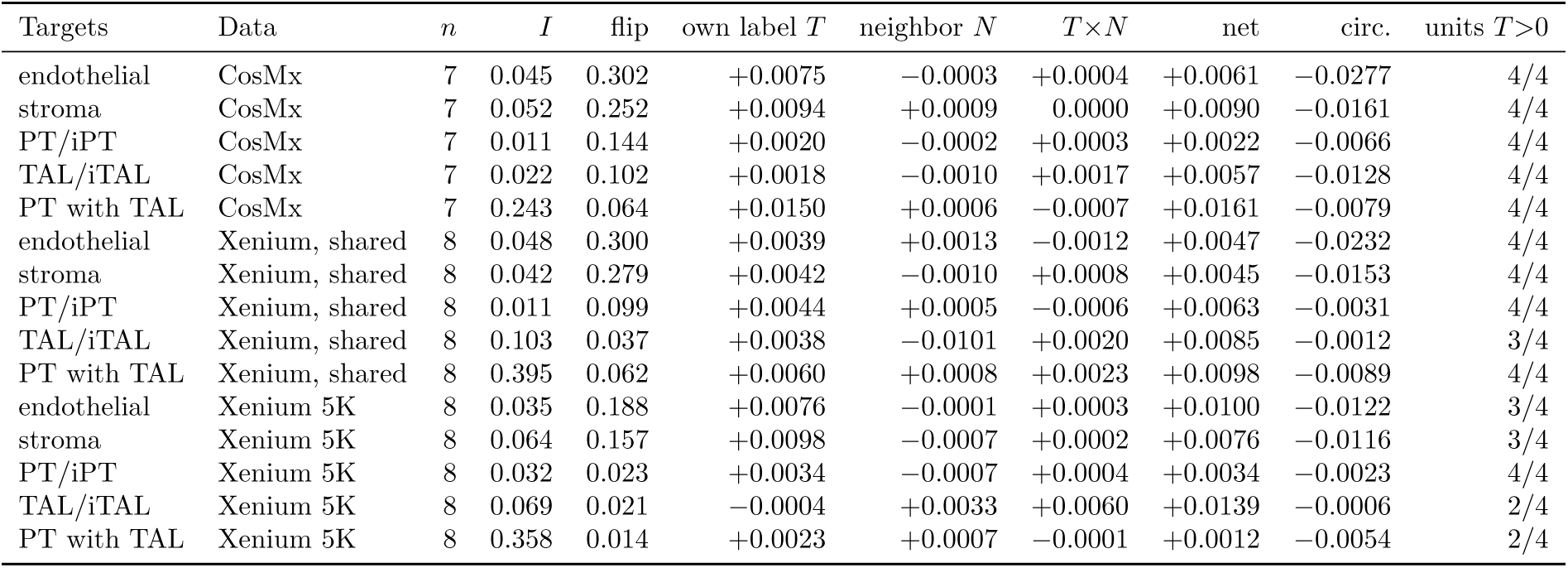
Kidney, samples from the same patients on two platforms. (middle thinning level *p* = 0.6; medians over sections or crops; count splitting, fixed targets). *I*, published identity; flip, native flip rate; circ., circularity on the neighborhood share at native depth (labels scored on the counts from which they were called); units, patients whose mean own-label effect is positive; *T×N*, interaction of the two channels. In every section the net effect equals *T* + *N* + *T×N* ; the medians, taken column by column, need not add up. PT with TAL, the proximal tubule and the thick ascending limb as one target set of two lineages.

**Table S7:** Lung cancer, whole-cell against nucleus labels. (middle thinning level *p* = 0.6; medians over crops; count splitting, fixed targets; published labels called from whole-cell counts). *I*, identity; flip, native flip rate; *T*, *N*, net, the protocol’s channels; dis., targets on which re-called and nucleus labels disagree; pred. and obs., predicted and observed effect on the neighborhood share of nucleus labels as own labels (*T*) or neighbor labels (*N*).

| Targets | Panel | *n* | *I* | flip | *T* | *N* | net | dis. | pred. *T* | obs. *T* | pred. *N* | obs. *N* |
|---|---|---|---|---|---|---|---|---|---|---|---|---|
| immune | lung | 10 | 0.052 | 0.126 | +0.0046 | −0.0015 | +0.0039 | 0.18 | +0.0089 | +0.0102 | −0.0018 | −0.0103 |
| stroma | lung | 10 | 0.053 | 0.106 | +0.0036 | −0.0015 | +0.0019 | 0.21 | +0.0097 | +0.0050 | −0.0037 | −0.0054 |
| immune | immuno-oncology | 10 | 0.059 | 0.147 | +0.0069 | +0.0018 | +0.0076 | 0.23 | +0.0108 | +0.0170 | +0.0021 | −0.0093 |
| stroma | immuno-oncology | 10 | 0.024 | 0.117 | +0.0035 | −0.0002 | +0.0034 | 0.28 | +0.0119 | +0.0080 | −0.0039 | −0.0205 |
| immune | Prime 5K | 4 | 0.003 | 0.184 | +0.0007 | −0.0006 | +0.0004 | 0.21 | +0.0007 | +0.0010 | −0.0011 | −0.0016 |
| stroma | Prime 5K | 4 | 0.014 | 0.148 | +0.0011 | −0.0001 | +0.0007 | 0.28 | +0.0034 | −0.0016 | −0.0031 | +0.0035 |

**Table S8:** Two labelers. Nearest-centroid (NC) and logistic-regression (LR) labelers on the same sections, middle thinning level; agreement, re-called against published labels among all cells of the section or crop (the three synovial populations share crops); units, units with positive mean own-label (*T*) or net effect.

| Population | labeler | agreement | flip | own label *T* | neighbor *N* | net | units *T*>0 / net>0 |
|---|---|---|---|---|---|---|---|
| Mouse colon stroma | NC | 0.83 | 0.099 | +0.032 | −0.001 | +0.030 | 12/12 of 12 |
|  | LR | 0.80 | 0.171 | +0.048 | −0.009 | +0.036 | 12/12 of 12 |
| Mouse colon fibroblasts | NC | 0.65 | 0.354 | +0.019 | −0.008 | +0.006 | 6/5 of 6 |
|  | LR | 0.56 | 0.436 | +0.027 | −0.010 | +0.006 | 6/5 of 6 |
| Human liver, healthy | NC | 0.86 | 0.085 | +0.058 | −0.020 | +0.030 | 3/2 of 3 |
|  | LR | 0.80 | 0.142 | +0.054 | −0.018 | +0.022 | 3/1 of 3 |
| Human liver, fibrotic | NC | 0.54 | 0.347 | +0.001 | −0.007 | −0.007 | 3/0 of 3 |
|  | LR | 0.51 | 0.371 | 0.000 | −0.002 | −0.003 | 3/0 of 3 |
| Synovial fibroblasts | NC | 0.62 | 0.217 | +0.027 | −0.005 | +0.022 | 10/10 of 10 |
|  | LR | 0.60 | 0.269 | +0.025 | −0.008 | +0.020 | 10/9 of 10 |
| Synovial myeloid | NC | 0.62 | 0.280 | +0.035 | −0.009 | +0.028 | 10/10 of 10 |
|  | LR | 0.60 | 0.338 | +0.028 | −0.014 | +0.014 | 10/8 of 10 |
| Synovial endothelial | NC | 0.62 | 0.186 | +0.050 | −0.010 | +0.035 | 10/10 of 10 |
|  | LR | 0.60 | 0.237 | +0.058 | −0.015 | +0.031 | 10/10 of 10 |

**Table S9:** Calibration of the first-level estimate on semi-synthetic data: estimate over directly measured own-label bias, by depth of the experiment and first thinning step. Deep: truth and experiment shares of equal depth (about 147 counts per cell in the experiment share in colon, such that the model sees about 74 after the split; the real colon data: 293 and 147); shallow: experiment at half that depth. Ratios are medians over independent units. Corrected: the measured bias extrapolated to a truth flip rate of zero (Supplementary Note 3). The liver values in parentheses did not pass the reliability rule and are shown for reference only. The table packaged with nichepeek carries both the uncorrected and the corrected factors.

| Tissue | first step | deep | shallow | shallow, deeper truth | deep, LR labeler |
|---|---|---|---|---|---|
| Colon stroma (Xenium) | *p* = 0.5 | 1.29 | 0.77 | 0.76 | 1.08 |
|  | *p* = 0.8 | 0.69 | 0.45 | – | – |
|  | *p* = 0.8, corrected | 0.29 | 0.29 | – | – |
| Healthy liver (MERFISH) | *p* = 0.5 | 1.28 | 0.56 | 0.43 | 0.98 |
|  | *p* = 0.8 | 0.70 | 0.35 | – | – |
|  | *p* = 0.8, corrected | (0.13) | (0.11) | – | – |

**Table S10:** Magnitude check and neighborhood groups. (Supplementary Note 3; semi-synthetic data built on the fly, *q* = 1, first step *p*_0_ = 0.8; re-calling labeler trained on the synthetic labels, except in the last block). NC, nearest-centroid; LR, logistic-regression labeler; *f*, truth share; *n*, sections or crops (units: mice, donors, or patients); flip, native flip rate; bias, measured own-label bias; est., first-level estimate; below, sections (units) with the estimate below the measured bias, a unit counted by the means over its sections; MC-SIMEX and oracle, misclassification SIMEX with the estimated and the true confusion matrix, linear extrapolation; quad., MC-SIMEX with quadratic extrapolation; ratios are ratios of medians over sections. Last three columns, median over sections of the Spearman correlation over groups between the identity loss and the score, the composition imbalance, or the group size. Last rows, further colon splits at the calibration setting: the file-based split of Fig. 4 and Supplementary Table S9, and seeds 1 to 9. Last block, the design comparison: the labeler that made the truth and the synthetic labels re-calls them, with the further colon splits of that design.

| Population | labeler, *f* | *n* (units) | flip | bias | est./bias | below | MC-SIMEX/bias | quad./bias | oracle/bias | ρ score | ρ comp. | ρ size |
|---|---|---|---|---|---|---|---|---|---|---|---|---|
| Mouse colon stroma (Xenium) | NC, *f* = 0.5 | 12 (12) | 0.159 | 0.0142 | 0.66 | 12 (12) | 2.12 | 3.40 | 2.21 | 0.85 | 0.84 | −0.21 |
|  | LR, *f* = 0.5 | 12 (12) | 0.242 | 0.0124 | 0.91 | 6 (6) | 2.57 | 4.71 | 2.82 | 0.69 | 0.64 | −0.22 |
|  | NC, *f* = 0.75 | 12 (12) | 0.229 | 0.0415 | 0.43 | 12 (12) | 0.84 | 1.74 | 0.81 | 0.91 | 0.89 | −0.12 |
| Mouse colon fibroblasts (MERFISH) | NC, *f* = 0.5 | 16 (6) | 0.414 | 0.0550 | 0.13 | 16 (6) | 0.43 | 1.12 | 0.38 | 0.66 | 0.58 | +0.23 |
| Human liver, healthy (MERFISH) | NC, *f* = 0.5 | 4 (3) | 0.117 | 0.0130 | 1.02 | 1 (1) | 2.64 | 6.25 | 2.71 | 0.46 | 0.06 | +0.40 |
| Synovial fibroblasts (Xenium) | NC, *f* = 0.5 | 20 (10) | 0.299 | 0.0227 | 0.61 | 18 (9) | 1.05 | 2.80 | 0.94 | 0.81 | 0.76 | +0.05 |
|  | LR, *f* = 0.5 | 20 (10) | 0.328 | 0.0082 | 1.49 | 5 (1) | 2.40 | 6.75 | 2.20 | 0.62 | 0.55 | −0.02 |
|  | NC, *f* = 0.75 | 20 (10) | 0.370 | 0.0611 | 0.29 | 20 (10) | 0.24 | 0.91 | 0.19 | 0.91 | 0.88 | −0.02 |
| Synovial myeloid (Xenium) | NC, *f* = 0.5 | 20 (10) | 0.382 | 0.0194 | 0.45 | 18 (9) | 0.58 | 2.26 | 0.46 | 0.68 | 0.66 | +0.04 |
|  | LR, *f* = 0.5 | 20 (10) | 0.397 | 0.0047 | 1.66 | 9 (5) | 1.66 | 7.72 | 1.01 | 0.54 | 0.44 | 0.00 |
|  | NC, *f* = 0.75 | 20 (10) | 0.434 | 0.0501 | 0.24 | 20 (10) | 0.11 | 0.57 | 0.08 | 0.82 | 0.74 | 0.00 |
| Synovial endothelial (Xenium) | NC, *f* = 0.5 | 20 (10) | 0.278 | 0.0353 | 0.35 | 19 (10) | 0.59 | 1.97 | 0.59 | 0.71 | 0.59 | +0.07 |
| Mouse colon stroma (Xenium) | NC, *f* = 0.5, files | 12 (12) | 0.158 | 0.0139 | 0.69 | 11 (11) | 2.15 | 3.60 | 2.25 | 0.81 | 0.85 | −0.18 |
|  | NC, *f* = 0.5, seed 1 | 12 (12) | 0.158 | 0.0133 | 0.77 | 11 (11) | 2.28 | 3.76 | 2.42 | 0.84 | 0.84 | −0.06 |
|  | NC, *f* = 0.5, seed 2 | 12 (12) | 0.159 | 0.0148 | 0.65 | 12 (12) | 2.05 | 3.41 | 2.16 | 0.77 | 0.86 | −0.07 |
|  | NC, *f* = 0.5, seed 3 | 12 (12) | 0.157 | 0.0134 | 0.78 | 12 (12) | 2.26 | 3.75 | 2.36 | 0.83 | 0.85 | −0.21 |
|  | NC, *f* = 0.5, seed 4 | 12 (12) | 0.158 | 0.0132 | 0.77 | 11 (11) | 2.29 | 3.71 | 2.34 | 0.84 | 0.84 | −0.01 |
|  | NC, *f* = 0.5, seed 5 | 12 (12) | 0.157 | 0.0136 | 0.73 | 11 (11) | 2.19 | 3.57 | 2.28 | 0.82 | 0.85 | −0.18 |
|  | NC, *f* = 0.5, seed 6 | 12 (12) | 0.157 | 0.0136 | 0.74 | 11 (11) | 2.21 | 3.70 | 2.28 | 0.82 | 0.85 | −0.21 |
|  | NC, *f* = 0.5, seed 7 | 12 (12) | 0.159 | 0.0137 | 0.76 | 11 (11) | 2.23 | 3.63 | 2.28 | 0.82 | 0.82 | −0.10 |
|  | NC, *f* = 0.5, seed 8 | 12 (12) | 0.159 | 0.0133 | 0.79 | 11 (11) | 2.28 | 3.82 | 2.42 | 0.83 | 0.83 | −0.15 |
|  | NC, *f* = 0.5, seed 9 | 12 (12) | 0.157 | 0.0148 | 0.68 | 11 (11) | 2.08 | 3.37 | 2.13 | 0.80 | 0.82 | −0.16 |
| Mouse colon stroma (Xenium) | NC, *f* = 0.5 | 12 (12) | 0.145 | 0.0105 | 0.90 | 9 (9) | 2.87 | 4.30 | 2.93 | 0.85 | 0.78 | −0.21 |
|  | LR, *f* = 0.5 | 12 (12) | 0.221 | 0.0172 | 0.75 | 12 (12) | 1.91 | 3.68 | 1.92 | 0.89 | 0.85 | 0.00 |
|  | NC, *f* = 0.75 | 12 (12) | 0.206 | 0.0326 | 0.51 | 12 (12) | 1.08 | 2.07 | 1.03 | 0.89 | 0.81 | −0.16 |
|  | NC, *f* = 0.5, seed 1 | 12 (12) | 0.145 | 0.0104 | 0.90 | 6 (6) | 2.84 | 4.44 | 2.89 | 0.83 | 0.75 | 0.00 |
|  | NC, *f* = 0.5, seed 2 | 12 (12) | 0.145 | 0.0107 | 0.95 | 9 (9) | 2.82 | 4.45 | 2.82 | 0.83 | 0.81 | −0.05 |
|  | NC, *f* = 0.5, seed 3 | 12 (12) | 0.144 | 0.0109 | 0.89 | 10 (10) | 2.74 | 4.21 | 2.74 | 0.82 | 0.78 | −0.23 |
|  | NC, *f* = 0.5, seed 4 | 12 (12) | 0.145 | 0.0101 | 0.99 | 7 (7) | 2.94 | 4.61 | 2.93 | 0.87 | 0.82 | −0.06 |
| Mouse colon fibroblasts (MERFISH) | NC, *f* = 0.5 | 16 (6) | 0.387 | 0.0501 | 0.14 | 16 (6) | 0.47 | 1.27 | 0.43 | 0.61 | 0.53 | +0.29 |
| Human liver, healthy (MERFISH) | NC, *f* = 0.5 | 4 (3) | 0.103 | 0.0146 | 0.97 | 2 (2) | 2.31 | 5.61 | 2.36 | 0.47 | 0.12 | +0.22 |
| Synovial fibroblasts (Xenium) | NC, *f* = 0.5 | 20 (10) | 0.284 | 0.0181 | 0.70 | 20 (10) | 1.35 | 3.34 | 1.32 | 0.84 | 0.80 | +0.06 |
|  | LR, *f* = 0.5 | 20 (10) | 0.330 | 0.0189 | 0.63 | 20 (10) | 0.90 | 2.52 | 0.78 | 0.90 | 0.85 | −0.08 |
|  | NC, *f* = 0.75 | 20 (10) | 0.348 | 0.0509 | 0.41 | 20 (10) | 0.39 | 1.25 | 0.34 | 0.91 | 0.90 | −0.01 |
| Synovial myeloid (Xenium) | NC, *f* = 0.5 | 20 (10) | 0.364 | 0.0187 | 0.47 | 20 (10) | 0.67 | 2.48 | 0.53 | 0.80 | 0.67 | +0.06 |
|  | LR, *f* = 0.5 | 20 (10) | 0.390 | 0.0106 | 0.60 | 19 (10) | 0.63 | 2.82 | 0.33 | 0.72 | 0.75 | +0.02 |
|  | NC, *f* = 0.75 | 20 (10) | 0.417 | 0.0517 | 0.21 | 20 (10) | 0.13 | 0.62 | 0.09 | 0.83 | 0.74 | 0.00 |
| Synovial endothelial (Xenium) | NC, *f* = 0.5 | 20 (10) | 0.257 | 0.0275 | 0.54 | 19 (10) | 0.87 | 2.40 | 0.86 | 0.60 | 0.52 | +0.04 |

**Table S11:** Published-style labels against exact truth. (simulated tissues; labels called by the nearest-centroid classifier on all counts and scored on the same counts; neighborhood share at native depth). Bias, published-style minus true labels, mean over sections; sections, sections with a positive bias. Identity strength sets how far apart the subtypes are: confusable at 0.5 and 1.0, well separated at 1.6.

| Counts per cell | Identity strength | Planted signal | True *V* | Bias | Sections >0 |
|---|---|---|---|---|---|
| 35 | 0.5 | none | 0.001 | +0.0090 | 4/6 |
| 35 | 0.5 | planted | 0.083 | +0.0178 | 5/6 |
| 35 | 1.0 | none | 0.000 | +0.0168 | 6/6 |
| 35 | 1.0 | planted | 0.070 | +0.0167 | 5/6 |
| 35 | 1.6 | none | 0.002 | +0.0056 | 4/6 |
| 35 | 1.6 | planted | 0.046 | −0.0008 | 4/6 |
| 70 | 0.5 | none | −0.001 | +0.0099 | 6/6 |
| 70 | 0.5 | planted | 0.078 | +0.0128 | 6/6 |
| 70 | 1.0 | none | 0.003 | +0.0041 | 5/6 |
| 70 | 1.0 | planted | 0.070 | +0.0088 | 6/6 |
| 70 | 1.6 | none | 0.001 | +0.0002 | 4/6 |
| 70 | 1.6 | planted | 0.040 | +0.0044 | 6/6 |
| 140 | 0.5 | none | 0.000 | +0.0062 | 5/6 |
| 140 | 0.5 | planted | 0.073 | +0.0101 | 6/6 |
| 140 | 1.0 | none | 0.000 | +0.0016 | 5/6 |
| 140 | 1.0 | planted | 0.067 | +0.0042 | 5/6 |
| 140 | 1.6 | none | −0.001 | +0.0023 | 5/6 |
| 140 | 1.6 | planted | 0.050 | −0.0014 | 2/6 |

**Table S12:** Published labels against semi-synthetic truth. (effect on the neighborhood share of the published labels against the truth labels, both on all counts of the experiment share, target population fixed; medians over sections). Rows with truth share 0.75 leave the experiment a quarter of the counts, such that they change the depth of the published labels and the quality of the truth together; they are shown but not used in the text.

| Population | Semi-synthetic data | Published labels, net | Sections >0 | Own-label channel |
|---|---|---|---|---|
| Colon stroma (Xenium) | built from the main run | −0.0053 | 0/12 | −0.0053 |
|  | file, full experiment share | −0.0049 | 0/12 | −0.0057 |
|  | file, experiment share halved | −0.0033 | 4/12 | −0.0029 |
|  | file, truth share 0.75 | +0.0021 | 7/12 | −0.0002 |
| Colon fibroblasts (MERFISH) | built from the main run | −0.0080 | 5/16 | −0.0111 |
| Liver, healthy (MERFISH) | built from the main run | −0.0084 | 0/4 | −0.0076 |
|  | file, full experiment share | −0.0042 | 0/4 | −0.0064 |
|  | file, experiment share halved | −0.0032 | 2/4 | −0.0035 |
|  | file, truth share 0.75 | −0.0019 | 1/4 | +0.0031 |
| Liver, fibrotic (MERFISH) | built from the main run | −0.0077 | 0/6 | −0.0070 |
| Synovial fibroblasts (Xenium) | built from the main run | −0.0194 | 0/20 | −0.0179 |
| Synovial myeloid (Xenium) | built from the main run | −0.0200 | 2/20 | −0.0188 |
| Synovial endothelial (Xenium) | built from the main run | −0.0148 | 1/20 | −0.0171 |

**Table S13:** Cluster-based labels against exact truth. (simulated tissues with a planted neighborhood signal; 30 principal components, a 15-nearest-neighbor graph, and Leiden clustering, each cluster annotated by its nearest reference centroid; labels called and scored on the same counts). Bias, published-style minus true labels as a fraction of the true neighborhood share, overall and by counts per cell; target error, share of target cells with an incorrect label.

| Labels | Clusters | Target error | Bias | 35 counts | 70 counts | 140 counts | Sections >0 |
|---|---|---|---|---|---|---|---|
| per cell, nearest centroid | – | 0.25 | 13% | 19% | 15% | 7% | 16/18 |
| Leiden, resolution 0.5 | 7.6 | 0.57 | 44% | 78% | 37% | 19% | 15/18 |
| Leiden, resolution 1.5 | 8.1 | 0.52 | 16% | 34% | 18% | −3% | 16/18 |
| Leiden, resolution 3 | 17.1 | 0.44 | 13% | 23% | 18% | −1% | 13/18 |

**Table S14:** Labelers with natural class priors in colon stroma. (Supplementary Note 2). Flat, the labelers of the main runs; prior, the nearest-centroid labeler with the published label frequencies as prior; uniform sample, logistic regression trained on a uniform sample of cells. Agreement, share of targets whose re-called label equals the published one; flip rate, native; own label *T*, at the middle level of these runs, which used the small first thinning step of the calibration and, therefore, differ from Supplementary Table S1; destination, share of label changes that land on the label typical of the neighborhood, observed and expected; estimate, first-level estimate of the own-label bias; DSS and water, its medians over the six mice of each group, with the Mann–Whitney *P* of their comparison.

| Labeler | Agreement | Flip rate | Own label *T* | Destination | Estimate | DSS | Water | *P* |
|---|---|---|---|---|---|---|---|---|
| Nearest centroid, flat | 0.83 | 0.099 | +0.0061 | – | 0.0045 | 0.0060 | 0.0040 | 0.002 |
| Nearest centroid, prior | 0.86 | 0.072 | +0.0062 | 0.33 / 0.21 | 0.0050 | 0.0058 | 0.0038 | 0.009 |
| Logistic regression, flat | 0.80 | 0.171 | +0.0100 | – | – | – | – | – |
| Logistic regression, uniform sample | 0.86 | 0.112 | +0.0086 | 0.34 / 0.23 | – | 0.0090 | 0.0055 | 0.015 |

**Table S15:** Label sets of the semi-synthetic colon data. (Supplementary Note 3). Truth labels are called on the truth share; synthetic published labels on the experiment share, at equal depth (deep) or thinned to half (shallow). Counts, median counts per cell from which the labels are called; flip, share of targets whose label changes on resampling; agreement, share of targets with the same label as the real published labels (truth labels) or as the truth labels (synthetic published labels). Agreement of the truth labels (truth share one half) with the published labels ranged from 0.75 to 0.81 over DSS sections and from 0.83 to 0.85 over water-treated sections.

| Label set | Counts | Flip | Agreement |
|---|---|---|---|
| Truth, truth share one half | 146 | 0.10 | 0.82 |
| Truth, truth share three quarters | 220 | 0.08 | 0.83 |
| Synthetic published, deep | 147 | – | 0.86 |
| Synthetic published, shallow | 74 | – | 0.82 |

**Table S16:** Calibrated own-label bias. (Supplementary Note 3). Estimate, first-level estimate with a small first thinning step; calibrated, the estimate divided by the ratio measured on semi-synthetic data of the same tissue; share, calibrated bias as a share of the published neighborhood share; interval, bootstrap over independent units, or the three donor values for liver. Corrected, with the measured bias extrapolated to a truth flip rate of zero by a linear or a quadratic fit. The corrected liver values did not pass the reliability rule and are a sensitivity analysis only. For MERFISH colon, the estimate is divided by the shallow ratios of both tissues and given as a range.

| Population | Truth labels | Estimate | Calibrated | Share | Interval |
|---|---|---|---|---|---|
| Colon stroma (Xenium) | as called | 0.0045 | 0.0065 | 10% | 8% to 16% |
|  | corrected, linear |  | 0.0153 | 23% | 20% to 40% |
|  | corrected, quadratic |  |  | 19% |  |
| Healthy liver (MERFISH) | as called | 0.0083 | 0.0118 | 9% | 11%, 8%, 6% |
|  | corrected, linear |  |  |  | (32% to 62%) |
| Colon fibroblasts (MERFISH) | as called |  | 0.015 to 0.019 | 13% to 18% | 7% to 94% |

**Table S17:** Predictions of the Niche-DE runs on semi-synthetic colon data. (Supplementary Note 4). Each prediction and its decision rule were written in a dated plan before the run. N, first run (counts, depth, and seed; sections other than the pilot section); C, confirmation on a third split; R, F, D, published labels, with the study’s labels (R2, F2) reported without a rule because they could be tested in three sections only. Turnover, calls made or lost against the true-label fit, as a share of its calls; label error, re-called labels in both roles; excess, label-error turnover minus counting-noise turnover.

|  | Prediction | Result | Held |
|---|---|---|---|
| N1 | Counting noise changes at least 0.40 of the calls | 11 of 11 sections; median 0.63 | yes |
| N2 | Label error changes no more calls than counting noise (excess at most 0.15) | 10 of 11; median excess −0.01 | yes |
| N3 | Under label error, effect sizes rank alike (Spearman at least 0.70) and strong calls stay (at least 0.75) | 10 of 11 | yes |
| N4 | A new split changes the excess by at most 0.10 | 7 of 11; median change 0.08 | no |
| N5 | Counting-noise turnover rises from 100% to 50% and 25% of the depth | 9 of 11; medians 0.63, 0.79, 0.96 | yes |
| N6 | A second seed of Niche-DE changes at most 0.05 of the calls | 0 of 11; at most 0.15 | no |
| C1 | Most strict reversals of the inflamed section recur | 47 of 55 | yes |
| C2 | Counting noise and the seed reverse at most two true-label calls per section | at most 2; 8 in all | yes |
| C3 | Most reversals of fibroblast genes in the inflamed section recur | 24 of 27 | yes |
| C4 | Most false claims of label error in the inflamed section recur | 1173 of 1513 | yes |
| C5 | False claims of label error recur more often than those of counting noise | 0.60 against 0.33 | yes |
| R1 | Synthetic published labels reverse claims, at least ten in at least three sections | 0 | no |
| F1 | Synthetic published labels make more than twice the false claims of counting noise in most sections | 3 of 12 sections | no |
| D1 | Their false claims rise with their disagreement from the truth (Spearman at least one half) | 0.16 | no |
| R2 | The study’s labels reverse claims | 26, in 3 of 3 sections | reported |
| F2 | The study’s labels make more than twice the false claims of counting noise | 2 of 3 sections | reported |

**Table S18:**
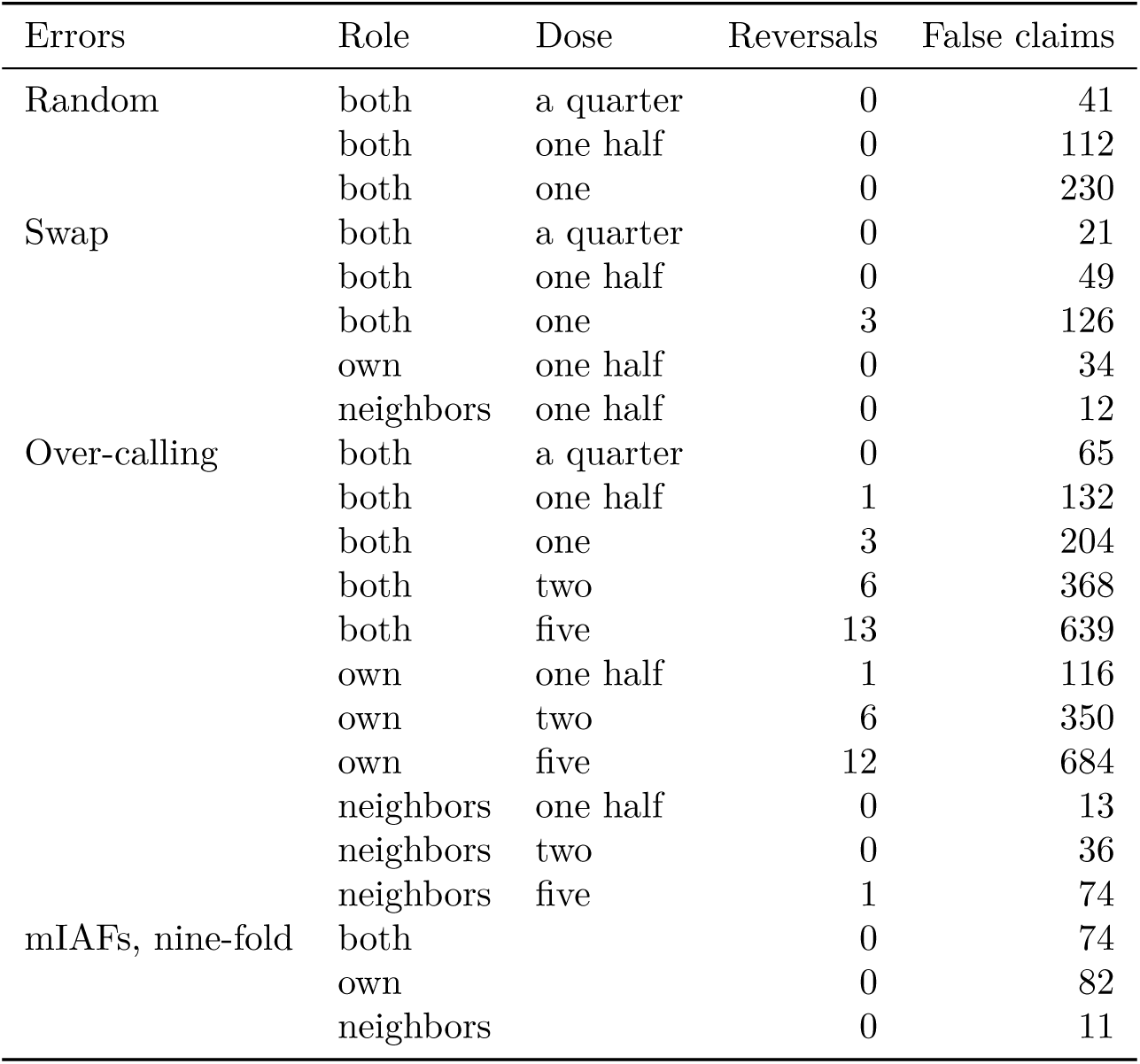
Injected errors. (Supplementary Note 4). Robust claims reversed and persistent false claims, summed over the four sections of the titration. Dose, number of changed labels as a multiple of the size of the smaller type of the most confused pair; random, errors among fibroblast subtypes that keep the composition; swap, within the pair; over-calling, cells of other subtypes given the label of the smaller type, placed at random among them. The nine-fold inflation of mIAFs ran in the two control sections.

**Table S19:**
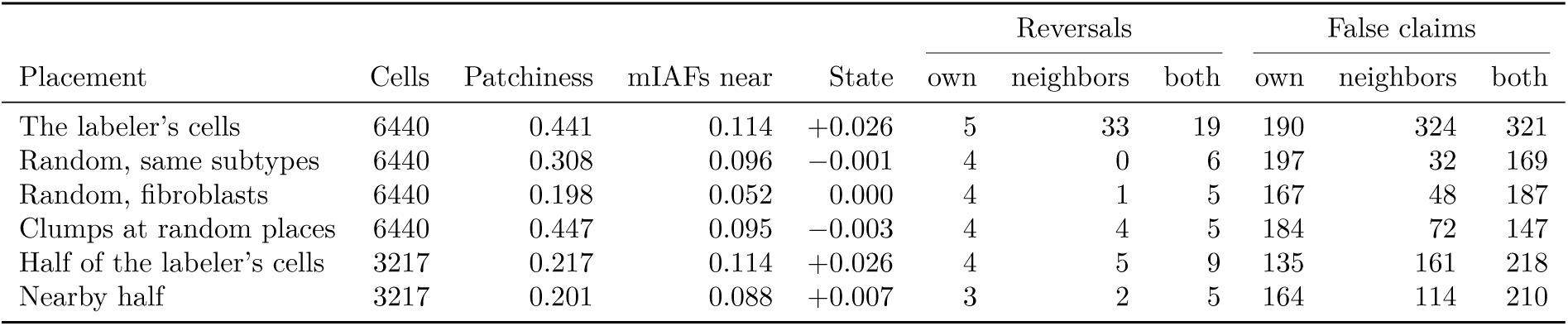
Placement of the labeler’s over-call in the inflamed section. (Supplementary Note 4). Cells, number of changed labels (mean of two splits); patchiness, share of mislabeled cells among the 20 nearest neighbors of a mislabeled cell; mIAFs near, share of true mIAFs among those neighbors; state, mean score of the changed cells for mIAFs on half B above the mean of cells of the same true subtypes. Reversals and false claims by the role of the changed labels: own, neighbors, both. Half and nearby half have half the labeler’s errors and their own base rate of mislabeled cells (0.097, against 0.193).

**Table S20:**
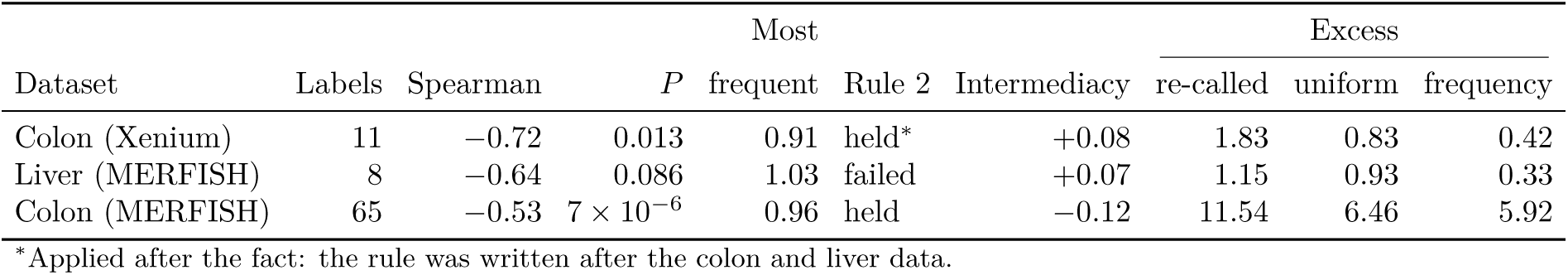
Labelers trained on re-called labels, in three datasets. (Supplementary Note 4). Chain, cells called on half A by a labeler trained on the synthetic published labels over those called by one trained on the study’s labels, per label, both on the experiment share; labels with at least 200 truth cells. Spearman, correlation of the chain with log truth share. Rule 2, written before the further MERFISH data were built (Supplementary Note 4). Intermediacy, Spearman correlation of the chain with how far a label lay between others in expression. Excess, sum over labels of the chain above one, for the re-called labels and for the two null labelers, trained on the study’s labels with random errors at the rate of the synthetic published labels, drawn uniformly or by frequency. No null correlated significantly negatively with log truth share.

**Table S21:**
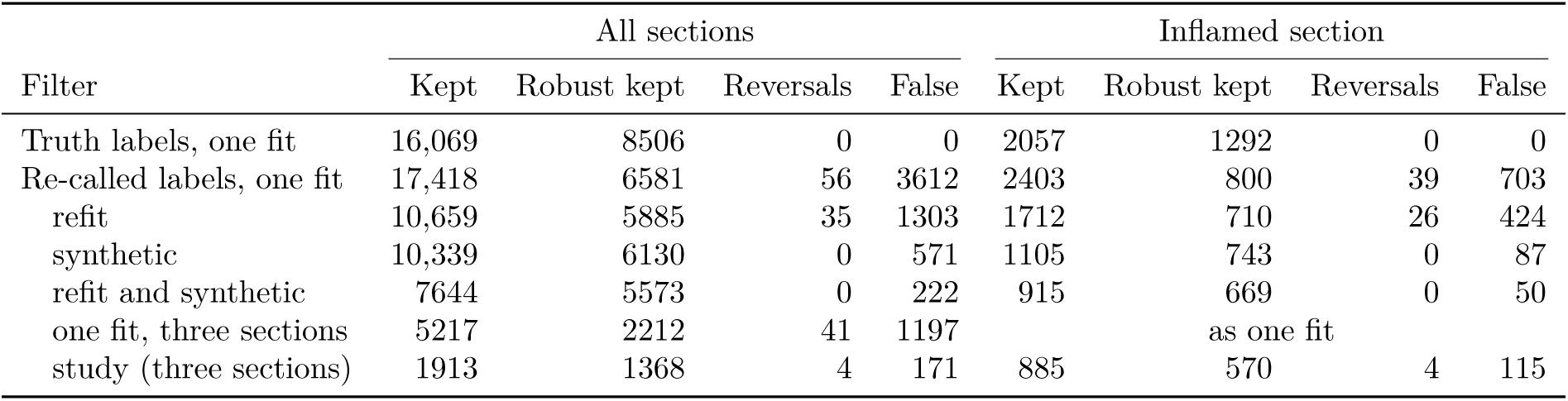
Agreement filters on the calls of Niche-DE. (Supplementary Note 4). Claims kept by each filter, robust claims kept out of all robust claims (8506 over all sections, 1292 in the inflamed section), reversals of robust claims in the fit on split 0 (not the strict reversals of Section 2.6, which recur on a second split), and false claims, scored against the true-label fits. Re-called labels: one fit on split 0; refit, the same claim on splits 0 and 1; synthetic or study, the same claim in a fit with the synthetic published labels or with the study’s labels; the study’s labels could be tested in three sections, and its row counts those sections only, as does the row for one fit in the same sections (3140 robust claims). Truth labels: the reference fit on split 0.

| Filter | All sections |  |  |  | Inflamed section |  |  |  |
|---|---|---|---|---|---|---|---|---|
|  | Kept | Robust kept | Reversals | False | Kept | Robust kept | Reversals | False |
| Truth labels, one fit | 16,069 | 8506 | 0 | 0 | 2057 | 1292 | 0 | 0 |
| Re-called labels, one fit | 17,418 | 6581 | 56 | 3612 | 2403 | 800 | 39 | 703 |
| refit | 10,659 | 5885 | 35 | 1303 | 1712 | 710 | 26 | 424 |
| synthetic | 10,339 | 6130 | 0 | 571 | 1105 | 743 | 0 | 87 |
| refit and synthetic | 7644 | 5573 | 0 | 222 | 915 | 669 | 0 | 50 |
| one fit, three sections | 5217 | 2212 | 41 | 1197 | as one fit |  |  |  |
| study (three sections) | 1913 | 1368 | 4 | 171 | 885 | 570 | 4 | 115 |

**Table S22:** Niche-DE with labels from a labeler trained on the study’s labels. (Supplementary Note 4). Study, labels re-called on half A by the labeler with centroids from the study’s labels; re-called, the labels of the main analysis, from the labeler with centroids from the synthetic published labels. Agreement with the truth on all analyzed cells and cells called mIAFs, split 0. Strict reversals of robust claims by the role of the labels (own, neighbors, both) and persistent false claims with the labels in both roles.

|  | Agreement |  | mIAFs cells |  |  | Reversals, study |  |  | Reversals, re-called |  |  | False claims |  |
|---|---|---|---|---|---|---|---|---|---|---|---|---|---|
| Section | study | re-called | truth | study | re-called | own | neighbors | both | own | neighbors | both | study | re-called |
| WT2-DSS | 0.80 | 0.68 | 1992 | 2272 | 7722 | 0 | 1 | 0 | 9 | 36 | 26 | 196 | 424 |
| KO2-Water | 0.85 | 0.83 | 139 | 215 | 325 | 0 | 0 | 0 | 0 | 5 | 5 | 43 | 91 |
| WT1-Water | 0.84 | 0.82 | 76 | 134 | 200 | 0 | 0 | 0 | 0 | 0 | 0 | 16 | 8 |
| KO3-DSS | 0.80 | 0.77 | 1142 | 1411 | 2036 | 0 | 0 | 0 | 0 | 2 | 0 | 66 | 137 |
| KO2-DSS | 0.80 | 0.76 | 1208 | 1476 | 2257 | 0 | 0 | 0 | 0 | 0 | 0 | 32 | 50 |

## References

[1] Damien Arnol, Denis Schapiro, Bernd Bodenmiller, Julio Saez-Rodriguez, and Oliver Stegle. Modeling cell-cell interactions from spatial molecular data with spatial variance component analysis. Cell Reports, 29(1):202–211.e6, 2019. doi: 10.1016/j.celrep.2019.08.077.

[2] Jovan Tanevski, Ricardo Omar Ramirez Flores, Attila Gabor, Denis Schapiro, and Julio Saez-Rodriguez. Explainable multiview framework for dissecting spatial relationships from highly multiplexed data. Genome Biology, 23:97, 2022. doi: 10.1186/s13059-022-02663-5.

[3] David S. Fischer, Anna C. Schaar, and Fabian J. Theis. Modeling intercellular communication in tissues using spatial graphs of cells. Nature Biotechnology, 41:332–336, 2023. doi: 10.1038/s41587-022-01467-z.

[4] Kaishu Mason, Anuja Sathe, Paul R. Hess, Jiazhen Rong, Chi-Yun Wu, Emma Furth, Katalin Susztak, Jonathan Levinsohn, Hanlee P. Ji, and Nancy Zhang. Niche-DE: niche-differential gene expression analysis in spatial transcriptomics data identifies context-dependent cell-cell interactions. Genome Biology, 25:14, 2024. doi: 10.1186/s13059-023-03159-6.

[5] Zoe Fewell, George Davey Smith, and Jonathan A. C. Sterne. The impact of residual and unmeasured confounding in epidemiologic studies: a simulation study. American Journal of Epidemiology, 166(6):646–655, 2007. doi: 10.1093/aje/kwm165.

[6] Jacob Westfall and Tal Yarkoni. Statistically controlling for confounding constructs is harder than you think. PLoS ONE, 11(3):e0152719, 2016. doi: 10.1371/journal.pone.0152719.

[7] Viktor Petukhov, Rosalind J. Xu, Ruslan A. Soldatov, Paolo Cadinu, Konstantin Khodosevich, Jeffrey R. Moffitt, and Peter V. Kharchenko. Cell segmentation in imaging-based spatial transcriptomics. Nature Biotechnology, 40:345–354, 2022. doi: 10.1038/s41587-021-01044-w.

[8] Jonathan Mitchel, Teng Gao, Viktor Petukhov, Eli Cole, and Peter V. Kharchenko. Impact and correction of segmentation errors in spatial transcriptomics. Nature Genetics, 58(2): 434–444, 2026. doi: 10.1038/s41588-025-02497-4.

[9] Can Ergen and Nir Yosef. ResolVI: addressing noise and bias in spatial transcriptomics. Nature Methods, 23:2003–2014, 2026. doi: 10.1038/s41592-026-03212-9.

[10] Mariia Bilous, Daria Buszta, Jonathan Bac, Senbai Kang, Yixing Dong, Stephanie Tissot, Sylvie Andre, Marina Alexandre Gaveta, Christel Voize, Solange Peters, Krisztian Homicsko, and Raphael Gottardo. Resolving sensitivity, specificity and signal contamination in Xenium spatial transcriptomics. Nature Methods, 23:1152–1162, 2026. doi: 10.1038/s41592-026-03089-8.

[11] Ken To, J. Patrick Pett, Olli Dufva, et al. Spatially resolved multiomics of human meningeal development reveal lineage and disease dynamics. Nature Cell Biology, 28:2169–2178, 2026. doi: 10.1038/s41556-026-02075-8.

[12] Chenxing Zhang, Lin Gao, Yuxuan Hu, and Zhengyang Huang. RobustCCC: a robustness evaluation tool for cell-cell communication methods. Frontiers in Genetics, 14:1236956, 2023. doi: 10.3389/fgene.2023.1236956.

[13] Jingyang Qian, Xin Shao, Hudong Bao, Yin Fang, Wenbo Guo, Chengyu Li, Anyao Li, Hua Hua, and Xiaohui Fan. Identification and characterization of cell niches in tissue from spatial omics data at single-cell resolution. Nature Communications, 16:1693, 2025. doi: 10.1038/s41467-025-57029-9.

[14] Runzhi Xie, Zekun Wang, Jianrui Liu, Han Xu, Yafei Xu, Jiadong Lin, Yuxuan Hu, and Lin Gao. HRCHY-CytoCommunity identifies hierarchical tissue organization in cell-type spatial maps. Nature Communications, 17:3312, 2026. doi: 10.1038/s41467-026-70069-z.

[15] Lucy L. Gao, Jacob Bien, and Daniela Witten. Selective inference for hierarchical clustering. Journal of the American Statistical Association, 119(545):332–342, 2024. doi: 10.1080/01621459.2022.2116331.

[16] Anna Neufeld, Lucy L. Gao, Joshua Popp, Alexis Battle, and Daniela Witten. Inference after latent variable estimation for single-cell RNA sequencing data. Biostatistics, 25(1): 270–287, 2024. doi: 10.1093/biostatistics/kxac047.

[17] Anna Neufeld, Ameer Dharamshi, Lucy L. Gao, and Daniela Witten. Data thinning for convolution-closed distributions. Journal of Machine Learning Research, 25(57):1–35, 2024. URL https://jmlr.org/papers/v25/23-0446.html.

[18] Vladislav Pokatayev, Alok Jaiswal, Angela R. Shih, Åsa Segerstolpe, Bihua Li, Elizabeth A. Creasey, Yanhua Zhao, Crystal Lin, Shane Murphy, Chih-Hung Chou, Daniel B. Graham, and Ramnik J. Xavier. Bidirectional CRISPR screens decode a GLIS3-dependent fibrotic cell circuit. Nature, 650:997–1006, 2026. doi: 10.1038/s41586-025-09907-x.

[19] Paolo Cadinu, Kisha N. Sivanathan, Aditya Misra, Rosalind J. Xu, Davide Mangani, Evan Yang, Joseph M. Rone, Katherine Tooley, Yoon-Chul Kye, Lloyd Bod, Ludwig Geistlinger, Tyrone Lee, Randall T. Mertens, Noriaki Ono, Gang Wang, Liliana Sanmarco, Francisco J. Quintana, Ana C. Anderson, Vijay K. Kuchroo, Jeffrey R. Moffitt, and Roni Nowarski. Charting the cellular biogeography in colitis reveals fibroblast trajectories and coordinated spatial remodeling. Cell, 187(8):2010–2028.e30, 2024. doi: 10.1016/j.cell.2024.03.013.

[20] Benjamin R. Watson, Bidisha Paul, Raza-Ur Rahman, Liat Amir-Zilberstein, Åsa Segerstolpe, Ephraim T. Epstein, Shaun Murphy, Ludwig Geistlinger, Taemoo Lee, Annie Shih, Jacques Deguine, Ramnik J. Xavier, Jeffrey R. Moffitt, and Alan C. Mullen. Spatial transcriptomics of healthy and fibrotic human liver at single-cell resolution. Nature Communications, 16:319, 2025. doi: 10.1038/s41467-024-55325-4.

[21] Ian Mantel, Haoxuan Zhang, Juan Vargas, Gao Ce, Hope Townsend, Richard Bell, Amit Lakhanpal, Miriam R. Fein, Accelerating Medicines Partnership: RA/SLE Network, Thomas M. Norman, Dana E. Orange, Daniel C. Ramirez, Edward F. DiCarlo, Susan M. Goodman, Melanie H. Smith, Fan Zhang, Kevin Wei, Kushal K. Dey, Alexander Y. Rudensky, Christina S. Leslie, and Laura T. Donlin. SPP1*^hi^* macrophages in fibrin niches promote hyperplastic tissue remodeling in rheumatoid arthritis synovium. Science Translational Medicine, 18(855):eaed3087, 2026. doi: 10.1126/scitranslmed.aed3087.

[22] Bernhard Dumoulin et al. Spatial atlas of diabetic kidney disease reveals a B cell-rich subgroup. Nature, 653(8116):1158–1169, 2026. doi: 10.1038/s41586-026-10363-4. Data: Zenodo 19868428.

[23] Helmut Küchenhoff, Samuel M. Mwalili, and Emmanuel Lesaffre. A general method for dealing with misclassification in regression: the misclassification SIMEX. Biometrics, 62 (1):85–96, 2006. doi: 10.1111/j.1541-0420.2005.00396.x.

[24] Vlad Elgart and Joseph Loscalzo. Measuring and reducing segmentation-driven cell-type labeling errors in imaging spatial transcriptomics. bioRxiv preprint, posted with this manuscript, 2026.

[25] Daniel Borcard, Pierre Legendre, and Pierre Drapeau. Partialling out the spatial component of ecological variation. Ecology, 73(3):1045–1055, 1992. doi: 10.2307/1940179.

[26] Charles F. Manski. Identification of endogenous social effects: the reflection problem. The Review of Economic Studies, 60(3):531–542, 1993. doi: 10.2307/2298123.

[27] Sebastian Tiesmeyer, Niklas Müller-Bötticher, Alexander Malt, Leyao Ma, Sergio Marco Salas, Paul Kiessling, Paul Horn, Adrien Guillot, Louis B. Kuemmerle, Frank Tacke, Fabian J. Theis, Christoph Kuppe, Mats Nilsson, Roland Eils, Brian Long, and Naveed Ishaque. Identifying 3D signal overlaps in spatial transcriptomics data with ovrlpy. Nature Biotechnology, 2026. doi: 10.1038/s41587-026-03004-8.

[28] Dylan M. Cable, Evan Murray, Vignesh Shanmugam, Simon Zhang, Luli S. Zou, Michael Diao, Haiqi Chen, Evan Z. Macosko, Rafael A. Irizarry, and Fei Chen. Cell type-specific inference of differential expression in spatial transcriptomics. Nature Methods, 19:1076–1087, 2022. doi: 10.1038/s41592-022-01575-3.

[29] Ankit Agrawal, Stefan Thomann, Sukanya Basu, and Dominic Grün. NiCo identifies extrinsic drivers of cell state modulation by niche covariation analysis. Nature Communications, 15:10628, 2024. doi: 10.1038/s41467-024-54973-w.

[30] Shi-Tong Yang, Pai Peng, Hui-Feng He, Meng-Guo Wang, Bo-Han Si, Xiao-Fei Zhang, and Luonan Chen. Microenvironment-aware transcriptome reconstruction in spatial transcriptomics. Nature Communications, 17:10514, 2026. doi: 10.1038/s41467-026-77349-8.

[31] Pierre Boyeau, Stephen Bates, Can Ergen, Michael I. Jordan, and Nir Yosef. Mitigating bias in spatial transcriptomic pipelines via human feedback. *bioRxiv*, 2026. doi: 10.64898/2026.01.15.699786.

[32] Doron Haviv, Jan Remsik, Mohamed Gatie, Catherine Snopkowski, Meril Takizawa, Nathan Pereira, John Bashkin, Stevan Jovanovich, Tal Nawy, Ronan Chaligne, Adrienne Boire, Anna-Katerina Hadjantonakis, and Dana Pe’er. The covariance environment defines cellular niches for spatial inference. Nature Biotechnology, 43(2):269–280, 2025. doi: 10.1038/s41587-024-02193-4.

[33] Mingze Dong, David G. Su, Harriet Kluger, Rong Fan, and Yuval Kluger. SIMVI disentangles intrinsic and spatial-induced cellular states in spatial omics data. Nature Communications, 16:2990, 2025. doi: 10.1038/s41467-025-58089-7.

[34] Stathis Megas, Daniel G. Chen, Krzysztof Polanski, Hesam Asadollahzadeh, Moshe Eliasof, Carola-Bibiane Schönlieb, and Sarah A. Teichmann. Celcomen: spatial causal disentanglement for single-cell and tissue perturbation modeling. Nature Communications, 17:4126, 2026. doi: 10.1038/s41467-026-69856-5.

[35] J. R. Cook and L. A. Stefanski. Simulation-extrapolation estimation in parametric measurement error models. Journal of the American Statistical Association, 89(428):1314–1328, 1994. doi: 10.1080/01621459.1994.10476871.

[36] Raymond J. Carroll, David Ruppert, Leonard A. Stefanski, and Ciprian M. Crainiceanu. Measurement Error in Nonlinear Models: A Modern Perspective. Chapman and Hall/CRC, Boca Raton, FL, 2nd edition, 2006. doi: 10.1201/9781420010138.

[37] Jeffrey R. Moffitt. Spatial gene expression in the mouse colon during experimental colitis measured with MERFISH. Dataset, Dryad, 2024.

[38] Yixing Dong, Chiara Saglietti, Quentin Bayard, Almudena Espin Perez, Sabrina Carpentier, Daria Buszta, Stephanie Tissot, Rémy Dubois, Atanas Kamburov, Senbai Kang, Carla Haignere, Rita Sarkis, Sylvie Andre, Marina Alexandre Gaveta, Silvia Lopez Lastra, Nathalie Piazzon, Rita Santos, Katharina Loga, Caroline Hoffmann, George Coukos, Solange Peters, Vassili Soumelis, Eric Yves Durand, Laurence Leval, Raphael Gottardo, Krisztian Homicsko, and Elo Madissoon. Transcriptome analysis of archived tumors by Visium, GeoMx DSP, and Chromium reveals patient heterogeneity. Nature Communications, 16:4400, 2025. doi: 10.1038/s41467-025-59005-9.

[39] Anna Neufeld, Joshua Popp, Lucy L. Gao, Alexis Battle, and Daniela Witten. Negative binomial count splitting for single-cell RNA sequencing data. arXiv:2307.12985, 2023.

[40] Christine Yiwen Yeh, Min Woo Sun, Dixian Zhu, and Livnat Jerby. DR-GEM enables self-supervised machine learning for single-cell embeddings and annotations. Nature Communications, 17:11088, 2026. doi: 10.1038/s41467-026-77908-z.

[41] Vlad Elgart and Joseph Loscalzo. nichepeek: cell-type label error in spatial niche analyses. Python package, version 0.3.5, 2026. URL https://github.com/velgart/nichepeek. Zenodo, doi:10.5281/zenodo.23987423.

[42] Vlad Elgart and Joseph Loscalzo. claimbreak: segmentation-driven cell-type label error in imaging spatial transcriptomics. Python package, version 1.7.103, 2026. URL https://github.com/velgart/claimbreak. Zenodo, doi:10.5281/zenodo.23987641.

